# Interleukin-17A causes osteoarthritis-like transcriptional changes in human osteoarthritis-derived chondrocytes and synovial fibroblasts *in vitro*

**DOI:** 10.1101/2021.03.05.434099

**Authors:** Jolet Y. Mimpen, Mathew J. Baldwin, Adam P. Cribbs, Martin Philpott, Andrew J. Carr, Stephanie G. Dakin, Sarah J.B. Snelling

## Abstract

Increased interleukin (IL)-17A has been identified in joints affected by osteoarthritis (OA), but it is unclear how IL-17A, and its family members IL-17AF and IL-17F, can contribute to human OA pathophysiology. Therefore, we aimed to evaluate the gene expression and signalling pathway activation effects of the different IL-17 family members in fibroblasts derived from cartilage and synovium of patients with end-stage knee OA. Immunohistochemistry staining confirmed that IL-17 receptors A (IL-17RA) and IL-17RC are expressed in end-stage OA-derived cartilage and synovium. Chondrocytes and synovial fibroblasts derived from end-stage OA patients were treated with IL-17A, IL-17AF, or IL-17F, and gene expression was assessed with bulk RNA-Seq. Hallmark pathway analysis showed that IL-17 cytokines regulated several OA pathophysiology-related pathways including immune-, angiogenesis-, and complement-pathways in both chondrocytes and synovial fibroblasts derived from end-stage OA patients. While overall IL-17A induced the strongest transcriptional response, followed by IL-17AF and IL-17F, not all genes followed this pattern. Disease-Gene Network analysis revealed that IL-17A-related changes in gene expression in these cells are associated with experimental arthritis, knee arthritis, and musculoskeletal disease gene-sets. Western blot analysis confirmed that IL-17A significantly activates p38 and p65 NF-κB. Incubation of chondrocytes and synovial fibroblasts with IL-17A antibody secukinumab significantly inhibited IL-17A-induced gene expression. In conclusion, the association of IL-17-induced transcriptional changes with arthritic gene-sets supports a role for IL-17A in OA pathophysiology. Therefore, secukinumab could be investigated as a potential therapeutic option in OA patients.

## 1 Introduction

Osteoarthritis (OA) is the most common musculoskeletal disease, affecting 8.75 million people in the UK alone. Characterised by the loss of articular cartilage, it causes pain, disability, a reduced quality of life, and has a significant socioeconomic impact[1]. There are currently no disease-modifying treatments for OA and treatment is limited to joint replacement surgery with its associated costs and morbidity. OA is a multifactorial disease with a complex pathophysiology[1, 2]. Newly gained knowledge on the pathophysiology of osteoarthritis (OA) has shifted the traditional dogma of OA as a degenerative “wear-and-tear” disease of the articular cartilage to a hypothesis that OA is a whole joint disease with a significant inflammatory component.

Despite the loss of articular cartilage being hallmark feature of OA, the mechanisms underlying this OA-related cartilage degradation are poorly understood. Histological analyses have shown a clear infiltration of inflammatory cells into the synovium. Molecular interrogation has shown complement pathway activation in cartilage, synovium, and synovial fluid, and an increase in inflammatory mediators in synovium and synovial fluid[3–5]. Many of these mediators are hypothesized to be (over)produced by the resident stromal cells - chondrocytes in cartilage and synovial fibroblasts in synovium[5]. In addition, there is evidence for angiogenesis in OA cartilage, subchondral bone, synovium, and menisci[6, 7].

The IL-17 family of cytokines is increasingly identified as a contributor to OA pathogenesis. IL-17 is a family of 6 cytokines (IL-17A-F), from which homodimers IL-17A, IL-17F and their heterodimer IL-17AF are most studied[8–11]. IL-17A has been found in increased concentrations in serum and synovial fluid from OA patients compared to healthy controls, showing positive correlations with different pain, function, and disease severity scores[12–15]. When comparing inflamed with non-inflamed synovium from OA patients, an increased concentration of IL-17A was found in inflamed OA tissue, correlating with the release of IL-6, IL-23, and TGF-β1[16]. Another study identified a subgroup of patients with high IL-17A in their synovial fluid, alongside higher concentrations of inflammatory mediators (including IL-6, leptin, resistin, CCL7, and NGF), and reduced osteophytes, sclerosis, and minimum joint space width, thereby describing a potential inflammatory OA phenotype[17]. Genetic associations between polymorphisms in IL-17 genes and OA have been reported in different populations[18, 19]. Several animal models have studied the role of IL-17A and its receptors in (inflammatory) arthritis; injection of IL-17A induced the manifestations of OA in a rabbit model[20].

IL-17A, IL-17F, and IL-17AF all signal through the same heterometric receptor complex, which consists of IL-17 receptor A (IL-17RA) and IL-17RC[21, 22]. Both of these receptor subunits are essential for IL-17A/F signalling. While these cytokines seem to induce qualitatively similar signals, IL-17A homodimer produces a much more potent signal than the IL-17F homodimer in human primary foreskin fibroblasts and mouse embryonic fibroblasts, with the IL-17AF heterodimer producing an intermediate signal[11, 23, 24]. IL-17RA is expressed by nearly every cell type of the body with particularly high expression on immune cells, in contrast, IL-17RC is mostly expressed by non-immune cells, thereby mainly limiting IL-17 signalling to non-hematopoietic epithelial and stromal cells[22, 25]. However, IL-17-induced effects can differ between cell types, underlining the importance of studying its effects in each specific cell type or organ system[26, 27].

Although a growing number of clinical, animal, and genetic studies have identified a potential role for IL-17 in OA, the molecular mechanisms underpinning its role in OA pathophysiology is unknown. In cells from end-stage OA patients, IL-17A can increase the gene or protein expression of selected inflammatory mediators, including IL-6, IL-8, CXCL1, CCL2, COX2, and iNOS[28, 29]. In addition, IL-17A has been shown to affect ECM by increasing MMP production[30]. However, the individual effects of IL-17A, IL-17F, and IL-17AF in cells derived from OA patients throughout the whole transcriptome remain understudied. Therefore, this study aimed to identify and compare the changes in gene expression and activation of intracellular signalling pathways induced by IL-17A, IL-17F, and IL-17AF in chondrocytes and synovial fibroblasts derived from patients with end-stage knee OA. A better understanding of the similarities and differences of the effects of these three IL-17 cytokines in OA-derived primary cells will provide critical insight to their contribution to OA pathogenesis.

## 2 Materials and methods

### 2.1 Ethics approval

Ethical approval was granted for the Oxford Musculoskeletal Biobank (09/H0606/11) and (19/SC/0134) by the local research ethics committee (Oxford Research Ethics Committee B) for all work on human cartilage and human synovium, and informed consent was obtained from all patients according to the Declaration of Helsinki.

### 2.2 Immunohistochemistry

Cartilage and synovial tissue from end-stage OA patients was collected during total knee replacement surgery. Cartilage was dissected from the tibial plateau. Cartilage and synovial samples were immersed in 10% formalin for 0.5 mm/hour, embedded in paraffin before cutting 5μm sections and baking onto adhesive glass slides. Deparaffinisation and antigen retrieval procedure was performed using a PT Link machine (Dako, Glostrup, Demark) using FLEX TRS antigen retrieval fluid (Dako). Immunostaining was performed using an Autostainer Link 48 machine using the EnVision FLEX visualisation system (Dako) with anti-human IL-17RA, anti-human IL-17RC antibodies (R&D systems, Abingdon, UK) or universal negative control mouse (Dako) (Supplementary Table 1). Antibody binding was visualized by FLEX 3,3’-diaminobenzidine (DAB) substrate working solution and haematoxylin counterstain (Dako) following the protocols provided by the manufacturer. Antibodies were validated in-house to determine the concentration of antibody needed for positive staining with minimal artefact from the tissue. After staining, slides were dehydrated before mounting using Pertex mounting medium (Histolab, Gothenburg, Sweden). Negative controls are provided in Supplementary Fig. 1.

### 2.3 Isolation of primary OA chondrocytes and synovial fibroblasts for in vitro culture

Tissue from end-stage OA patients was collected during total knee replacement surgery. Cartilage was dissected from the tibial plateau and minced before overnight collagenase digestion in DMEM-F12 supplemented with 1% P/S and 1mg/ml collagenase IA (Sigma, supplied by Merck, Darmstadt, Germany). Synovium was minced before being collagenase digested for 2 hours. Collagenase-digested tissue was filtered through a 70μm cell strainer, and cells were plated out in 10-cm dishes in D10 (DMEM-F12 (Gibco, supplied by Fisher Scientific, Loughborough, UK) with 10% Foetal Bovine Serum (FBS) (Labtech International, Heathfield, UK) and 1% Penicillin/Streptomycin (P/S) (Gibco)). Once 95% confluent, chondrocytes were cryopreserved, while synovial fibroblasts were passaged once (p=1) and then cryopreserved.

### 2.4 Cell culture for RNA Sequencing

Chondrocytes (n=6 patients) were expanded to p=2 and synovial fibroblasts (n=6 patients) to p=3 in D10 after which they were plated out in 12-well plates. Cells were left to attach for 24 hours, before medium was changed to serum-free DMEM-F12 with P/S (D0) and left for 24 hours. On the day of the experiments, vehicle control, and 10ng/ml IL-17A, IL-17F, or IL-17AF (all from Biolegend UK Ltd, London, UK) was made up in D0. Old D0 was removed and D0 with vehicle control or IL-17 was added. After 24 hours, treatment medium was removed and cells were washed with PBS, before being harvested in Trizol (Invitrogen, supplied by ThermoFisher Scientific, Waltham, MA, USA), and stored at −80°C.

### 2.5 Bulk RNA Sequencing

RNA was extracted using a Direct-zol MicroPrep kit with DNase treatment (Zymo Research, Irvine, CA, USA) following the manufacturer’s instructions. RNA concentration was measured using a NanoDrop spectrophotometer. RNA quality of eight randomly chosen samples were assessed using High Sensitivity RNA ScreenTapes (Agilent) on an Agilent 2200 TapeStation. Library preparation was done using a NEBNext Ultra II Directional RNA Library Prep Kit for Illumina with poly-A selection (Illumina, San Diego, CA, USA) following the manufacturer’s instructions. Every library was quantified for DNA content with High Sensitivity DNA ScreenTapes (Agilent). Libraries from 24 samples with unique identifiers were pooled and run on an Illumina NextSeq 500 using the 75 cycles NextSeq High Output kit (Illumina).

Raw FASTQ files containing reads were generated by the Illumina software CASAVA v1.8. The raw FASTQ files were processed using CGAT-flow readqc and mapping workflows (https://github.com/cgat-developers/cgat-flow)[31]. The quality of the reads was assessed using FASTQC and ReadQC. Raw reads were aligned to the GRCh38 reference genome using HiSat2 (v2.0.5). The mapped reads were visualised using IGV (v2.3.74) to further assess quality of mapping. The quantification of mapped reads against GCRh38 reference genome annotation was carried out using FeatureCounts (v1.5.0). Downstream analyses were performed using R version 3.5.1 (R Foundation, Vienna, Austria), and RStudio version 1.1.456 (RStudio, Boston, MA, USA). Differential expression analysis was performed using the DESeq2 package[32] using ‘apeglm’ method to apply the shrinkage of logarithmic fold change[33]. The adjusted p-value (padj) and significance of changes in gene expression were determined by applying the Bonferroni-Hochberg correction of 5% false discovery rate. PCA plots were generated using the package ggplot2[34], heatmaps were generated using the package pheatmap (PCA and heatmaps in Supplementary Fig. 2 and 3), and EnhancedVolcano was used to create volcano plots. Gene-set enrichment analysis for hallmark pathways were performed using clusterProfiler on genes that were significantly changed compared to control (padj < 0.05) ranked by log2 fold change (LFC)[35]. Disease-gene network (DisGeNET) analysis was performed using the DOSE package on genes padj<0.05 and LFC±1[36].

### 2.6 Cell culture for western blot

Chondrocytes (n=3) and synovial fibroblasts (n=3) were expanded, seeded in 12-well plates as described before. On the day of the experiments, vehicle control, IL-17A, IL-17F, and IL-17AF were diluted in D0 medium to 10ng/ml. Old D0 medium was removed and stimulation medium was added. Cells were harvested for western blotting 0 mins, 10 mins, 30 mins, 1 hour and 8 hours after treatment commenced as described previously[37], frozen, and stored at −20°C until analysis.

### 2.7 SDS-PAGE and Western blot

Cell lysates were mixed with 2x Laemmli Sample Buffer (Bio-Rad) in a 1-to-1 ratio and separated by gel electrophoresis in a 10% Mini-PROTEAN TGX Precast gels (Bio-Rad Watford, UK). After gel electrophoresis, proteins were blotted on PVDF membrane (Bio-Rad) using a Trans-Blot Turbo Transfer System (Bio-Rad). Membranes were blocked in blocking buffer (10% milk powder and 2% BSA in TBS-T) and subsequently stained with loading control (vinculin) and phosphorylated-protein antibody overnight in antibody buffer (5% BSA and 1% Tween-20 in TBS-T) (Supplementary Table 2). The blots were washed in TBS-T and incubated in secondary antibody in antibody buffer for 2 hours before washing again in TBS-T before visualising with ECL (GE Healthcare, Chicago, IL, USA) and imaging in an ALLIANCE 6.7 Chemiluminescence Imaging System (UVITEC, Cambridge, UK). The membranes were subsequently stripped (Takara BioInc, Kusatsu, Japan), blocked, and stained with total-protein antibody in antibody buffer overnight. Blots were then washed, stained with secondary antibody, washed again, and imaged. ImageJ was used to analyse the intensity of the stained bands for semi-quantitative analysis using the 0h control for each patient as a reference. Supplementary Fig. 4 shows representative western blots for each antibody.

### 2.8 Cell culture for inhibitor treatment and cells-to-cDNA synthesis

Chondrocytes and synovial fibroblasts were cultured and plated out in 96-well plates as described earlier. Secukinumab with 10ng/ml IL-17A or vehicle control was added to the cells. After 24 hours, cells were washed with PBS before being harvested in cells-to-cDNA Cell Lysis Buffer (Ambion Inc, Foster City, CA, USA), and transferred to a PCR plate for cells-to-cDNA synthesis. cDNA was prepared using a cells-to-cDNA kit following the manufacturer’s instructions (Ambion).

### 2.9 Real time quantitative PCR

Real-time quantitative polymerase chain reaction (RT-qPCR) were carried out in a 10μL volume in 384-well plates using Fast SYBR Green Master Mix (Applied Biosystems, Foster City, California, USA). Primers (Supplementary Table 3) were purchased from Primerdesign Ltd (Primerdesign Ltd, Eastleigh, UK). All RT-qPCRs were performed using a ViiA7 (Life Technologies, Paisley, UK), which included 40 cycles and a melt-curve. Samples were analysed against two reference genes, glyceraldehyde 3-phosphate dehydrogenase (*GAPDH*) and β-actin (*ACTB*) (Primerdesign Ltd), using the delta-CT or delta-delta Ct method[38].

### 2.10 Statistical analysis

Statistical analyses for bulk RNA-Seq were performed using R and RStudio. All other statistical analyses were performed using in GraphPad Prism 8.1.2 (GraphPad Software, La Jolla, CA, USA). For treatment with IL-17 for western blot analysis, Friedman test with Dunn’s multiple comparisons test was used and data is shown as mean±SD. For all experiments with IL-17A and inhibitors, differences between treatment with IL-17A alone and other treatments Friedman test with Dunn’s test to correct for multiple comparisons, except when there was missing data in which Kruskal-Wallis test with Dunn’s multiple comparisons test was used. All data for inhibitor experiments is shown as the individual measurements with mean. * = p<0.05, ** = p<0.01, *** = p<0.001, **** = p<0.0001.

## 3 Results

### 3.1 IL-17 receptors are expressed by end-stage osteoarthritis chondrocytes and synovial fibroblasts

Cartilage and synovium from end-stage OA patients undergoing total knee replacement surgery were either formalin-fixed or digested to isolate and expand resident fibroblasts (chondrocytes and synovial fibroblasts respectively). To validate that these fibroblasts can respond to IL-17 cytokines, mRNA expression of their receptors *IL17RA* and *IL17RC* was assessed by RT-qPCR (Fig. 1A). Both genes were expressed by chondrocytes and synovial fibroblasts, although *IL17RA* was more highly expressed than *IL17RC* with a difference of 2.03 CT (n=5, 0.84-3.18) in chondrocytes and 3.39 CT (n=4, 2.79-4.20) in synovial fibroblasts. Protein expression of IL-17RA and IL-17RC was confirmed in end-stage OA cartilage and synovial tissue (Fig. 1B).

**Figure 1.**
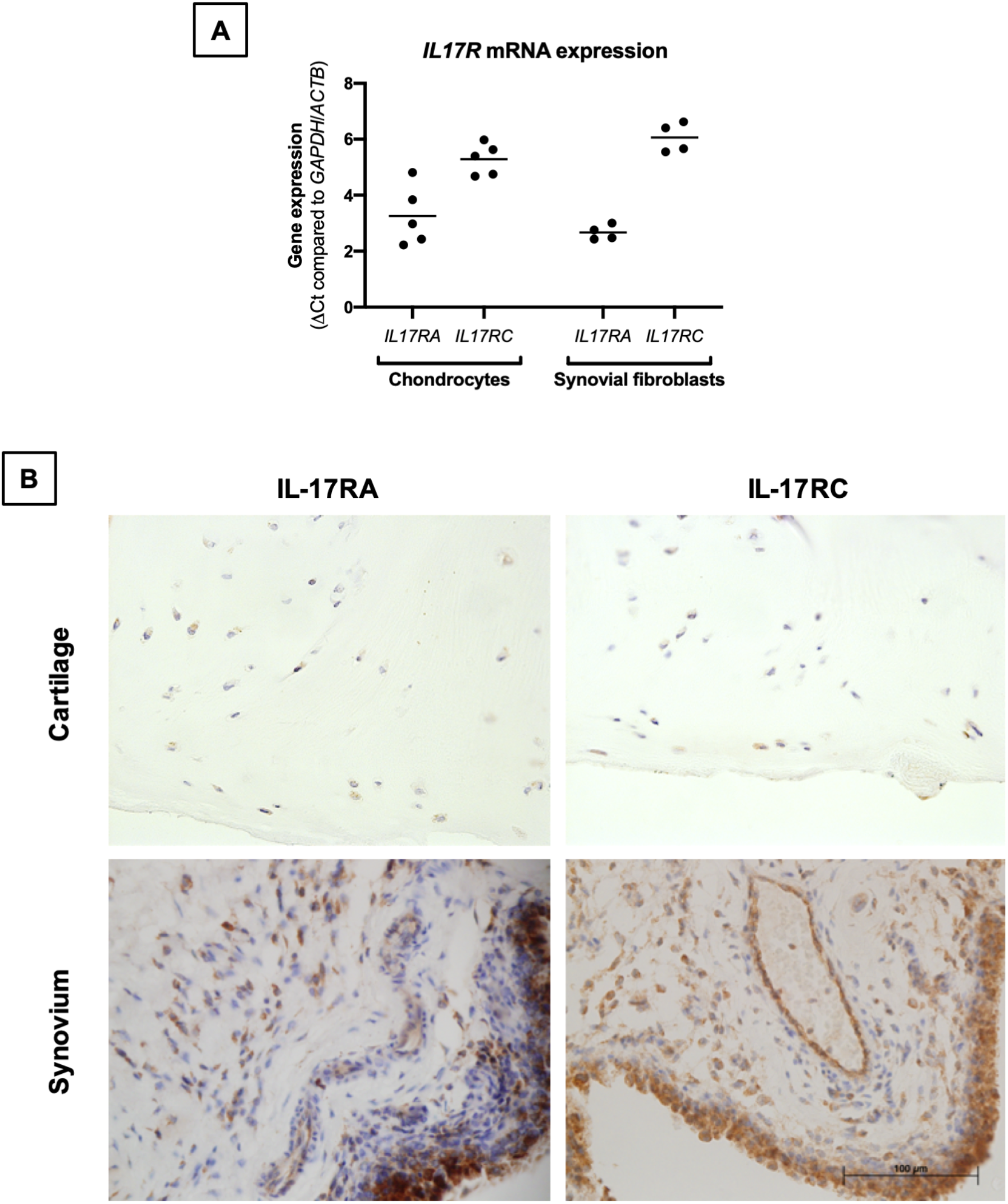
Expression of IL-17 receptor A (IL-17RA) and IL-17RC in end-stage OA. (A) mRNA expression of *IL17RA* and *IL17RC* in primary cultured chondrocytes (n=5) and synovial fibroblasts (n=4). Gene expression was calculated using the dCT-method using both *GAPDH* and *ACTB* as housekeeper. (B) Protein expression of IL-17RA and IL-17RC in end-stage OA cartilage and synovium. Antibodies were visualised with DAB (brown) and counterstained with haematoxylin (blue). Images were taken at 40x magnification.

### 3.2 IL-17A induces the most potent and greatest changes in gene expression

The transcriptome of sets of samples (24 samples of chondrocytes and 24 samples of synovial fibroblasts representing n=6 patients, treated for 24 hours with vehicle control, or IL-17A, IL-17F, or IL-17AF, all at 10 ng/ml) were analysed using poly-A tail selected, mRNA sequencing. After normalisation and accounting for donor variability, log-fold2 (LFC) changes were calculated with the adjusted p<0.05. Treatment with IL-17 cytokines significantly changed a range of genes compared to control (Fig. 2). The number of changed genes that were changed at least LFC±1 varied by IL-17 treatment type, with the highest number changed by IL-17A (856 genes for chondrocytes, 330 for synovial fibroblasts), followed by IL-17AF (188 genes/55 genes), and finally IL-17F (39 genes/17 genes) (full list in Supplementary Tables 4 and 5; all volcano plots in Supplementary Fig. 5 and 6). The most significantly changed gene in both cell types by all three IL-17 cytokines was *NFKBIZ*, a gene that encodes for the protein “Inhibitor of NF-κB Zeta” (IκBζ), a central regulator of IL-17 signalling. The three most highly regulated genes by IL-17A were *CCL20*, *IL6*, and *NOS2* in chondrocytes, and *CSF3*, *CXCL1*, and *CCL20* in synovial fibroblasts, which are known IL-17-induced genes.

**Figure 2.**
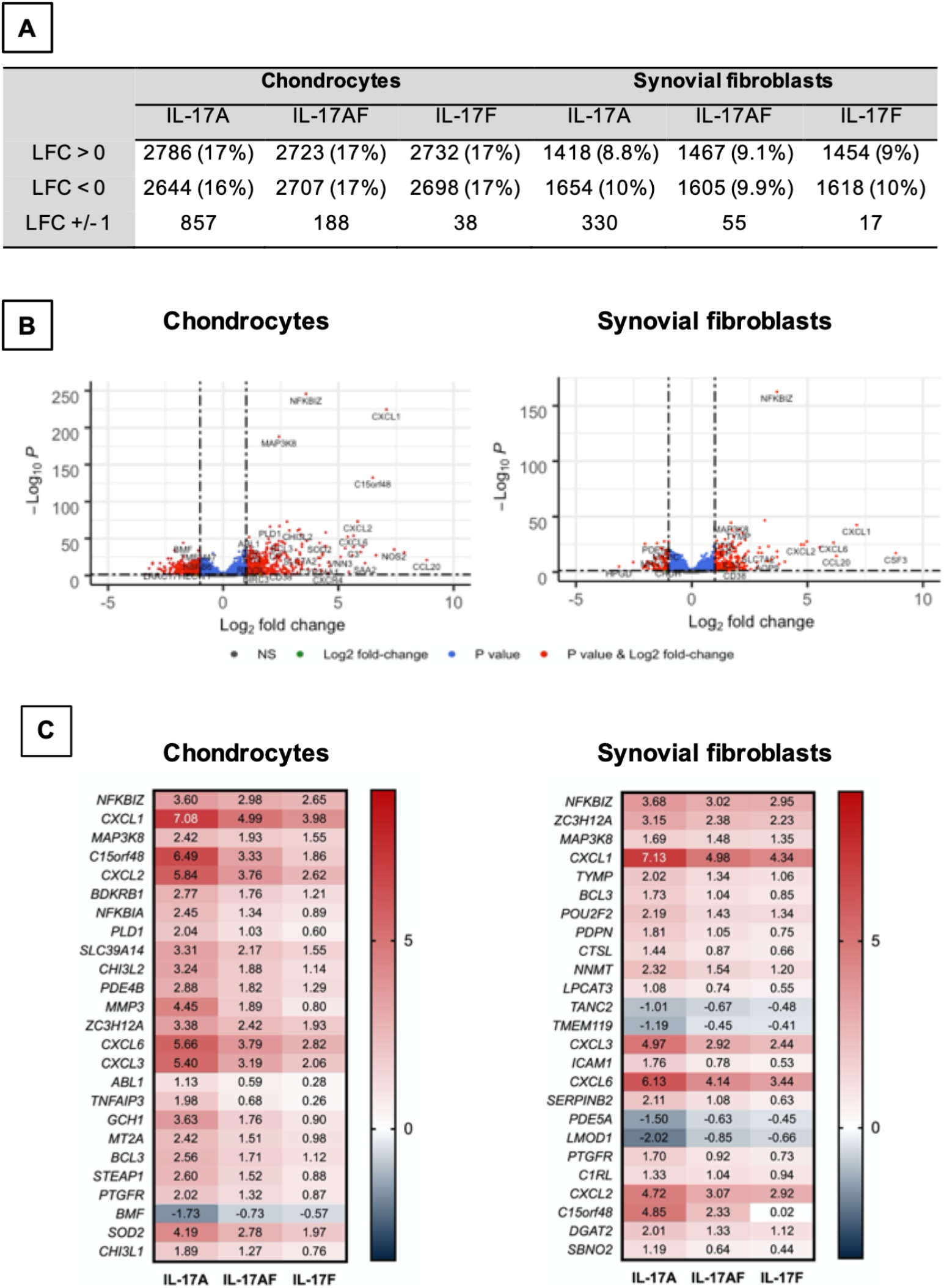
Changes in gene expression after treatment with IL-17A, IL-17AF, or IL-17F compared to vehicle control measured by bulk RNA-Seq. (A) Overview of changes induced by IL-17 cytokines (padj<0.05). (B) Volcano plots showing the changes in gene expression after treatment with IL-17A in chondrocytes (left) and synovial fibroblasts (right). **Grey** = not significant (NS); **green** = Log2 fold change of at least 1 or −1, but p>0.05; **blue** = p-value < 0.05, but log2 fold change >-1 and <1; **red** = p-value<0.05 and log2 fold change of at least −1 or +1. (C) Heatmap of the 25 most significantly differentially expressed genes by chondrocytes (left) and synovial fibroblasts (right). **Red** = upregulated, **blue** = downregulated. n=6

Overall, IL-17A induced the strongest transcriptional response, followed by IL-17AF, and finally IL-17F. For 81% of these genes in chondrocytes and 91% in synovial fibroblasts, the differences in transcriptional response between IL-17AF and IL-17A were larger than between IL-17AF and IL-17F. Therefore, IL-17AF-induced gene expression changes more closely resemble IL-17F-induced effects than IL-17A-induced effects. However, not all genes followed this pattern. For example, while IL-17A, IL-17AF, and IL-17F all induced a similar increase in *NFKBIZ* in synovial fibroblasts (padj=2.09E-163, LFC of 3.68, 3.03, and 2.95 respectively), only IL-17A induced the expression of *CCL20* (padj=3.46E-15, LFC of 6.25, 0.07, and 0.02, respectively), and only IL-17A and IL-17AF were induced the expression of *CSF3* (padj=8.69E-18, LFC of 8.82, 5.97, and 0.02, respectively). In addition, some genes were only expressed and strongly increased in either chondrocytes (*NOS2*, padj=7.49E-36, LFC 7.42, 5.15, and 4.31, respectively) or synovial fibroblasts (*CSF3* as described above), showing that IL-17 cytokine culture can induce transcriptional responses that are cell type dependent.

### 3.3 IL-17A treatment causes changes in inflammation-related biological pathways

To elucidate the biological processes that are affected by IL-17 in chondrocytes and synovial fibroblasts, the dataset was analysed using the Gene Set Enrichment Analysis (GSEA) hallmark gene set. The GSEA is a collection of 50 hallmark gene pathways which condense information from over 4,000 original overlapping gene sets from specific collections[39]. In chondrocytes, 21, 24, and 21 pathways were significantly changed by treatment with IL-17A, IL-17AF, and IL-17F, respectively. In synovial fibroblasts, IL-17 cytokines caused significant changes in 20, 14, and 11 pathways after treatment with IL-17A, IL-17F, and IL-17AF, respectively. Generally, the largest changes were made by IL-17A, followed by IL-17AF, and finally IL-17F (Fig. 3A-B, Supplementary Table 6). Pathways including inflammatory responses, complement, hypoxia, angiogenesis, and glycolysis were changed in both chondrocytes and synovial fibroblasts and have been linked to OA pathophysiology.

**Figure 3.**
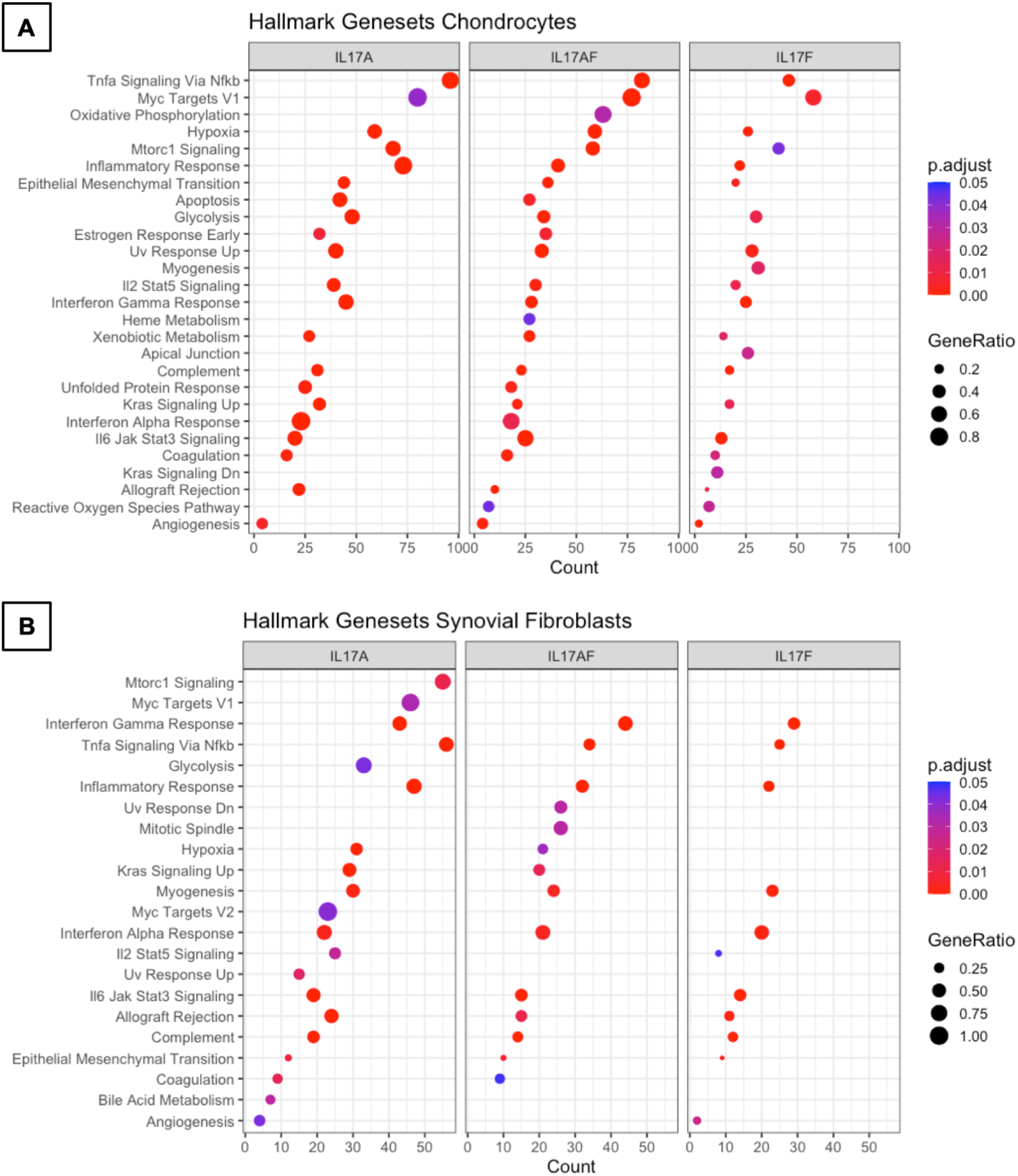
Results from GSEA Hallmark pathway analysis. Dot plots representing the changes in GSEA hallmark pathways after IL-17A, IL-17AF, and IL-17F treatment in (A) chondrocytes and (B) synovial fibroblasts, including the gene ratio (ratio of DEG over the total number of genes in pathway), gene count (number of DEG in pathway), and the adjusted p-value.

### 3.4 IL-17A induced changes are associated with experimental arthritis

Disease-gene network (DisGeNET) analysis was used to study the gene-disease associations of the IL-17-induced changes in gene expression (padj<0.05, LFC>±1). The 25 most significantly associated diseases after IL-17A treatment in chondrocytes included juvenile arthritis, experimental arthritis, musculoskeletal diseases, anoxia, and hyperalgesia (Fig. 4A) In synovial fibroblasts the 25 most significantly associated diseases after IL-17A treatment included experimental arthritis, musculoskeletal diseases, and pain, but mostly included IL-17 associated diseases including lung disease, viral bronchiolitis, and respiratory syncytial virus infections (Fig. 4B). Although the 25 most significantly associated disease after IL-17AF- and IL-17F treatment were similar to those induced by IL-17A, the gene counts were notably lower than the changes associated with IL-17A.

**Figure 4.**
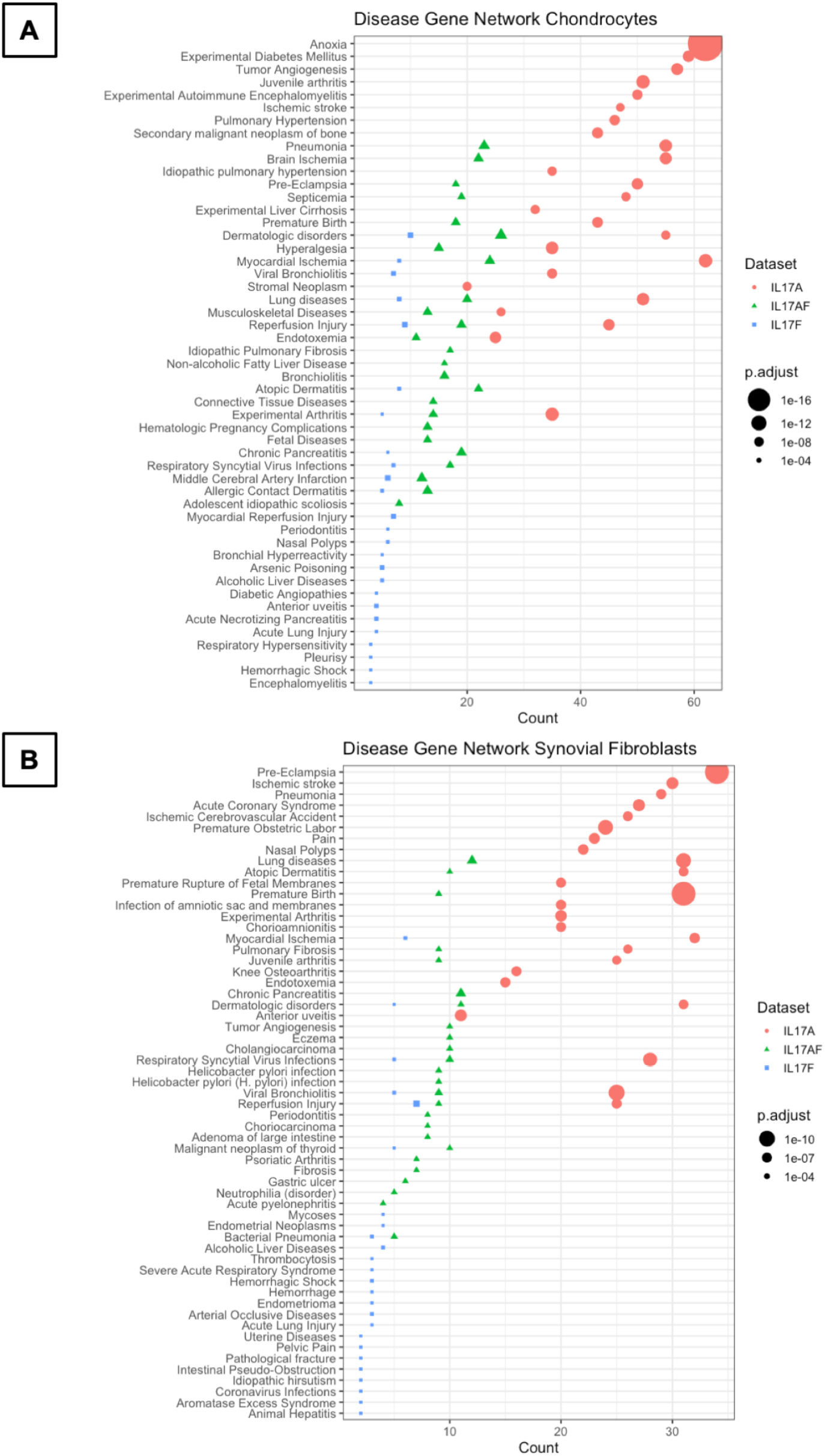
Results from Disease-Gene Network (DiGeNeT) in (A) chondrocytes and (B) synovial fibroblasts. Dotplots displaying 25 most significantly associated disease gene sets with the differently expressed genes (DEGs) (padj<0.05, LFC±1) after treatment with IL-17A (**coral**), IL-17AF (**green**), and IL-17F (**blue**), including gene count (number of DEG in each disease geneset) and the adjusted p-value.

### 3.5 IL-17A activates p38 and NF-kB pathways

To better understand the similarities and differences between the signalling induced by IL-17 cytokines, chondrocytes or synovial fibroblasts from end-stage OA patients were treated with control medium or 10ng/ml IL-17 for 10 minutes (Fig. 5), 30 mins, 1 hour, or 8 hours (Supplementary Fig. 7). Activation of the ERK1/2 (p44/p42 MAP kinases), p38 and p65 NF-κB intracellular cell signalling pathways was assessed by western blot analysis. Semi-quantitative analysis showed that IL-17A induced the strongest activation in p38 and NF-κB, followed by IL-17AF, with IL-17F causing only subtle activation. In contrast, ERK1/2 was most strongly activated by IL-17AF, followed by IL-17A and IL-17F which both caused no/minor increases.

**Figure 5.**
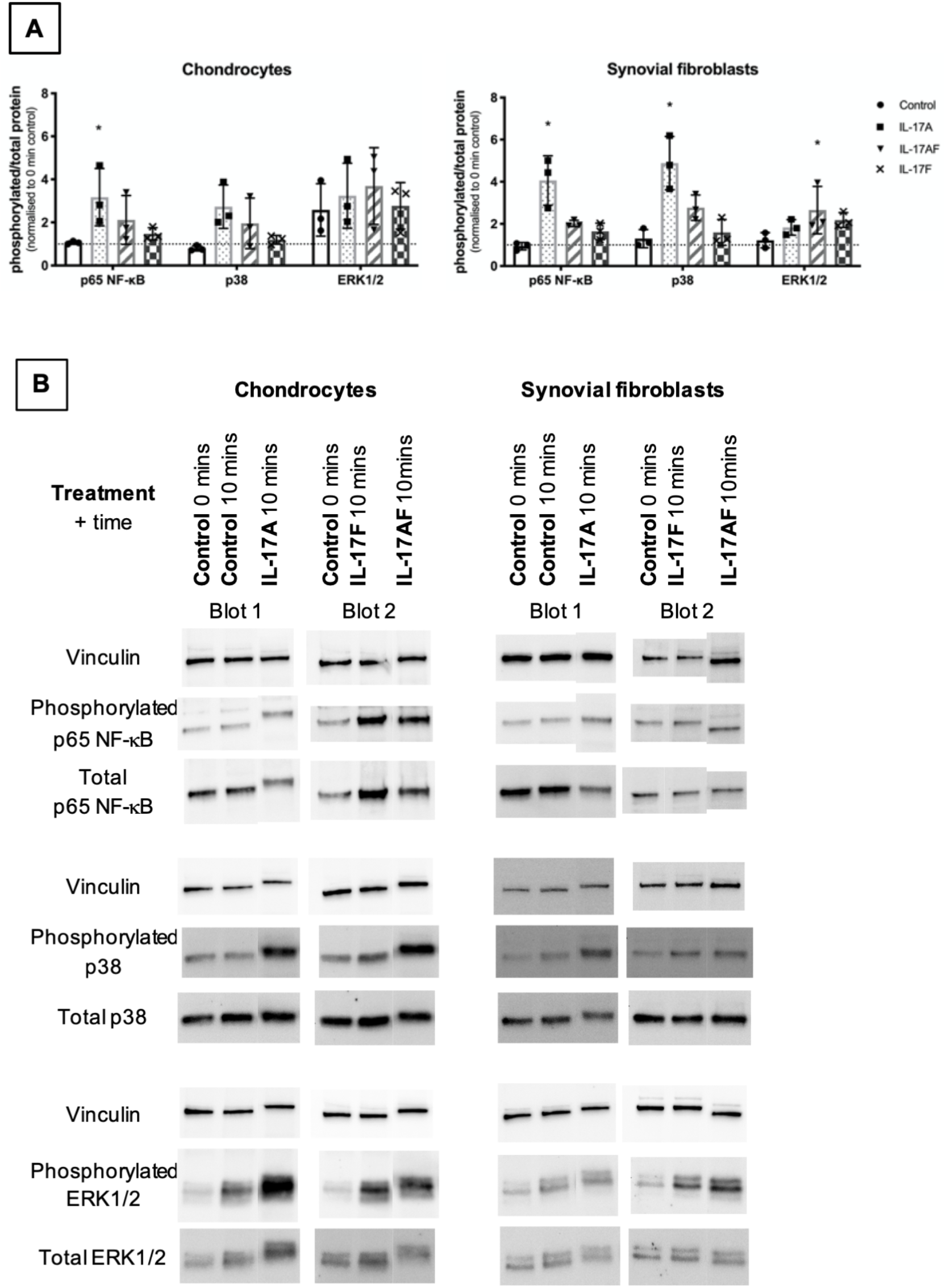
Activation of p65 NF-κB, p38 MAP kinase, and ERK1/2 (p44/p42 MAP kinases) in chondrocytes and synovial fibroblasts by IL-17A, IL-17AF, and IL-17F (all 10 ng/ml) after 10 minutes stimulation. (A) Relative ratio of phosphorylated protein over total protein compared to baseline control (0 mins). Friedman test for each intracellular protein with Dunn’s multiple comparisons test. Mean±SD. N=3. * = p<0.05, ** = p<0.01, *** = p<0.001, **** = p<0.0001. (B) Representative western blots for vinculin (loading control), phosphorylated, and total protein of each intracellular signalling protein.

### 3.6 Secukinumab inhibits IL-17A-induced expression of OA-relevant genes

The clinically-used IL-17A-antibody secukinumab was used to confirm that IL-17A drives the changes seen in expression of OA-relevant genes related to inflammation, matrix turnover, fibroblast activation, and intracellular signalling. Chondrocytes and synovial fibroblasts from end-stage OA patients were treated with 10ng/ml IL-17A for 24 hours without or with increasing, clinically-relevant concentrations of secukinumab (0.5, 5.0, and 50 μg/ml), and changes in expression of 8 genes that were found to have been significantly changed by IL-17A in the RNA-Seq experiment were analysed with RT-qPCR. Secukinumab decreased the IL-17A-induced increase in gene expression for every gene tested in a dose-related manner (Fig. 6). In chondrocytes, 5 μg/ml secukinumab caused a statistically significant inhibition of IL-17A induced *MMP1* and *MAP3K8* mRNA expression, and 50 μg/ml secukinumab was able to significantly inhibit IL-17A-induced gene expression for all genes tested. In synovial fibroblasts, 5 μg/ml secukinumab caused a statistically significant decrease in IL-17A-induced *IL6*, *PDPN*, and *NFKBIZ* mRNA expression, and 50 μg/ml secukinumab caused a statistically significant decrease in the IL-17A-induced gene expression of all genes tested except for *PDPN*. Inhibition of p38 or NF-κB p65 in end-stage OA chondrocytes or synovial fibroblasts did not significantly decrease IL-17A-induces genes in gene expression. These findings suggest that there is redundancy in IL-17A-induced intracellular signalling (Supplementary Fig. 8 and 9).

**Figure 6.**
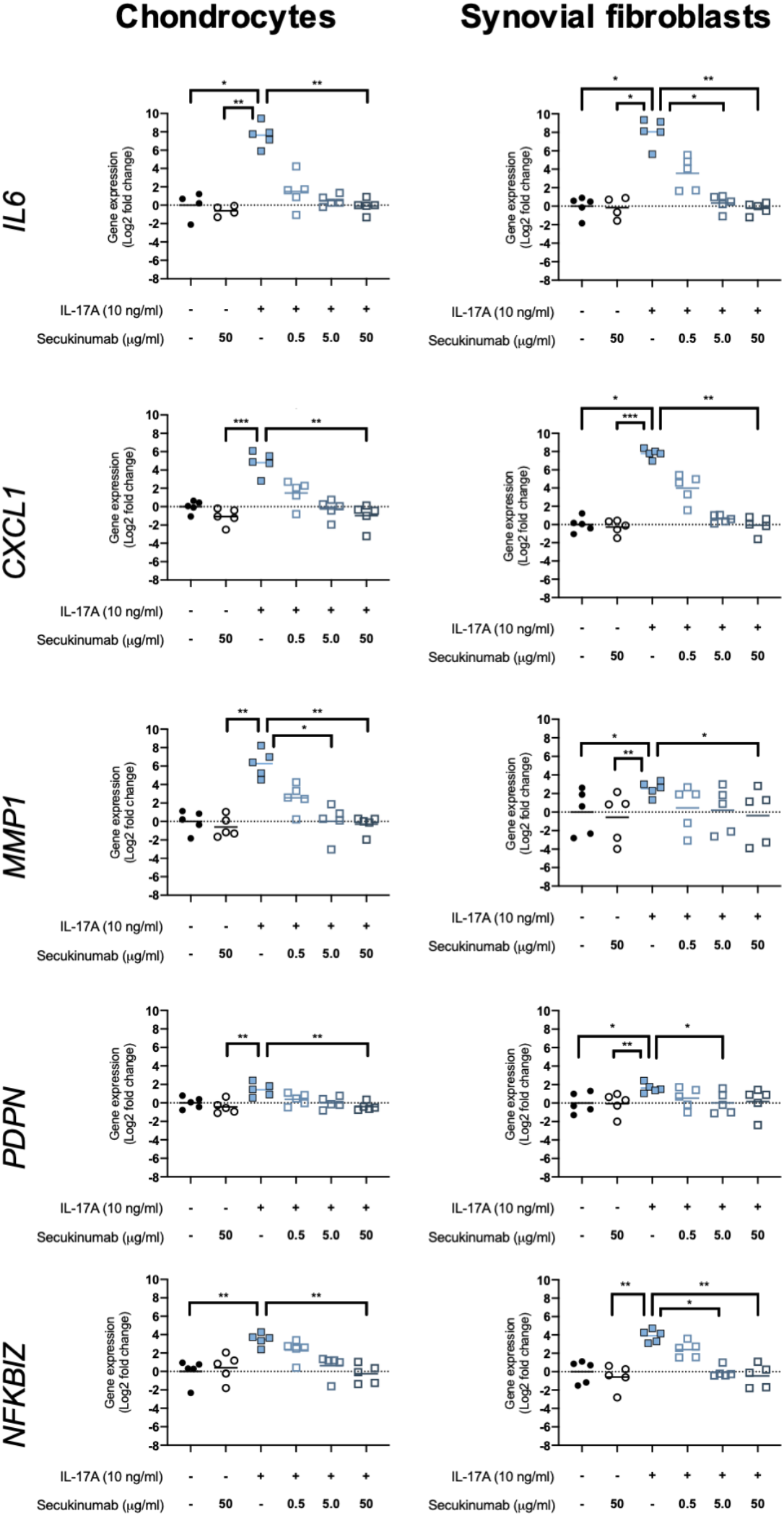
Gene expression of chondrocytes and synovial fibroblasts after treatment with IL-17A (10 ng/ml) with or without clinically-used IL-17A-antibody secukinumab in different concentrations (0.5, 5.0, or 50 μg/ml). Gene expression was calculated using the ddCT-method using both *GAPDH* and *ACTB* as housekeepers. Gene expression is expressed as log2 fold change compared to control. Changes in gene expression were calculated comparing each treatment to IL-17A treatment alone. Kruskal-Wallis test with Dunn’s multiple comparisons test (*IL6*). Friedman test with Dunn’s multiple comparisons test (all other genes). Individual values and mean. N=5. * = p<0.05, ** = p<0.01, *** = p<0.001, **** = p<0.0001.

## 4 Discussion

There is a growing number of reports in clinical, animal, and genetic studies that support a role for IL-17 cytokines (including IL-17A, IL-17AF, and IL-17F) in OA. However, the specific effect IL-17 cytokines could have in OA disease pathophysiology remains unclear. Therefore, the overarching aim of this study was to investigate and compare the effect of IL-17A, IL-17AF, and IL-17F on the transcriptome and activation of intracellular signalling pathways in end-stage OA derived fibroblasts. The most active IL-17 cytokine identified, IL-17A, was then inhibited to assess its potential as a therapeutic option in OA.

IL-17 cytokines induced changes in expression of diverse set of genes in both chondrocytes and synovial fibroblasts derived from end-stage OA patients. IL-17 targeted genes included those encoding signalling molecules, cytokines, and chemokines. Overall, IL-17A caused the strongest transcriptional response, followed by IL-17AF, and finally IL-17F. For most genes, IL-17AF-induced strength of response was closer to that of IL-17F than IL-17A, suggesting that the IL-17F-subunit strongly reduces its potential to induce transcriptional changes. However, the fact that not all genes followed this pattern underlines the complexity of IL-17-signalling and the likeliness that these cytokines have different roles[27, 40]. Therefore, further work should investigate the similarities and differences of the effects of these three cytokines and their potential role in the OA pathophysiology.

Pathway analysis of the IL-17-induced genes revealed regulation of multiple cellular pathways including those related to immune responses, complement, and angiogenesis. Many of these pathways have been implicated in the development and/or progression of OA. DisGeNet analysis showed that especially IL-17A-induced changes in the transcriptome of chondrocytes and synovial fibroblasts are associated with juvenile arthritis, experimental arthritis, and musculoskeletal diseases, supporting that IL-17A could play an important role in OA. While IL-17AF- and IL-17F-induced transcriptional changes were also associated with these diseases, these associations the transcriptional changes were weaker for IL-17AF and IL-17F compared to IL-17A. Therefore, if these cytokines occur at similar concentrations in the joint, IL-17A is the most likely cytokine of this family to play a role in OA pathophysiology.

Western blot analysis of the activation of intracellular signalling proteins was used to better understand the differences in IL-17-induced gene expression between IL-17A, IL-17AF, and IL-17F. The intracellular signalling proteins p38 and NF-κB p65 were activated by IL-17 cytokines, the extent of the activation mirrored the number of genes transcriptionally regulated by each of these cytokines in chondrocytes and synovial fibroblasts: IL-17A causing a relatively strong activation, IL-17AF causing a modest activation, and IL-17F causing a small activation. As the activation of NF-κB and p38 have led to increases inflammation in other types of fibroblasts[41, 42], these results are consistent with the IL-17-induced expression of immune-related genes seen in the RNA-Seq data. IL-17AF caused the activation of ERK1/2, with both IL-17A and IL-17F causing no/minor activation. In IL-17 signalling in ST2 cells and mouse fibroblasts, ERK1/2 is not only able to induce transcription of IL-17A/F target genes, but it can also contribute to the negatively regulation of IL-17 signalling by phosphorylating C/EBPβ which ultimately leads to the deubiquitylation of TRAF6, a protein that when ubiquitinylated mediates MAPK and NF-κB signalling[43–46]. Therefore, IL-17AF-induced activation of ERK1/2 in this study suggests there is a higher chance of ERK-dependent phosphorylation of C/EBPβ, leading to a stronger dampening of IL-17-induced signalling. However, better insight in the differential signalling of the three IL-17 cytokines in different cell types is needed to get a better understanding of their potential functions, which can be used to further unravel their potential role in promoting inflammation, influx of immune cells, and matrix destruction in OA.

As IL-17A is already a clinical target in other inflammatory (musculoskeletal) diseases, inhibition of IL-17A-induced gene expression was investigated. Inhibition of IL-17A-induced gene expression changes by the IL-17A antibody secukinumab confirmed that the expression of these genes is increased by IL-17A. As several clinical studies have demonstrated that OA patients have increased concentrations of IL-17A in their synovial fluid[12–17], our results suggest that secukinumab could be a potential therapeutic option for those OA patients with high concentrations of IL-17A in the OA joint.

The limitations of this study include the use only one concentration of IL-17 cytokines at only one time point for the gene expression studies. This concentration was selected based on dose-response curves but was necessary due to the limited availability of primary fibroblasts from OA patients. Future studies should study a range of concentrations over time to study the concentration- and time dependent effects of IL-17, which could especially be important as OA is a chronic disease that likely develops over several years or even decades. Lastly, expression changes in RNA-Seq do not necessarily correlate to changes in protein expression. Therefore, it is crucial that future studies confirm that the IL-17-induced changes in gene expression that were seen in this study are translated into changes in protein expression and functional activity of hallmark pathways. This all will give more insight in the potential benefit of inhibitors of these IL-17 cytokines for OA patients with increased concentrations of IL-17 in OA joints.

Future studies should determine the (relative) concentrations of IL-17 cytokines in the OA joint to determine which of these cytokines is likely to be most important in this disease. In addition, not only chondrocytes and synovial fibroblasts, but also fibroblasts and immune cells from other key joint tissues including meniscus and fat pad, should be used in isolation and in combination to look at the effect these cytokines could have across the joint organ.

In conclusion, this study shows that IL-17A, and to a lower extent IL-17AF and IL-17F, induced changes in many genes that are linked to several OA pathophysiology-related pathways. While overall IL-17A caused the strongest transcriptional response, followed by IL-17AF and IL-17F, not all genes followed this pattern. The chronic, low-grade inflammation seen in OA fits well with the effects IL-17A on immune-related pathways. Disease-gene network analysis showed that IL-17A-induced changes in chondrocytes and synovial fibroblasts are associated with experimental arthritis, knee osteoarthritis, and musculoskeletal disease, which further highlights the potential importance of this cytokine in OA. The clinically-used IL-17A antibody secukinumab was able to significantly inhibit IL-17A-induced gene expression, confirming that IL-17A is responsible for these changes in gene expression. As there are currently no disease-modifying OA drugs, secukinumab should be investigated as a potential therapeutic option in (a subgroup of) patients with OA.

## Author contributions

Jolet Y. Mimpen, Sarah J.B. Snelling, Mathew J. Baldwin, Stephanie G. Dakin, Adam P. Cribbs, and Andrew J. Carr contributed to the study conception and design. Material preparation, data collection and analysis were performed by Jolet Y. Mimpen, Mathew J. Baldwin, Adam P. Cribbs, and Martin Philpott. The first draft of the manuscript was written by Jolet Y. Mimpen and all authors commented on previous versions of the manuscript. All authors read and approved the final manuscript.

## Acknowledgements

The authors would like to thank Dr Ash Maroof (UCB) for providing the secukinumab for the inhibitor experiments. In addition, the authors would like to thank the Oxford Musculoskeletal Biobank, our research assistant Louise Appleton, research nurses Debra Beazley, Bridget Watkins, and Kim Wheway, and the knee surgeons Professor Andrew Price, Mr Nick Bottomley, Mr Will Jackson, and Mr Abtin Alvand for their invaluable help in obtaining the human tissue used for this study.

## Availability of data and material

The dataset generated from the bulk RNA Sequencing experiment will be made available through the NCBI Gene Expression Omnibus, https://www.ncbi.nlm.nih.gov/geo/.

