## Supplementary Material for "Interleukin-17A causes osteoarthritis-like transcriptional changes in human osteoarthritis-derived chondrocytes and synovial fibroblasts *in vitro*"

### 1 Supplementary Methods

#### 1.1 Cell culture for p38 and NF-κB inhibitor treatment (regarding Supplementary Figures 8 and 9)

Chondrocytes and synovial fibroblasts were cultured and plated out in 96-well plates as described earlier. On the day of the experiment, cells were pre-treated for 2 hours with JSH-23, SB204580, or vehicle control (Table 4-4) before the addition of 10 ng/mL concentration of IL-17A for 24 hours. SB203580 inhibits p38 signalling by suppressing the activation of MAPKAP kinase-2 and inhibiting the phosphorylation of heat shock protein 27. JSH-23 inhibits NF-κB signalling by interfering with the nuclear localisation signals of p65. After 24 hours, cells were washed with PBS before being harvested in cells-to-cDNA Cell Lysis Buffer (Ambion Inc, Foster City, CA, USA), and transferred to a PCR plate for cells-to-cDNA synthesis. cDNA was prepared using a cells-to-cDNA kit following the manufacturer's instructions (Ambion).

### 2 Supplementary Figures

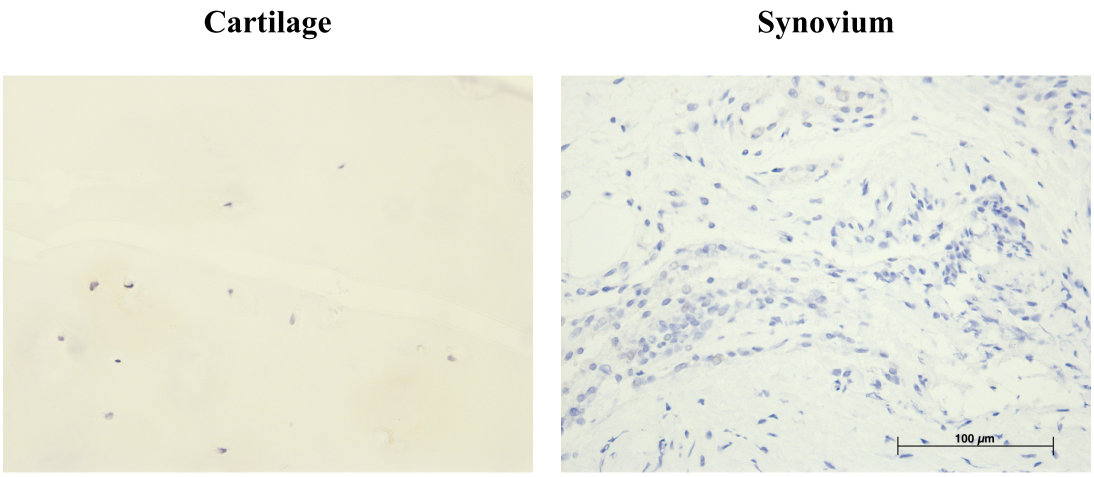


**Supplementary Figure 1.** Immunohistochemistry DAB staining with universal mouse negative control on cartilage or synovium. Antibodies were visualised with DAB (brown) and counterstained with haematoxylin (blue). Images were taken at 40x magnification

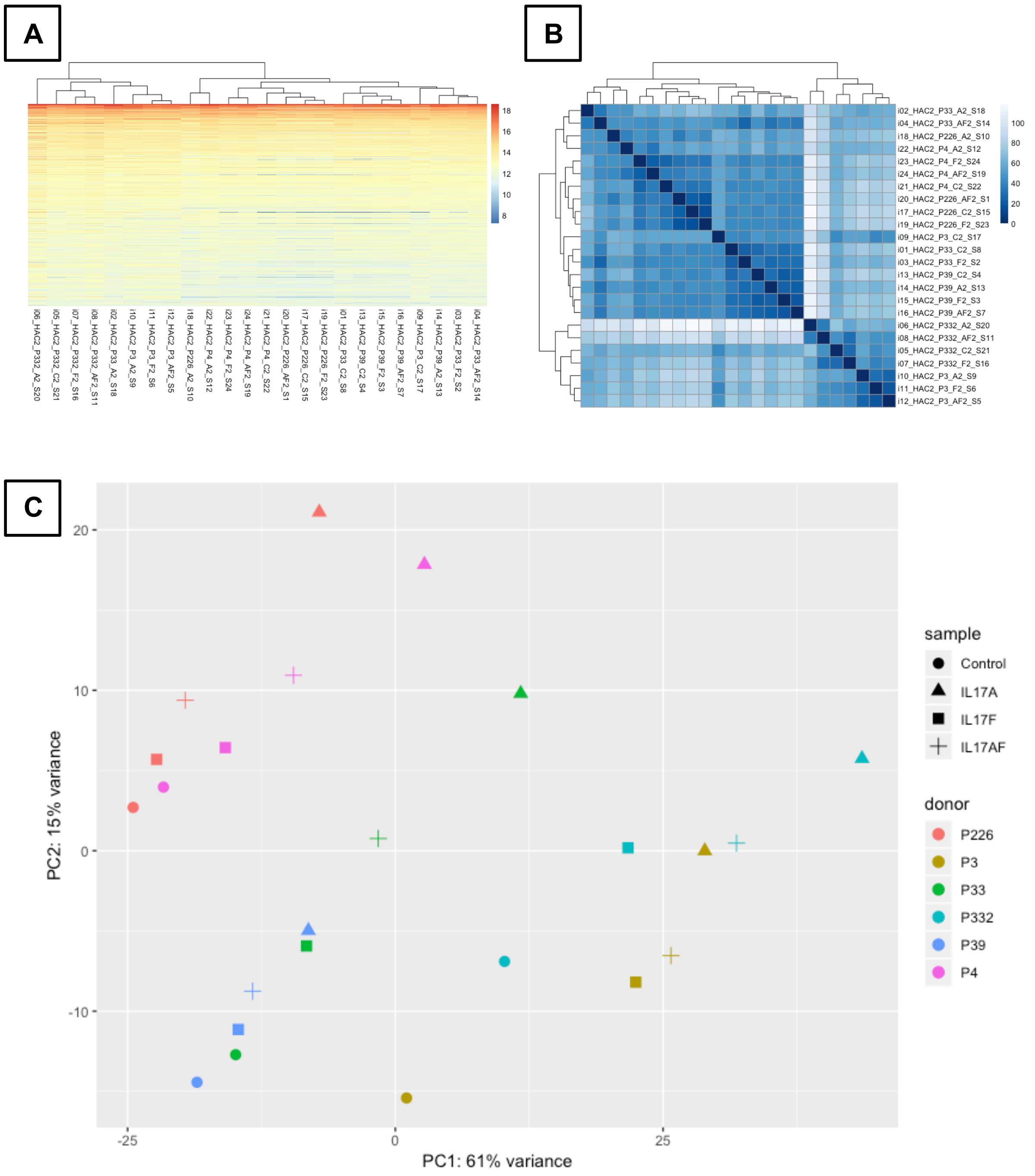


**Supplementary Figure 2.** Overview of different plots showing clustering of the chondrocytes. (A) Heatmap of the counts matrix for chondrocytes samples, blue shows low counts while red shows high counts. (B) Heatmap of the sample-to-sample distances, dark blue represents no to little difference while white represents large differences between samples. (C) Principle component analysis (PCA) plot for chondrocytes samples. Different sample types are represented with different shapes (circle = control, triangle = IL-17A, square = IL-17F, plus sign = IL-17AF) and different donors are represented with different colours (P226 = red, P3 = mustard, P33 = green, P332 = turquoise, P39 = blue, P4 = pink)

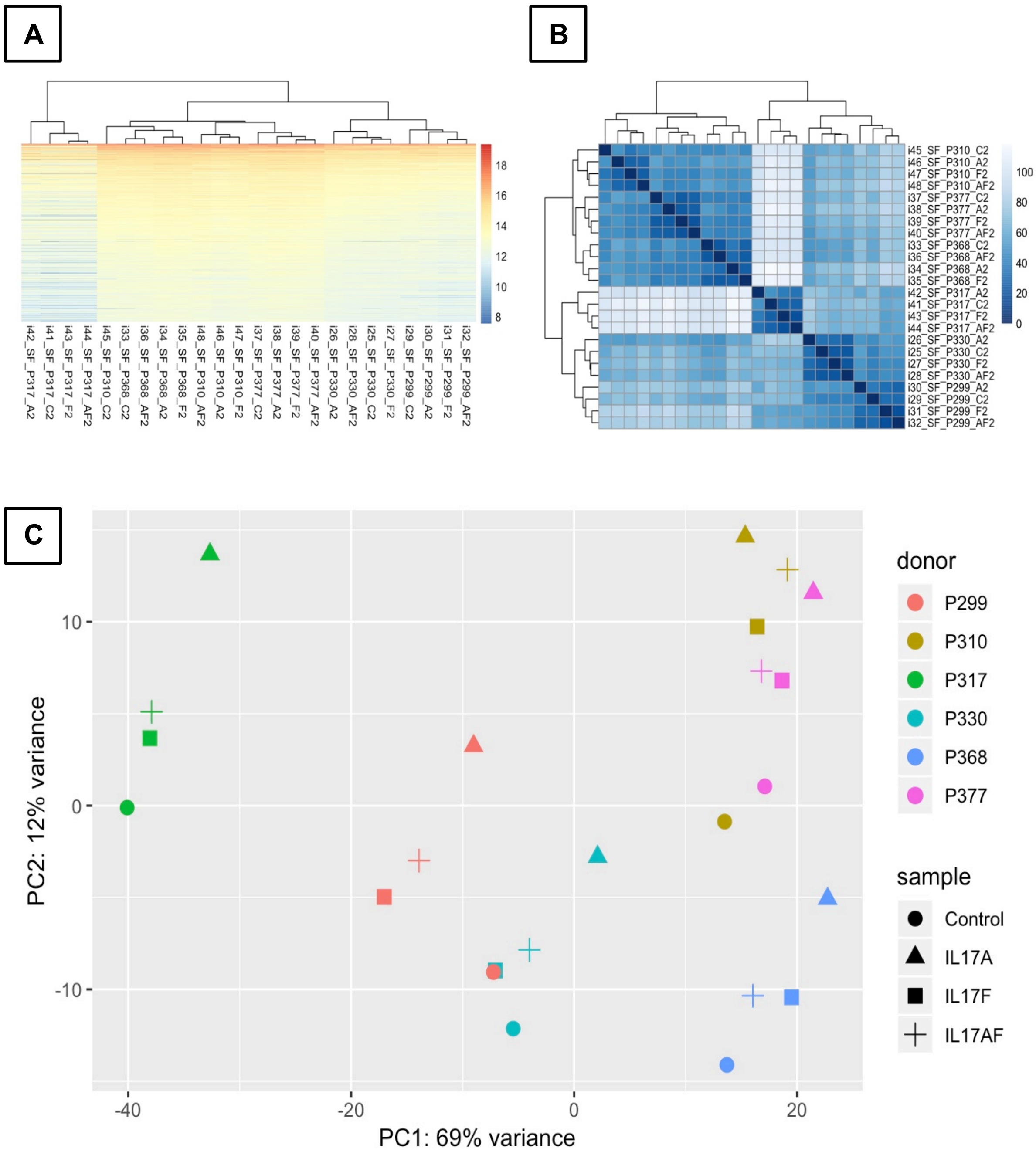


**Supplementary Figure 3.** Overview of different plots showing clustering of the synovial fibroblasts. (A) Heatmap of the counts matrix, blue shows low counts while red shows high counts. (B) Heatmap of the sample-to-sample distances, dark blue represents no to little difference while white represents large differences between samples. (C) Principle component analysis (PCA) plot for synovial fibroblast. Different sample types are represented with different shapes (circle = control, triangle = IL-17A, square = IL-17F, plus sign = IL-17AF) and different donors are represented with different colours (P299 = red, P310 = mustard, P317 = green, P330 = turquoise, P368 = blue, P377 = pink)

**Supplementary Figure 4.** Representative blots for the activation of p65 NF-κB, p38 MAP kinase, and ERK1/2 (p44/p42 MAP kinases) in chondrocytes and synovial fibroblasts by IL-17A, IL-17AF, and IL-17F (all 10 ng/ml) after 10 minutes, 30 minutes, 60 minutes, and 480 minutes (8 hours) of stimulation as can be seen in Figure 5 and Supplementary Figure 7. Blots were stained using vinculin (reference protein), and phosphorylated p38 or phosphorylated p65 NF-κB or phosphorylated ERK1/2, and total p38 or total p65 NF-κB or total ERK1/2, and blots were visualised with an enhanced chemiluminescent reagent.

*(A) Protein lysate from end-stage chondrocytes stained for vinculin, phosphorylated p38, and total p38.*

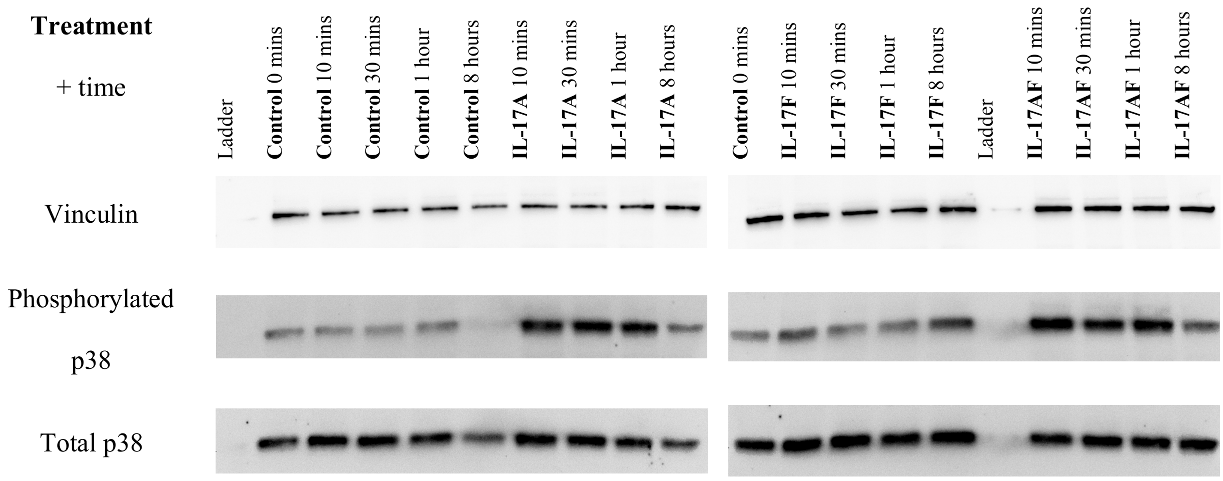


*(B) Protein lysate from end-stage chondrocytes stained for vinculin, phosphorylated* p65 NF-κB*, and total* p65 NF-κB.

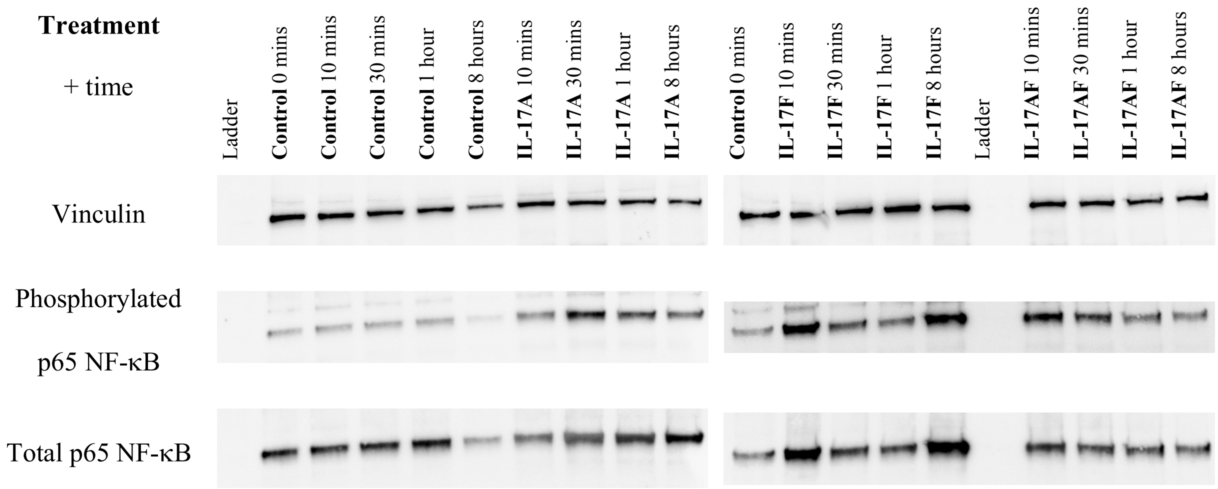

*(C) Protein lysate from end-stage chondrocytes stained for vinculin, phosphorylated* ERK1/2, *and total* ERK1/2.

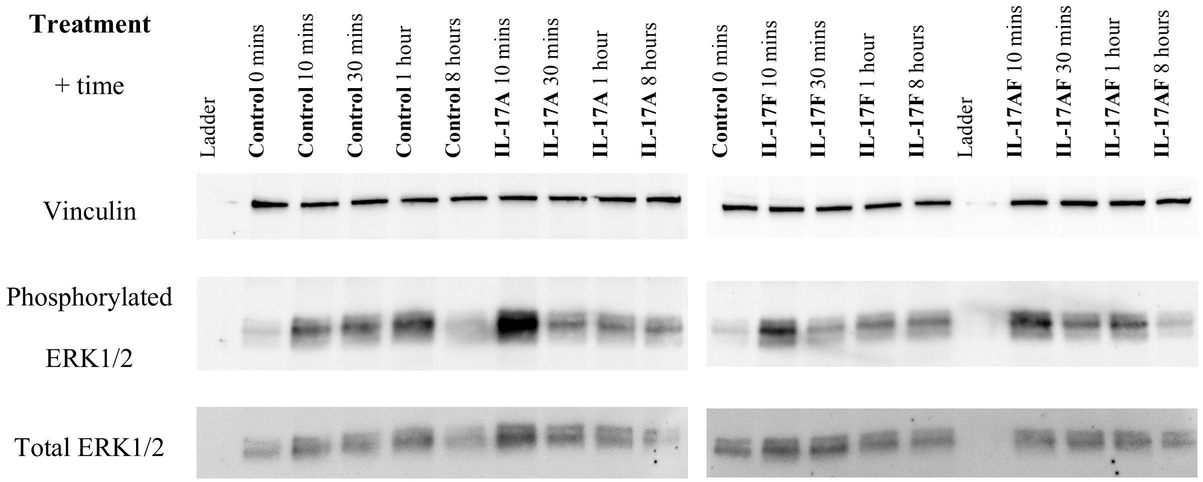


*(D) Protein lysate from end-stage synovial fibroblasts stained for vinculin, phosphorylated p38, and total p38.*

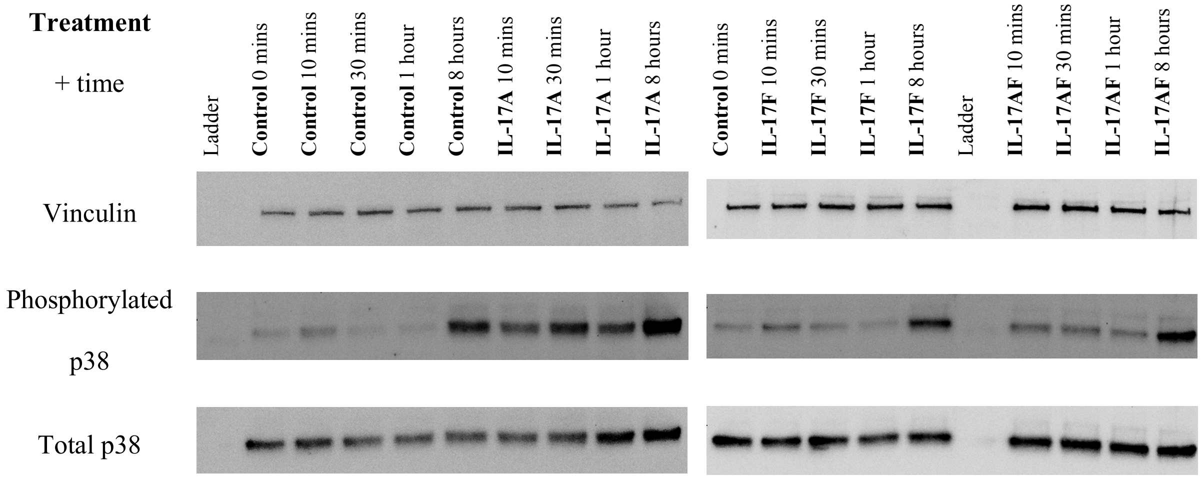

*(E) Protein lysate from end-stage synovial fibroblasts treated with control or IL-17A stained for vinculin, phosphorylated* p65 NF-κB*, and total* p65 NF-κB.

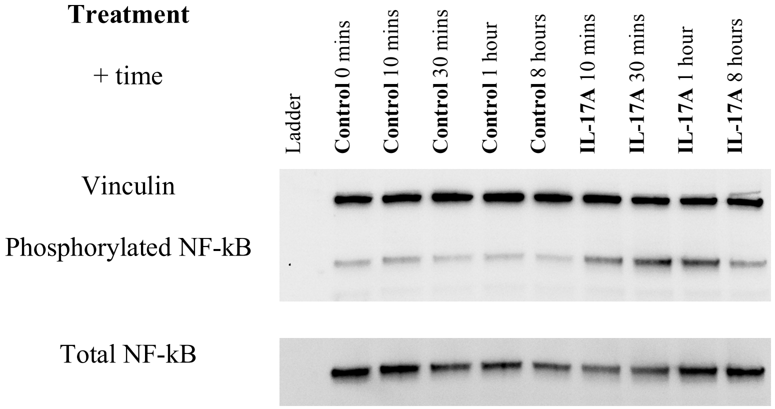


*(F) Protein lysate from end-stage synovial fibroblasts treated with IL-17F and IL-17AF stained for vinculin, phosphorylated* p65 NF-κB*, and total* p65 NF-κB.

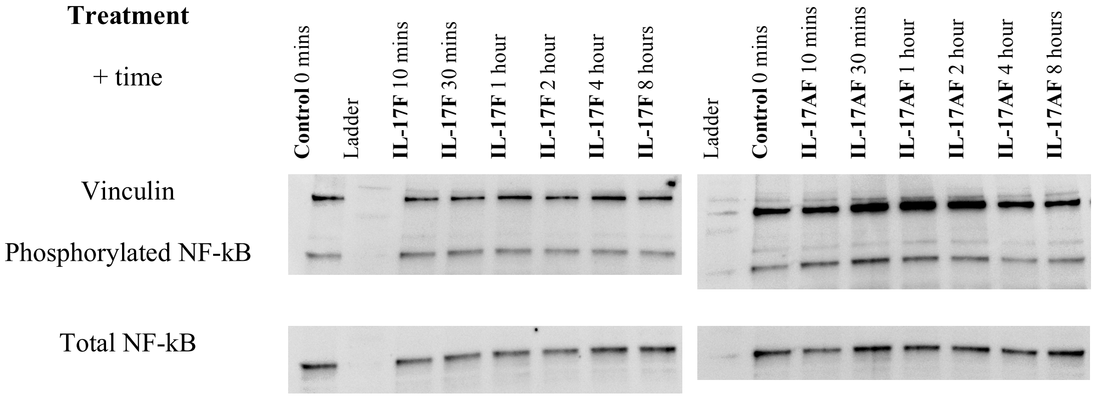

(G) *Protein lysate from end-stage synovial fibroblasts stained for vinculin, phosphorylated* ERK1/2*, and total* ERK1/2.

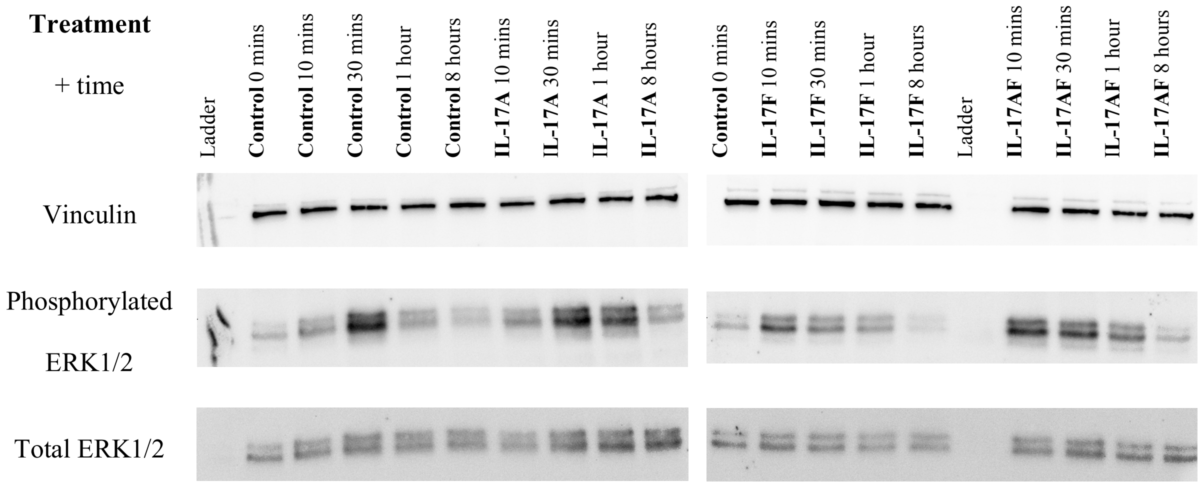

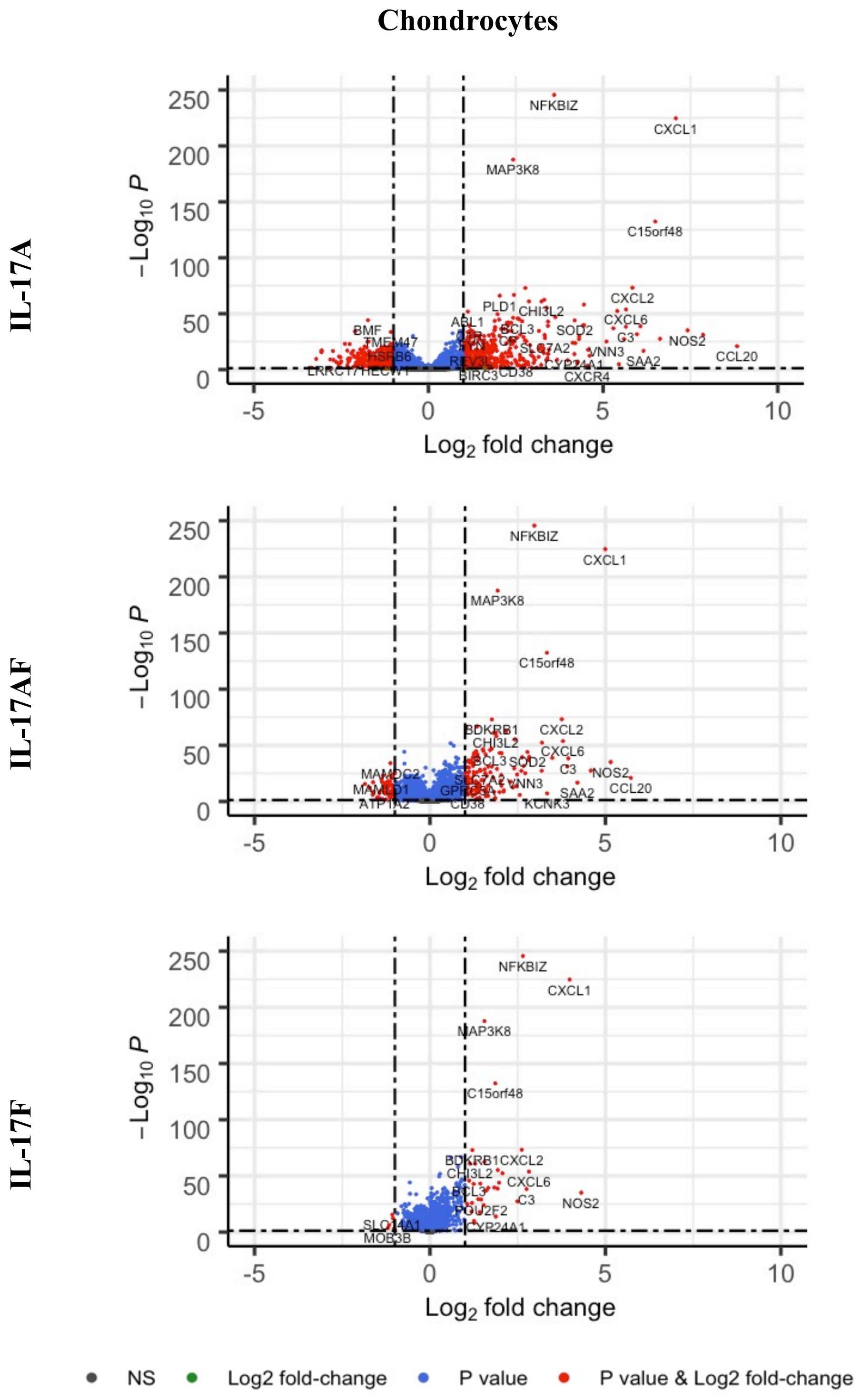


**Supplementary Figure 5.** Volcano plots showing the changes in gene expression in chondrocytes after treatment with IL-17A, IL-17AF, or IL-17F. Grey = not significant (NS); green = Log2 fold change of at least 1 or -1, but p>0.05; blue = p-value < 0.05, but log2 fold change >-1 and <1; red = p-value<0.05 and log2 fold change of at least -1 or +1. n=6

#### Synovial fibroblasts

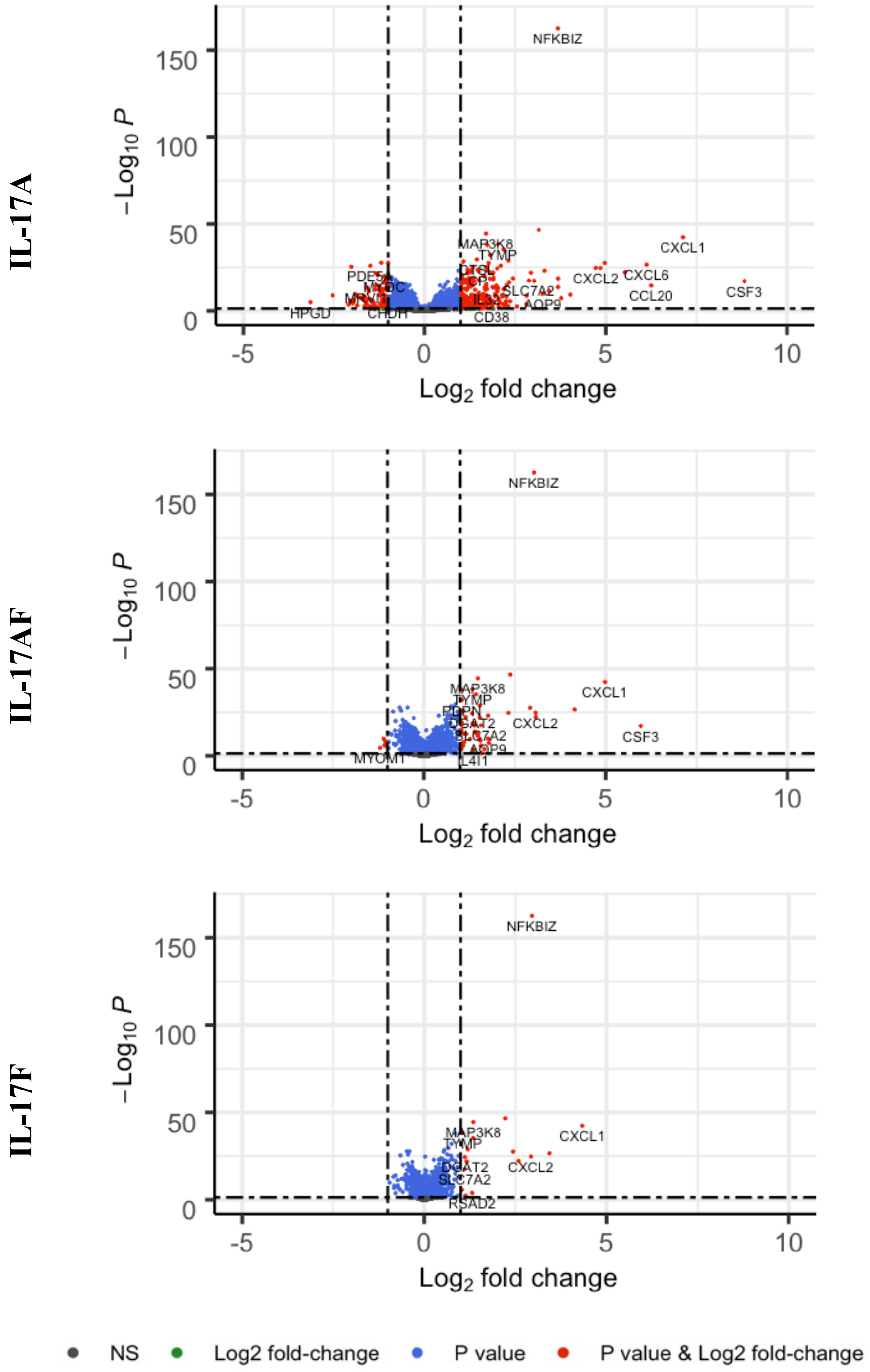


**Supplementary Figure 6.** Volcano plots showing the changes in gene expression in synovial fibroblasts after treatment with IL-17A, IL-17AF, or IL-17F. Grey = not significant (NS); green = Log2 fold change of at least 1 or -1, but p>0.05; blue = p-value < 0.05, but log2 fold change >-1 and <1; red = p-value<0.05 and log2 fold change of at least -1 or +1. n=6

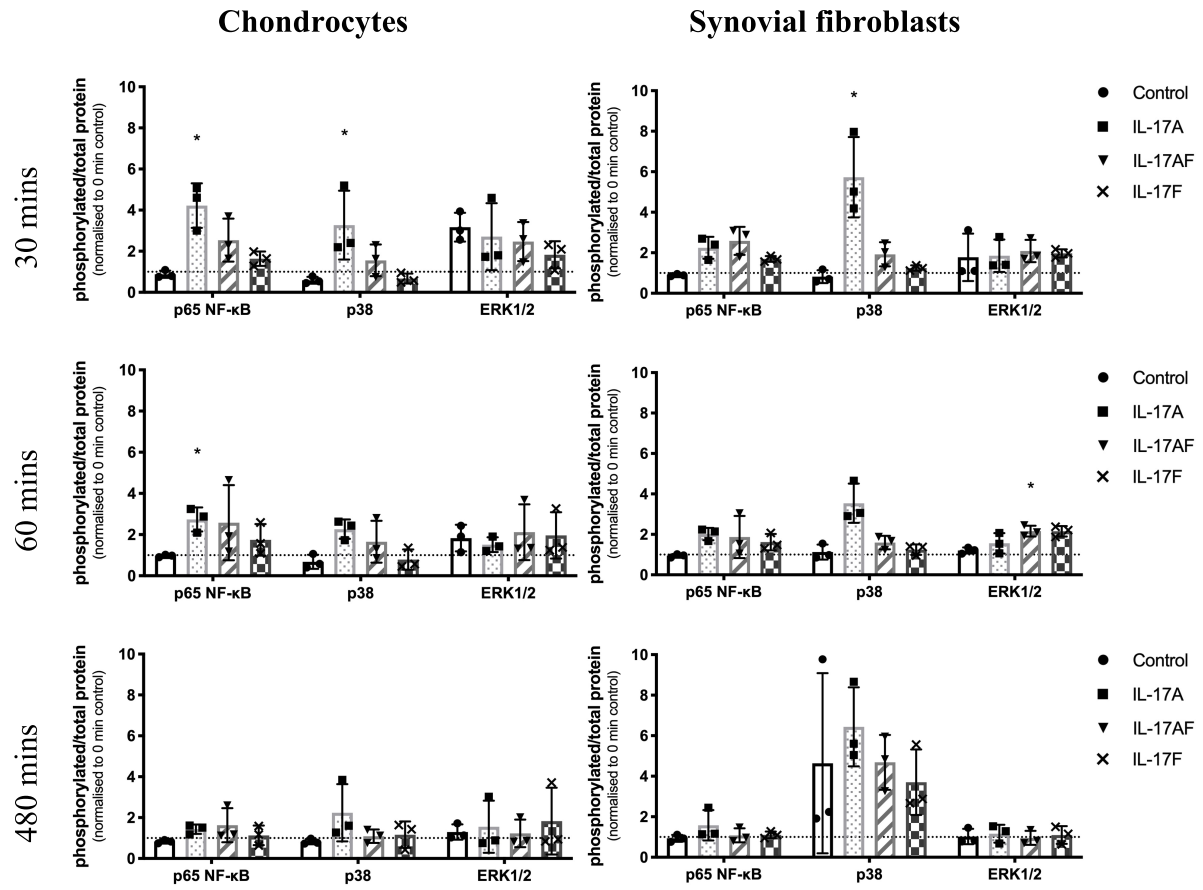


**Supplementary Figure 7.** Activation of p65 NF-κB, p38 MAP kinase, and ERK1/2 (p44/p42 MAP kinases) in chondrocytes and synovial fibroblasts by IL-17A, IL-17AF, and IL-17F (all 10 ng/ml) after 30 minutes, 60 minutes, and 480 minutes (8 hours) of stimulation. Relative ratio of phosphorylated protein over total protein compared to baseline control (0 mins). * = p<0.05, ** = p<0.01, *** = p<0.001, **** = p<0.0001. Friedman test for each intracellular protein with Dunn's multiple comparisons test. Mean±SD. N=3

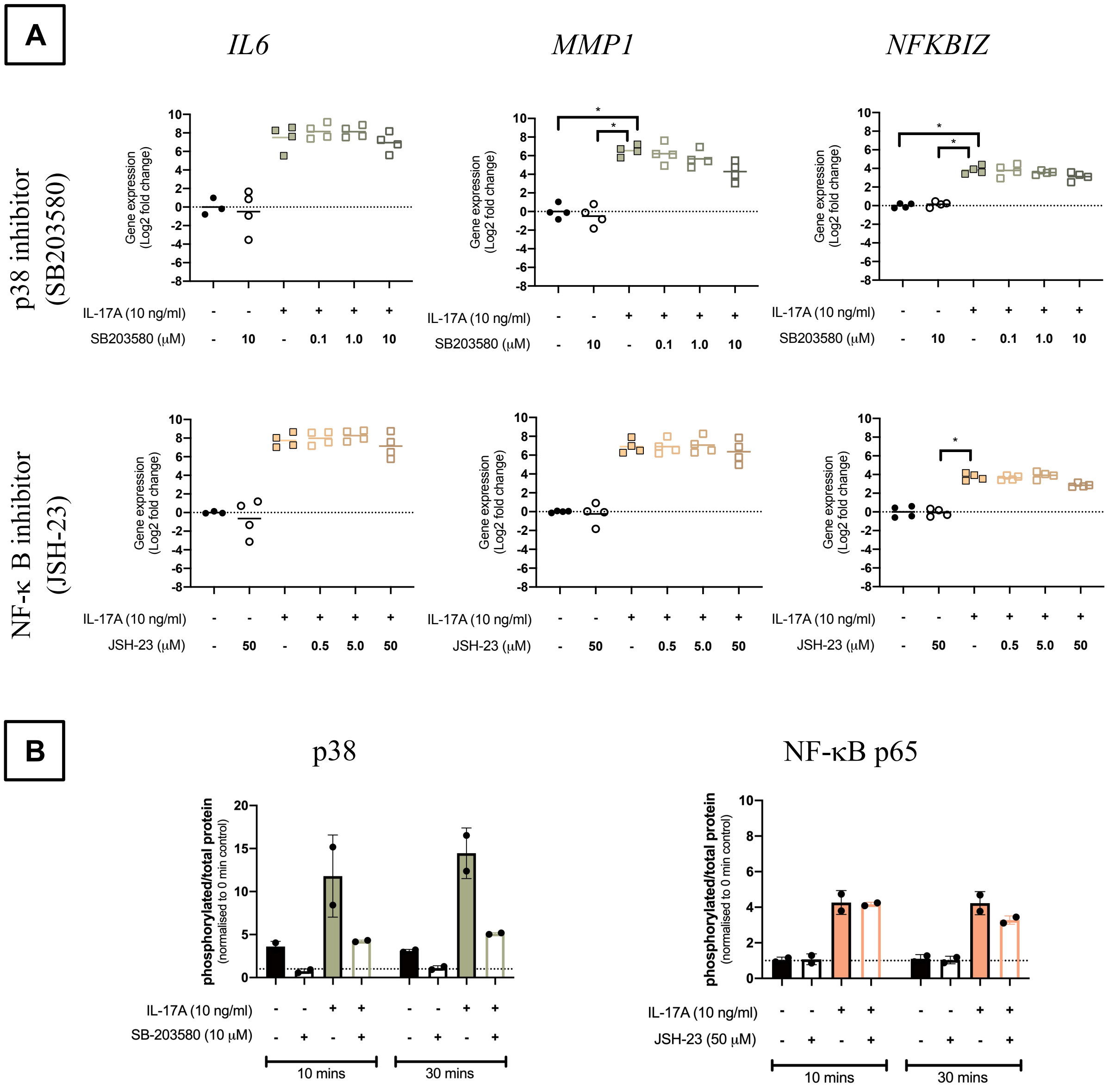


**Supplementary Figure 8.** Chondrocytes were pre-treated for 2 hours with SB203580 (**green**) (Sigma) at 0.1, 1.0, and 10 μM, JSH-23 (**orange**) (Sigma) at 0.5, 5.0, and 50 μM, or vehicle control before the addition of vehicle control or 10 ng/mL IL-17A for 24 hours. (A) mRNA expression of *IL6*, *MMP1*, and *NFKBIZ*. Gene expression was calculated using the ddCT-method using both *GAPDH* and *ACTB* as housekeepers. Gene expression is expressed as log2 fold change compared to control. Changes in gene expression were calculated comparing each treatment to IL-17A treatment alone. Kruskal-Wallis test with Dunn's multiple comparisons test (*IL6*). Friedman test with Dunn's multiple comparisons test (*CXCL1* and *MMP1*). Individual values and mean. N=4. (B) Activation of p38 or NF-κB p65 after 10 and 30 minutes after the start of IL-17A treatment. Relative ratio of phosphorylated protein over total protein compared to baseline control (0 mins). Mean±SD. N=2. * = p<0.05, ** = p<0.01, *** = p<0.001

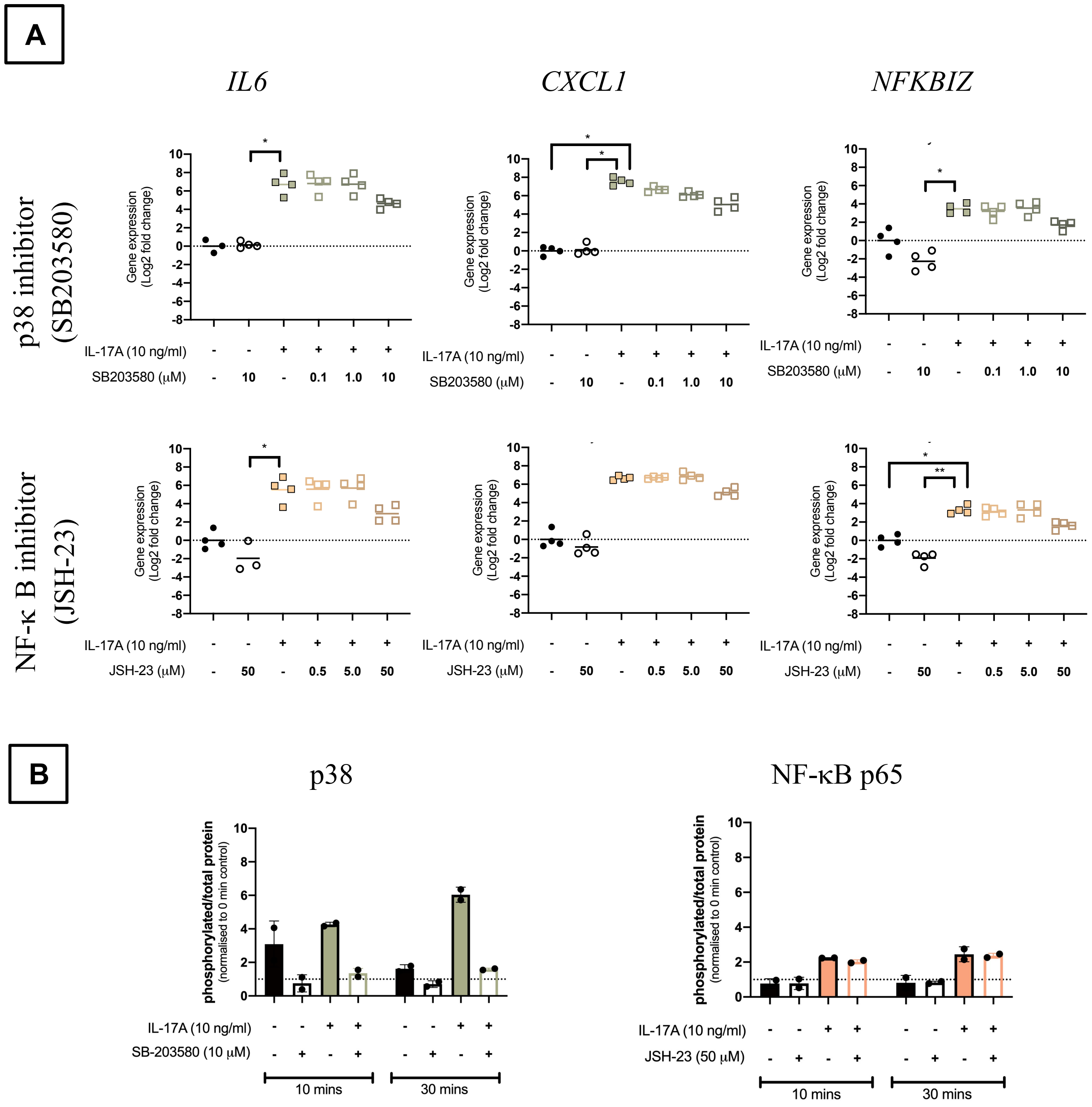


**Supplementary Figure 9.** Synovial fibroblasts were pre-treated for 2 hours with SB203580 (**green**) (Sigma) at 0.1, 1.0, and 10 μM, JSH-23 (**orange**) (Sigma) at 0.5, 5.0, and 50 μM, or vehicle control before the addition of vehicle control or 10 ng/mL IL-17A for 24 hours. (A) mRNA expression of *IL6*, *CXCL1*, and *NFKBIZ*. Gene expression was calculated using the ddCT-method using both *GAPDH* and *ACTB* as housekeepers. Gene expression is expressed as log2 fold change compared to control. Changes in gene expression were calculated comparing each treatment to IL-17A treatment alone. Kruskal-Wallis test with Dunn's multiple comparisons test (*IL6*). Friedman test with Dunn's multiple comparisons test (*CXCL1* and *NFKBIZ*). Individual values and mean. N=4. (B) Activation of p38 or NF-κB p65 after 10 and 30 minutes after the start of IL-17A treatment. Relative ratio of phosphorylated protein over total protein compared to baseline control (0 mins). Mean±SD. N=2. * = $p<0.05$, ** = $p<0.01$, *** = $p<0.001$.

#### 3 Supplementary Tables

**Supplementary Table 1.** Primary antibodies used for histology, including the target, isotype, supplier, reference number, and dilution at which each antibody was used.

| Target | Isotype | Supplier | Ref number | Dilution |
|---|---|---|---|---|
| Human IL-17RA | Monoclonal mouse IgG2B | R&D Systems, Abingdon, UK | MAB1771 | 1:300 |
| Human IL-17RC | Monoclonal mouse IgG2B | R&D Systems | MAB22691 | 1:2000 |
| Universal negative control mouse | Cocktail of mouse IgG1, IgG2a, IgG2b, IgG3, and IgM | Dako, Glostrup, Denmark | IR750 | Ready to use |

**Supplementary Table 2.** Primary and secondary antibodies used for western blot experiments, including target, supplier, reference number, and the dilution at which each antibody was used.

| Target | Supplier | Ref number | Dilution |
|---|---|---|---|
| Vinculin (reference protein) | Cell Signalling Technology, Danvers, MA, USA | 13901 | 1:1000 |
| phospho-ERK1/2 | Cell Signalling Technology | 4370 | 1:2000 |
| ERK1/2 | Cell Signalling Technology | 4695 | 1:1000 |
| phospho-p65 NF-κB | Cell Signalling Technology | 3033 | 1:1000 |
| p65 NF-κB | Cell Signalling Technology | 8242 | 1:1000 |
| phospho-p38 | Cell Signalling Technology | 4511 | 1:1000 |
| p38 | Cell Signalling Technology | 8690 | 1:1000 |
| Anti-rabbit IgG, HRP-linked | Cell Signalling Technology | 7074 | 1:1000 (phospho-antibodies)<br>1:2000 (total-antibodies) |

**Supplementary Table 3.** Overview of primers, including the source of the primers, and the forward and reverse primer sequences where available.

| Target | Supplier | Forward | Reverse |
|---|---|---|---|
| *GAPDH* | Designed in-house | GAAGGTTGAAGGTCGGAGTC | GAAGATGGTGATGGGATTTC |
| *ACTB* | Designed in-house | CCTGGCACCCAGCACAAT | GCCGATCCACACGGAGTACT |
| *IL6* | Primerdesign Ltd | GCAGAAAACAACCTGAACCTT | ACCTCAAACTCCAAAAGACCA |
| *IL17RA* | Primerdesign Ltd | TGAGCACATGCACCACATAC | TTAAGGTTGCGTAGAGTGAGTG |
| *IL17RC* | Primerdesign Ltd | AGTCGTGCTCTCCTTCCAG | AGATTCGTACCTTCACTCCCTAG |
| *CXCL1* | Designed in-house | GGCGGAAAGCTTGCCTCAATCCTG | GCCTCCTTCAGGAACAGCCACCA |
| *MMP1* | Primerdesign Ltd | GCACTGAGAAAGAAGACAAAGG | CTAAGTCCACATCTTGCTCTTG |
| *MMP3* | Primerdesign Ltd | ATGATGAACAATGGACAAAGGATAC | AGTGTTGGCTGAGTGAAAGAG |
| *NFKBIZ* | Qiagen | *Proprietary sequence* | *Proprietary sequence* |
| *MAP3K8* | Designed in-house | ATGGAGTACATGAGCACTGGA | GCTGGCTCTTCACTTGCATAAAG |
| *PTGS2* | Primerdesign Ltd | CAGGCTTCCATTGACCAGAG | TTTCTCCTGTAAGTTCTTCAAATGAT |
| *NOS2* | Qiagen | *Proprietary sequence* | *Proprietary sequence* |
| *CSF3* | Designed in-house | GCTGCTTGAGCCAACTCCATA | GAACGCGGTACGACACCTC |
| *PDPN* | Primerdesign Ltd | CCGAAGATGATGTGGTGACTC | CGATGCGAATGCCTGTTACA |
| *CCL20* | Designed in-house | TGCTGTACCAAGAGTTTGCTC | CGCACACAGACAACTTTTTCTTT |

**Supplementary Table 4.** Overview of differentially expressed genes in chondrocytes between vehicle control and IL-17 treatment (10 ng/ml) with an adjusted p-value of <0.05. Green boxes note logarithmic fold changes (LFC) of at least +1 and red boxes note LFC of at least -1, while blank boxes show an LFC -1 and 1. Genes are sorted alphabetically on SYMBOL gene name.

| **Ensemble gene name** | **SYMBOL gene name** | **Adjusted p-value** | **Log2 Fold Change (LFC)** | | |
|---|---|---|---|---|---|
| | | | IL-17A | IL-17F | IL-17AF |
| ENSG00000183044 | *ABAT* | 3.74E-11 | -1.073 | -0.357 | -0.744 |
| ENSG00000165029 | *ABCA1* | 2.28E-17 | 1.199 | 0.187 | 0.450 |
| ENSG00000167972 | *ABCA3* | 4.89E-10 | -1.475 | -0.491 | -0.923 |
| ENSG00000005471 | *ABCB4* | 7.13E-09 | -1.011 | -0.364 | -0.989 |
| ENSG00000069431 | *ABCC9* | 7.20E-05 | 1.850 | 0.005 | 0.053 |
| ENSG00000143994 | *ABHD1* | 5.23E-03 | -1.571 | -0.026 | -0.177 |
| ENSG00000011198 | *ABHD5* | 3.70E-11 | 1.051 | 0.039 | 0.405 |
| ENSG00000097007 | *ABL1* | 1.35E-52 | 1.128 | 0.285 | 0.589 |
| ENSG00000240303 | *ACAD11* | 3.42E-18 | -1.009 | -0.309 | -0.566 |
| ENSG00000115361 | *ACADL* | 1.18E-03 | -1.519 | -0.026 | -0.234 |
| ENSG00000144476 | *ACKR3* | 5.73E-31 | 2.033 | 0.645 | 1.110 |
| ENSG00000129048 | *ACKR4* | 9.29E-10 | -2.812 | -0.022 | -1.313 |
| ENSG00000122729 | *ACO1* | 7.38E-16 | 1.051 | 0.201 | 0.425 |
| ENSG00000068366 | *ACSL4* | 7.02E-18 | 1.712 | 0.049 | 0.521 |
| ENSG00000107796 | *ACTA2* | 4.53E-08 | -1.184 | -0.328 | -0.727 |
| ENSG00000159251 | *ACTC1* | 1.80E-17 | -2.156 | -0.923 | -1.205 |
| ENSG00000148848 | *ADAM12* | 5.00E-18 | 1.387 | 0.269 | 0.642 |
| ENSG00000042980 | *ADAM28* | 1.27E-14 | 3.424 | 0.026 | 2.038 |
| ENSG00000138316 | *ADAMTS14* | 2.34E-17 | -2.734 | -0.926 | -1.281 |
| ENSG00000166106 | *ADAMTS15* | 8.81E-09 | -1.549 | -0.684 | -1.166 |
| ENSG00000049192 | *ADAMTS6* | 4.16E-04 | 1.158 | 0.015 | 0.109 |
| ENSG00000163638 | *ADAMTS9* | 2.97E-05 | 1.616 | 0.013 | 0.347 |
| ENSG00000173567 | *ADGRF3* | 1.50E-02 | -1.032 | -0.027 | -0.256 |
| ENSG00000198099 | *ADH4* | 1.85E-02 | 1.357 | 0.018 | 0.207 |
| ENSG00000196344 | *ADH7* | 8.39E-04 | -1.117 | -0.019 | -0.178 |
| ENSG00000128271 | *ADORA2A* | 5.07E-03 | 1.074 | 0.001 | 0.096 |
| ENSG00000150594 | *ADRA2A* | 1.96E-09 | -2.284 | -0.040 | -1.134 |
| ENSG00000111863 | *ADTRP* | 1.73E-06 | 1.270 | 0.052 | 0.545 |
| ENSG00000144218 | *AFF3* | 8.87E-08 | -2.297 | -0.025 | -0.980 |
| ENSG00000187546 | *AGMO* | 1.97E-02 | -1.307 | -0.022 | -0.183 |
| ENSG00000026652 | *AGPAT4* | 1.57E-10 | 1.227 | 0.047 | 0.421 |
| ENSG00000162433 | *AK4* | 5.46E-09 | 1.092 | 0.102 | 0.484 |

| | | | | | |
|---|---|---|---|---|---|
| ENSG00000154027 | *AK5* | 2.21E-12 | -2.558 | -0.040 | -1.029 |
| ENSG00000131016 | *AKAP12* | 1.66E-14 | -1.242 | -0.704 | -1.268 |
| ENSG00000151320 | *AKAP6* | 4.58E-09 | -1.153 | -0.478 | -1.004 |
| ENSG00000227471 | *AKR1B15* | 2.50E-03 | 1.352 | 0.027 | 0.852 |
| ENSG00000187134 | *AKR1C1* | 3.14E-07 | 1.965 | 0.750 | 1.276 |
| ENSG00000151632 | *AKR1C2* | 2.03E-07 | 1.269 | 0.644 | 1.006 |
| ENSG00000144908 | *ALDH1L1* | 4.75E-02 | -1.196 | -0.019 | -0.063 |
| ENSG00000112294 | *ALDH5A1* | 6.15E-08 | -1.341 | -0.070 | -0.821 |
| ENSG00000133805 | *AMPD3* | 4.23E-12 | 1.453 | 0.045 | 0.424 |
| ENSG00000091879 | *ANGPT2* | 3.13E-10 | -1.587 | -0.040 | -0.828 |
| ENSG00000136859 | *ANGPTL2* | 7.50E-14 | 1.008 | 0.308 | 0.677 |
| ENSG00000167772 | *ANGPTL4* | 3.14E-09 | 1.289 | 0.360 | 0.858 |
| ENSG00000165887 | *ANKRD2* | 1.50E-02 | -1.636 | -0.019 | -0.122 |
| ENSG00000089847 | *ANKRD24* | 1.18E-03 | -1.049 | -0.035 | -0.838 |
| ENSG00000138772 | *ANXA3* | 5.96E-03 | -1.096 | 0.001 | -0.096 |
| ENSG00000138356 | *AOX1* | 2.39E-09 | 1.343 | 0.033 | 0.397 |
| ENSG00000154856 | *APCDD1* | 3.94E-09 | 1.367 | 0.395 | 0.864 |
| ENSG00000171388 | *APLN* | 3.82E-06 | 2.090 | 0.003 | 0.113 |
| ENSG00000084674 | *APOB* | 2.51E-02 | -1.071 | -0.015 | -0.210 |
| ENSG00000128284 | *APOL3* | 4.77E-13 | 1.100 | 0.283 | 0.565 |
| ENSG00000136044 | *APPL2* | 3.06E-23 | -1.084 | -0.345 | -0.595 |
| ENSG00000165272 | *AQP3* | 1.33E-14 | -1.623 | -0.083 | -0.400 |
| ENSG00000165269 | *AQP7* | 2.07E-10 | -2.289 | -0.602 | -1.063 |
| ENSG00000103569 | *AQP9* | 1.60E-02 | 3.101 | 0.005 | 0.045 |
| ENSG00000169083 | *AR* | 1.38E-12 | -1.150 | -0.415 | -0.719 |
| ENSG00000146376 | *ARHGAP18* | 2.83E-05 | -1.089 | -0.038 | -0.614 |
| ENSG00000137727 | *ARHGAP20* | 1.77E-08 | -1.442 | -0.053 | -0.853 |
| ENSG00000111348 | *ARHGDIB* | 7.34E-06 | -1.345 | -0.482 | -0.334 |
| ENSG00000183111 | *ARHGEF37* | 1.34E-07 | -1.136 | -0.283 | -0.733 |
| ENSG00000116017 | *ARID3A* | 1.49E-19 | 1.072 | 0.129 | 0.485 |
| ENSG00000169126 | *ARMC4* | 2.72E-03 | -1.309 | -0.027 | -0.391 |
| ENSG00000113369 | *ARRDC3* | 4.44E-16 | 1.691 | 0.060 | 0.606 |
| ENSG00000099889 | *ARVCF* | 4.56E-11 | -1.192 | -0.077 | -0.524 |
| ENSG00000110881 | *ASIC1* | 1.21E-06 | -1.006 | -0.072 | -0.563 |
| ENSG00000108381 | *ASPA* | 7.01E-03 | -1.037 | -0.027 | -0.359 |
| ENSG00000106819 | *ASPN* | 1.12E-15 | -1.910 | -0.606 | -1.141 |
| ENSG00000130707 | *ASS1* | 1.20E-23 | 1.190 | 0.346 | 0.735 |
| ENSG00000148219 | *ASTN2* | 8.34E-04 | -1.119 | -0.028 | -0.440 |
| ENSG00000162772 | *ATF3* | 1.50E-08 | 1.265 | 0.101 | 1.044 |
| ENSG00000018625 | *ATP1A2* | 2.36E-08 | -2.275 | -0.755 | -1.289 |

| | | | | | |
|---|---|---|---|---|---|
| ENSG00000070961 | *ATP2B1* | 7.01E-08 | 1.003 | 0.076 | 0.355 |
| ENSG00000132932 | *ATP8A2* | 1.27E-02 | -0.792 | -0.019 | -1.255 |
| ENSG00000104043 | *ATP8B4* | 9.96E-05 | 1.323 | 0.023 | 0.374 |
| ENSG00000107518 | *ATRNL1* | 9.38E-08 | -1.002 | -0.285 | -0.624 |
| ENSG00000119986 | *AVPI1* | 9.65E-19 | 1.141 | 0.359 | 0.722 |
| ENSG00000162630 | *B3GALT2* | 2.98E-10 | -3.217 | -0.011 | -0.179 |
| ENSG00000086062 | *B4GALT1* | 1.19E-15 | 1.001 | 0.105 | 0.436 |
| ENSG00000164929 | *BAALC* | 4.05E-04 | -1.115 | -0.011 | -0.211 |
| ENSG00000095739 | *BAMBI* | 6.21E-10 | -1.034 | -0.039 | -0.270 |
| ENSG00000176788 | *BASP1* | 1.07E-13 | 1.150 | 0.431 | 0.836 |
| ENSG00000156127 | *BATF* | 4.55E-04 | 1.203 | 0.003 | 0.105 |
| ENSG00000140379 | *BCL2A1* | 3.52E-03 | 2.286 | 0.001 | 0.042 |
| ENSG00000069399 | *BCL3* | 1.74E-46 | 2.557 | 1.118 | 1.713 |
| ENSG00000113916 | *BCL6* | 9.25E-15 | 1.181 | 0.252 | 0.600 |
| ENSG00000100739 | *BDKRB1* | 9.95E-74 | 2.772 | 1.207 | 1.764 |
| ENSG00000168398 | *BDKRB2* | 5.42E-23 | 1.883 | 0.723 | 1.086 |
| ENSG00000176697 | *BDNF* | 1.35E-04 | 1.121 | 0.016 | 0.198 |
| ENSG00000167995 | *BEST1* | 7.21E-31 | 1.871 | 0.425 | 0.971 |
| ENSG00000134107 | *BHLHE40* | 2.26E-08 | 1.342 | 0.067 | 0.742 |
| ENSG00000123095 | *BHLHE41* | 7.70E-24 | 1.313 | 0.123 | 0.421 |
| ENSG00000015475 | *BID* | 3.25E-24 | 1.017 | 0.288 | 0.452 |
| ENSG00000023445 | *BIRC3* | 7.20E-05 | 1.436 | 0.007 | 0.072 |
| ENSG00000104081 | *BMF* | 6.54E-45 | -1.733 | -0.574 | -0.733 |
| ENSG00000125845 | *BMP2* | 4.94E-03 | 1.038 | 0.010 | 0.185 |
| ENSG00000153162 | *BMP6* | 6.62E-13 | 2.375 | 0.031 | 1.093 |
| ENSG00000078725 | *BRINP1* | 6.40E-05 | 1.137 | 0.023 | 0.207 |
| ENSG00000133639 | *BTG1* | 1.53E-15 | 1.059 | 0.386 | 0.651 |
| ENSG00000154493 | *C10orf90* | 6.09E-14 | 1.446 | 0.791 | 1.073 |
| ENSG00000120055 | *C10orf95* | 1.61E-02 | -1.150 | -0.016 | -0.148 |
| ENSG00000110665 | *C11orf21* | 1.46E-02 | -1.137 | -0.004 | -0.130 |
| ENSG00000174370 | *C11orf45* | 4.37E-07 | -2.072 | -1.161 | -1.111 |
| ENSG00000149300 | *C11orf52* | 3.30E-05 | -1.092 | -0.380 | -0.853 |
| ENSG00000187479 | *C11orf96* | 1.00E-21 | 3.216 | 0.948 | 1.801 |
| ENSG00000235162 | *C12orf75* | 1.06E-23 | -1.036 | -0.370 | -0.537 |
| ENSG00000166920 | *C15orf48* | 4.24E-133 | 6.492 | 1.860 | 3.333 |
| ENSG00000132016 | *C19orf57* | 1.87E-03 | -1.231 | -0.032 | -0.123 |
| ENSG00000106392 | *C1GALT1* | 4.18E-37 | 1.908 | 0.703 | 1.091 |
| ENSG00000173918 | *C1QTNF1* | 6.70E-21 | 1.844 | 0.027 | 0.602 |
| ENSG00000145861 | *C1QTNF2* | 6.45E-10 | -1.790 | -0.048 | -0.804 |
| ENSG00000163145 | *C1QTNF7* | 4.42E-10 | -2.307 | -0.748 | -1.415 |

| | | | | | |
|---|---|---|---|---|---|
| ENSG00000205863 | *C1QTNF9B* | 8.86E-03 | -1.211 | -0.020 | -0.088 |
| ENSG00000159403 | *C1R* | 6.20E-40 | 1.174 | 0.413 | 0.742 |
| ENSG00000139178 | *C1RL* | 7.07E-35 | 1.241 | 0.455 | 0.750 |
| ENSG00000198535 | *C2CD4A* | 1.97E-06 | 4.262 | 0.015 | 2.558 |
| ENSG00000205502 | *C2CD4B* | 1.51E-05 | 5.453 | 0.009 | 0.089 |
| ENSG00000125730 | *C3* | 4.39E-39 | 5.653 | 2.751 | 3.940 |
| ENSG00000197405 | *C5AR1* | 5.40E-05 | 1.264 | 0.016 | 0.141 |
| ENSG00000178776 | *C5orf46* | 4.35E-02 | 1.714 | 0.007 | 0.056 |
| ENSG00000074410 | *CA12* | 3.53E-35 | 2.385 | 0.658 | 1.177 |
| ENSG00000107159 | *CA9* | 3.95E-04 | 1.660 | 0.024 | 0.215 |
| ENSG00000183346 | *CABCOCO1* | 3.51E-08 | -1.205 | -0.119 | -0.604 |
| ENSG00000070808 | *CAMK2A* | 1.08E-05 | -1.522 | -0.052 | -1.159 |
| ENSG00000058404 | *CAMK2B* | 4.19E-04 | -2.109 | -0.023 | -0.775 |
| ENSG00000105519 | *CAPS* | 9.33E-10 | -1.012 | -0.134 | -0.419 |
| ENSG00000204682 | *CASC10* | 4.69E-08 | -1.645 | -0.044 | -1.292 |
| ENSG00000168497 | *CAVIN2* | 5.99E-14 | -2.012 | -0.077 | -1.198 |
| ENSG00000150636 | *CCDC102B* | 1.60E-03 | -1.307 | -0.018 | -0.121 |
| ENSG00000168491 | *CCDC110* | 1.67E-07 | -1.046 | -0.402 | -0.800 |
| ENSG00000198003 | *CCDC151* | 2.12E-02 | -1.032 | -0.024 | -0.216 |
| ENSG00000166510 | *CCDC68* | 1.39E-02 | 1.033 | 0.021 | 0.115 |
| ENSG00000055813 | *CCDC85A* | 3.05E-12 | -1.225 | -0.043 | -0.536 |
| ENSG00000181374 | *CCL13* | 4.54E-02 | 1.739 | 0.001 | 0.035 |
| ENSG00000108691 | *CCL2* | 1.17E-24 | 4.129 | 1.530 | 2.484 |
| ENSG00000115009 | *CCL20* | 1.03E-21 | 8.836 | 0.022 | 5.726 |
| ENSG00000213927 | *CCL27* | 1.11E-02 | -1.397 | -0.011 | -0.136 |
| ENSG00000108688 | *CCL7* | 3.78E-13 | 4.614 | 0.022 | 2.363 |
| ENSG00000108700 | *CCL8* | 4.55E-28 | 5.598 | 0.010 | 2.603 |
| ENSG00000135083 | *CCNJL* | 1.29E-02 | -1.102 | -0.031 | -0.175 |
| ENSG00000163823 | *CCR1* | 1.52E-02 | 1.585 | 0.005 | 0.088 |
| ENSG00000126353 | *CCR7* | 1.31E-15 | 2.201 | 0.785 | 0.912 |
| ENSG00000158473 | *CD1D* | 4.79E-19 | 3.168 | 1.179 | 2.009 |
| ENSG00000174807 | *CD248* | 3.03E-02 | -1.244 | -0.008 | -0.109 |
| ENSG00000004468 | *CD38* | 7.16E-08 | 2.508 | 0.009 | 1.078 |
| ENSG00000085117 | *CD82* | 1.69E-15 | 1.081 | 0.156 | 0.464 |
| ENSG00000074276 | *CDHR2* | 4.85E-02 | -1.122 | -0.020 | -0.076 |
| ENSG00000138395 | *CDK15* | 1.80E-02 | -2.317 | -0.011 | -0.071 |
| ENSG00000138769 | *CDKL2* | 2.99E-03 | -1.391 | -0.023 | -0.617 |
| ENSG00000129596 | *CDO1* | 3.01E-11 | 1.199 | 0.416 | 0.669 |
| ENSG00000172216 | *CEBPB* | 2.40E-32 | 1.838 | 0.761 | 1.227 |
| ENSG00000221869 | *CEBPD* | 7.11E-44 | 2.682 | 1.434 | 1.985 |

| | | | | | |
|---|---|---|---|---|---|
| ENSG00000184524 | *CEND1* | 2.02E-03 | 1.365 | 0.017 | 0.690 |
| ENSG00000188452 | *CERKL* | 1.47E-08 | -1.523 | -0.615 | -0.657 |
| ENSG00000105792 | *CFAP69* | 1.68E-18 | 1.933 | 0.051 | 0.917 |
| ENSG00000243649 | *CFB* | 1.31E-22 | 2.906 | 0.030 | 0.834 |
| ENSG00000128849 | *CGNL1* | 2.59E-07 | 1.058 | 0.031 | 0.204 |
| ENSG00000138135 | *CH25H* | 3.44E-26 | 2.801 | 0.030 | 1.085 |
| ENSG00000100399 | *CHADL* | 5.07E-07 | -1.652 | -0.026 | -0.516 |
| ENSG00000016391 | *CHDH* | 1.53E-15 | -1.299 | -0.283 | -0.471 |
| ENSG00000183765 | *CHEK2* | 1.11E-10 | 1.379 | 0.479 | 0.809 |
| ENSG00000133048 | *CHI3L1* | 2.71E-44 | 1.892 | 0.758 | 1.273 |
| ENSG00000064886 | *CHI3L2* | 1.22E-61 | 3.236 | 1.137 | 1.880 |
| ENSG00000255112 | *CHMP1B* | 4.05E-18 | 1.514 | 0.451 | 0.876 |
| ENSG00000128656 | *CHN1* | 1.12E-12 | -1.419 | -0.039 | -0.429 |
| ENSG00000054938 | *CHRDL2* | 6.11E-05 | 1.706 | 0.011 | 0.192 |
| ENSG00000171310 | *CHST11* | 2.96E-15 | 1.547 | 0.042 | 0.508 |
| ENSG00000163815 | *CLEC3B* | 2.87E-05 | -1.061 | -0.063 | -0.451 |
| ENSG00000169583 | *CLIC3* | 2.64E-07 | -1.020 | -0.330 | -0.436 |
| ENSG00000159212 | *CLIC6* | 4.74E-21 | 1.064 | 0.157 | 0.546 |
| ENSG00000165959 | *CLMN* | 2.29E-10 | 1.448 | 0.354 | 0.634 |
| ENSG00000149970 | *CNKSR2* | 3.19E-04 | -1.114 | -0.029 | -0.288 |
| ENSG00000153721 | *CNKSR3* | 1.78E-17 | -1.189 | -0.300 | -0.547 |
| ENSG00000103647 | *CORO2B* | 3.84E-07 | -1.076 | -0.417 | -0.594 |
| ENSG00000047457 | *CP* | 3.73E-35 | 2.279 | 0.759 | 1.220 |
| ENSG00000177685 | *CRACR2B* | 2.73E-11 | -1.369 | -0.021 | -0.315 |
| ENSG00000213145 | *CRIP1* | 2.06E-19 | -1.028 | -0.215 | -0.353 |
| ENSG00000121005 | *CRISPLD1* | 7.42E-26 | -1.716 | -0.598 | -1.002 |
| ENSG00000103196 | *CRISPLD2* | 5.89E-08 | 1.684 | 0.015 | 0.326 |
| ENSG00000226321 | *CROCC2* | 8.83E-03 | 1.235 | 0.000 | 0.145 |
| ENSG00000095713 | *CRTAC1* | 2.18E-13 | 1.499 | 0.045 | 0.664 |
| ENSG00000109943 | *CRTAM* | 2.23E-03 | 1.835 | 0.010 | 0.113 |
| ENSG00000172346 | *CSDC2* | 6.74E-05 | -1.402 | -0.053 | -0.553 |
| ENSG00000178662 | *CSRNP3* | 1.80E-04 | -2.323 | -0.027 | -1.496 |
| ENSG00000118523 | *CTGF* | 1.27E-14 | -1.700 | -0.413 | -0.908 |
| ENSG00000135047 | *CTSL* | 1.61E-06 | 1.024 | 0.084 | 0.716 |
| ENSG00000163131 | *CTSS* | 7.87E-39 | 1.910 | 0.366 | 0.803 |
| ENSG00000006210 | *CX3CL1* | 2.52E-05 | 1.445 | 0.008 | 0.112 |
| ENSG00000163739 | *CXCL1* | 1.92E-225 | 7.081 | 3.981 | 4.993 |
| ENSG00000081041 | *CXCL2* | 5.60E-74 | 5.839 | 2.616 | 3.757 |
| ENSG00000163734 | *CXCL3* | 4.31E-53 | 5.404 | 2.064 | 3.190 |
| ENSG00000163735 | *CXCL5* | 2.00E-18 | 4.569 | 0.023 | 2.300 |

| | | | | | |
|---|---|---|---|---|---|
| ENSG00000124875 | *CXCL6* | 1.63E-54 | 5.656 | 2.821 | 3.790 |
| ENSG00000169429 | *CXCL8* | 2.09E-32 | 5.969 | 0.035 | 2.672 |
| ENSG00000121966 | *CXCR4* | 3.15E-03 | 4.545 | 0.007 | 0.042 |
| ENSG00000137869 | *CYP19A1* | 1.41E-02 | 1.259 | 0.012 | 0.161 |
| ENSG00000019186 | *CYP24A1* | 1.29E-14 | 4.175 | 1.879 | 2.462 |
| ENSG00000146233 | *CYP39A1* | 1.20E-08 | -1.949 | -0.535 | -0.991 |
| ENSG00000186377 | *CYP4X1* | 3.75E-04 | -1.046 | -0.027 | -0.310 |
| ENSG00000172817 | *CYP7B1* | 2.17E-10 | 1.557 | 0.501 | 0.659 |
| ENSG00000205795 | *CYS1* | 3.28E-13 | -1.851 | -0.389 | -0.888 |
| ENSG00000100592 | *DAAM1* | 2.96E-20 | 1.149 | 0.175 | 0.420 |
| ENSG00000146122 | *DAAM2* | 1.98E-07 | -1.014 | -0.089 | -0.593 |
| ENSG00000163331 | *DAPL1* | 1.26E-06 | -2.523 | -0.015 | -1.630 |
| ENSG00000003249 | *DBNDD1* | 1.76E-05 | -1.215 | -0.034 | -0.648 |
| ENSG00000105516 | *DBP* | 2.01E-06 | -1.056 | -0.053 | -0.330 |
| ENSG00000133083 | *DCLK1* | 7.01E-03 | -1.515 | -0.018 | -0.070 |
| ENSG00000163357 | *DCST1* | 3.81E-02 | -1.075 | -0.009 | -0.083 |
| ENSG00000153904 | *DDAH1* | 9.92E-19 | -1.398 | -0.400 | -0.813 |
| ENSG00000168209 | *DDIT4* | 4.35E-05 | 1.702 | 0.018 | 0.355 |
| ENSG00000145358 | *DDIT4L* | 5.91E-04 | 1.253 | 0.012 | 0.162 |
| ENSG00000164825 | *DEFB1* | 3.61E-03 | 1.231 | 0.021 | 0.088 |
| ENSG00000165507 | *DEPP1* | 3.82E-10 | 1.926 | 0.046 | 0.972 |
| ENSG00000155792 | *DEPTOR* | 1.63E-15 | -2.207 | -0.083 | -0.908 |
| ENSG00000062282 | *DGAT2* | 9.04E-12 | 1.749 | 0.630 | 1.305 |
| ENSG00000077044 | *DGKD* | 1.81E-17 | 1.152 | 0.326 | 0.655 |
| ENSG00000197406 | *DIO3* | 9.87E-03 | 2.942 | 0.008 | 0.049 |
| ENSG00000140323 | *DISP2* | 7.79E-04 | -1.275 | -0.017 | -0.984 |
| ENSG00000132837 | *DMGDH* | 4.38E-07 | -1.143 | -0.032 | -0.200 |
| ENSG00000115423 | *DNAH6* | 4.98E-02 | -1.167 | -0.005 | -0.103 |
| ENSG00000187957 | *DNER* | 2.10E-08 | 1.899 | 0.014 | 0.134 |
| ENSG00000149927 | *DOC2A* | 1.23E-02 | -1.316 | -0.024 | -0.276 |
| ENSG00000128512 | *DOCK4* | 9.31E-14 | 1.097 | 0.049 | 0.173 |
| ENSG00000136048 | *DRAM1* | 1.54E-13 | 1.369 | 0.399 | 0.733 |
| ENSG00000136982 | *DSCC1* | 5.17E-07 | -1.138 | -0.051 | -0.384 |
| ENSG00000134769 | *DTNA* | 7.01E-06 | 1.080 | 0.001 | 0.265 |
| ENSG00000120129 | *DUSP1* | 3.91E-16 | 1.671 | 0.317 | 0.870 |
| ENSG00000138166 | *DUSP5* | 6.11E-12 | 1.098 | 0.071 | 0.579 |
| ENSG00000139318 | *DUSP6* | 5.52E-17 | 1.719 | 0.501 | 0.919 |
| ENSG00000184545 | *DUSP8* | 6.49E-21 | 1.645 | 0.630 | 0.957 |
| ENSG00000158560 | *DYNC1I1* | 2.02E-11 | -1.193 | -0.150 | -0.651 |
| ENSG00000122547 | *EEPD1* | 1.16E-07 | -1.108 | -0.292 | -0.355 |

| | | | | | |
|---|---|---|---|---|---|
| ENSG00000163576 | *EFHB* | 7.43E-03 | -1.231 | -0.025 | -0.209 |
| ENSG00000115468 | *EFHD1* | 4.00E-17 | -1.699 | -0.257 | -0.857 |
| ENSG00000129521 | *EGLN3* | 2.00E-05 | 1.583 | 0.022 | 0.383 |
| ENSG00000013016 | *EHD3* | 1.27E-05 | -1.025 | -0.054 | -0.676 |
| ENSG00000163435 | *ELF3* | 4.95E-12 | 3.004 | 0.973 | 1.601 |
| ENSG00000128886 | *ELL3* | 9.64E-05 | -1.023 | -0.042 | -0.688 |
| ENSG00000197977 | *ELOVL2* | 1.21E-10 | 2.657 | 1.254 | 2.106 |
| ENSG00000164181 | *ELOVL7* | 1.08E-03 | 1.891 | 0.006 | 0.054 |
| ENSG00000168913 | *ENHO* | 1.13E-02 | -2.315 | -0.003 | -0.034 |
| ENSG00000198018 | *ENTPD7* | 7.13E-20 | 1.620 | 0.290 | 0.682 |
| ENSG00000188833 | *ENTPD8* | 2.37E-02 | -2.824 | -0.003 | -0.051 |
| ENSG00000082397 | *EPB41L3* | 7.16E-18 | 1.390 | 0.014 | 0.118 |
| ENSG00000044524 | *EPHA3* | 3.68E-04 | -1.334 | -0.028 | -0.795 |
| ENSG00000145242 | *EPHA5* | 3.96E-02 | -1.025 | -0.009 | -0.059 |
| ENSG00000182580 | *EPHB3* | 1.21E-13 | -1.359 | -0.224 | -0.607 |
| ENSG00000120915 | *EPHX2* | 6.41E-08 | -1.096 | -0.242 | -0.595 |
| ENSG00000124882 | *EREG* | 1.59E-02 | 1.064 | 0.014 | 0.172 |
| ENSG00000178752 | *ERFE* | 8.48E-32 | 2.633 | 0.878 | 1.763 |
| ENSG00000178607 | *ERN1* | 2.85E-15 | 1.294 | 0.067 | 0.542 |
| ENSG00000116285 | *ERRFI1* | 7.96E-08 | 1.221 | 0.039 | 0.484 |
| ENSG00000164283 | *ESM1* | 5.96E-07 | 1.759 | 0.037 | 0.343 |
| ENSG00000157557 | *ETS2* | 4.64E-13 | 1.180 | 0.136 | 0.569 |
| ENSG00000139083 | *ETV6* | 9.18E-21 | 1.194 | 0.298 | 0.600 |
| ENSG00000158008 | *EXTL1* | 4.27E-02 | -1.109 | -0.016 | -0.081 |
| ENSG00000181104 | *F2R* | 2.26E-16 | -1.916 | -0.065 | -0.713 |
| ENSG00000198734 | *F5* | 9.27E-04 | 1.236 | 0.016 | 0.368 |
| ENSG00000121769 | *FABP3* | 1.08E-20 | -1.331 | -0.307 | -0.499 |
| ENSG00000170323 | *FABP4* | 1.48E-03 | -1.263 | -0.011 | -0.320 |
| ENSG00000169122 | *FAM110B* | 7.92E-09 | 1.083 | 0.069 | 0.561 |
| ENSG00000135842 | *FAM129A* | 1.32E-18 | -1.874 | -0.619 | -1.150 |
| ENSG00000179083 | *FAM133A* | 1.28E-04 | -1.181 | -0.034 | -0.603 |
| ENSG00000109794 | *FAM149A* | 2.06E-06 | -1.280 | -0.032 | -0.607 |
| ENSG00000154319 | *FAM167A* | 7.33E-08 | 1.511 | 0.046 | 0.734 |
| ENSG00000185442 | *FAM174B* | 6.57E-10 | 1.186 | 0.113 | 0.624 |
| ENSG00000197520 | *FAM177B* | 4.55E-28 | 4.310 | 2.489 | 3.184 |
| ENSG00000108950 | *FAM20A* | 1.30E-26 | 2.764 | 1.197 | 1.720 |
| ENSG00000177706 | *FAM20C* | 6.92E-20 | 1.619 | 0.427 | 0.966 |
| ENSG00000185614 | *FAM212A* | 4.71E-07 | -1.540 | -0.060 | -0.557 |
| ENSG00000157470 | *FAM81A* | 4.30E-02 | -1.023 | -0.013 | -0.067 |
| ENSG00000162981 | *FAM84A* | 1.02E-04 | -1.271 | -0.026 | -0.546 |

| | | | | | |
|---|---|---|---|---|---|
| ENSG00000165323 | *FAT3* | 7.84E-09 | -1.293 | -0.090 | -0.837 |
| ENSG00000170271 | *FAXDC2* | 1.38E-09 | -1.342 | -0.350 | -0.796 |
| ENSG00000165140 | *FBP1* | 9.42E-12 | -1.987 | -0.069 | -0.856 |
| ENSG00000141665 | *FBXO15* | 3.04E-03 | -1.709 | -0.018 | -0.176 |
| ENSG00000162894 | *FCMR* | 1.11E-04 | 1.898 | 0.001 | 0.061 |
| ENSG00000113578 | *FGF1* | 8.21E-06 | 1.410 | 0.013 | 0.200 |
| ENSG00000070193 | *FGF10* | 4.31E-24 | 2.227 | 0.697 | 1.249 |
| ENSG00000102466 | *FGF14* | 7.16E-13 | -1.089 | -0.236 | -0.500 |
| ENSG00000140285 | *FGF7* | 4.26E-31 | 1.365 | 0.505 | 0.888 |
| ENSG00000137460 | *FHDC1* | 1.28E-04 | 1.071 | 0.016 | 0.131 |
| ENSG00000022267 | *FHL1* | 2.13E-21 | -1.263 | -0.371 | -0.648 |
| ENSG00000118407 | *FILIP1* | 8.06E-17 | 2.964 | 0.021 | 0.363 |
| ENSG00000204131 | *FLJ44635* | 1.20E-02 | -1.523 | -0.019 | -0.179 |
| ENSG00000075420 | *FNDC3B* | 1.29E-29 | 1.237 | 0.369 | 0.613 |
| ENSG00000143107 | *FNDC7* | 2.80E-02 | -1.636 | -0.015 | -0.035 |
| ENSG00000103241 | *FOXF1* | 7.08E-04 | 2.075 | 0.024 | 1.855 |
| ENSG00000114861 | *FOXP1* | 3.28E-25 | 1.069 | 0.185 | 0.449 |
| ENSG00000165694 | *FRMD7* | 1.32E-02 | -1.121 | -0.026 | -0.305 |
| ENSG00000167996 | *FTH1* | 5.18E-31 | 1.410 | 0.322 | 0.757 |
| ENSG00000010810 | *FYN* | 4.87E-31 | 1.242 | 0.203 | 0.555 |
| ENSG00000180340 | *FZD2* | 7.41E-08 | -1.184 | -0.019 | -0.474 |
| ENSG00000123689 | *G0S2* | 3.45E-30 | 4.313 | 1.375 | 2.400 |
| ENSG00000139112 | *GABARAPL1* | 4.83E-19 | 1.177 | 0.355 | 0.681 |
| ENSG00000143891 | *GALM* | 7.21E-09 | -1.350 | -0.081 | -0.552 |
| ENSG00000131386 | *GALNT15* | 3.92E-28 | 2.351 | 0.886 | 1.441 |
| ENSG00000131979 | *GCH1* | 1.11E-47 | 3.628 | 0.903 | 1.758 |
| ENSG00000130513 | *GDF15* | 2.39E-07 | 1.137 | 0.528 | 0.882 |
| ENSG00000125965 | *GDF5* | 3.78E-19 | -1.339 | -0.349 | -0.615 |
| ENSG00000156466 | *GDF6* | 3.71E-12 | 1.012 | 0.095 | 0.398 |
| ENSG00000143869 | *GDF7* | 2.40E-05 | -1.355 | -0.009 | -0.583 |
| ENSG00000158555 | *GDPD5* | 4.68E-08 | -1.045 | -0.079 | -0.435 |
| ENSG00000131459 | *GFPT2* | 7.90E-36 | 1.887 | 0.514 | 0.997 |
| ENSG00000168546 | *GFRA2* | 1.59E-02 | 1.679 | 0.004 | 0.066 |
| ENSG00000179855 | *GIPC3* | 3.99E-02 | -1.063 | -0.005 | -0.115 |
| ENSG00000198814 | *GK* | 2.40E-04 | 1.148 | 0.035 | 0.274 |
| ENSG00000074047 | *GLI2* | 8.46E-09 | -1.453 | -0.045 | -0.931 |
| ENSG00000107249 | *GLIS3* | 4.98E-18 | 1.345 | 0.207 | 0.594 |
| ENSG00000173221 | *GLRX* | 1.29E-19 | 1.729 | 0.382 | 0.950 |
| ENSG00000156689 | *GLYATL2* | 8.77E-03 | -1.236 | -0.007 | -0.217 |
| ENSG00000112312 | *GMNN* | 2.57E-07 | 1.101 | 0.084 | 0.579 |

| | | | | | |
|---|---|---|---|---|---|
| ENSG00000186469 | *GNG2* | 7.01E-08 | -1.149 | -0.041 | -0.661 |
| ENSG00000176533 | *GNG7* | 6.70E-05 | -1.317 | -0.028 | -0.213 |
| ENSG00000155265 | *GOLGA7B* | 6.67E-05 | 1.299 | 0.013 | 0.247 |
| ENSG00000164850 | *GPER1* | 1.47E-12 | -1.042 | -0.225 | -0.465 |
| ENSG00000112293 | *GPLD1* | 4.15E-12 | -1.010 | -0.195 | -0.821 |
| ENSG00000180998 | *GPR137C* | 1.62E-02 | -1.281 | -0.012 | -0.156 |
| ENSG00000158292 | *GPR153* | 3.62E-15 | -1.223 | -0.279 | -0.611 |
| ENSG00000169508 | *GPR183* | 2.41E-07 | 2.841 | 0.018 | 0.947 |
| ENSG00000170075 | *GPR37L1* | 2.68E-09 | 3.043 | 0.010 | 0.106 |
| ENSG00000181656 | *GPR88* | 2.01E-03 | 1.175 | 0.005 | 0.205 |
| ENSG00000013588 | *GPRC5A* | 1.99E-19 | 1.863 | 0.541 | 1.080 |
| ENSG00000115290 | *GRB14* | 4.83E-06 | 1.441 | -0.003 | 0.094 |
| ENSG00000155511 | *GRIA1* | 1.73E-08 | -1.862 | -0.527 | -1.065 |
| ENSG00000147697 | *GSDMC* | 8.53E-03 | 1.049 | 0.005 | -0.034 |
| ENSG00000169181 | *GSG1L* | 1.70E-02 | -3.352 | -0.004 | -0.053 |
| ENSG00000134201 | *GSTM5* | 8.93E-09 | -1.463 | -0.463 | -0.688 |
| ENSG00000164116 | *GUCY1A1* | 8.76E-06 | -1.553 | -0.402 | -0.745 |
| ENSG00000061918 | *GUCY1B1* | 3.26E-06 | -1.210 | -0.022 | -0.388 |
| ENSG00000179240 | *GVQW3* | 7.24E-06 | -1.230 | -0.334 | -0.821 |
| ENSG00000162882 | *HAAO* | 5.59E-05 | -1.156 | -0.067 | -0.161 |
| ENSG00000188921 | *HACD4* | 8.18E-21 | -1.165 | -0.286 | -0.705 |
| ENSG00000170961 | *HAS2* | 1.39E-07 | 1.691 | 0.046 | 0.944 |
| ENSG00000135077 | *HAVCR2* | 2.96E-04 | -1.034 | -0.002 | -0.301 |
| ENSG00000103145 | *HCFC1R1* | 9.27E-11 | -1.200 | -0.343 | -0.450 |
| ENSG00000002746 | *HECW1* | 4.64E-09 | -1.226 | -0.060 | -0.548 |
| ENSG00000135547 | *HEY2* | 6.04E-05 | -1.301 | -0.024 | -0.563 |
| ENSG00000177374 | *HIC1* | 7.88E-10 | -1.406 | -0.061 | -0.499 |
| ENSG00000100644 | *HIF1A* | 1.86E-12 | 1.171 | 0.359 | 0.662 |
| ENSG00000135245 | *HILPDA* | 8.64E-05 | 2.547 | 0.011 | 0.142 |
| ENSG00000184678 | *HIST2H2BE* | 4.19E-13 | 1.125 | 0.120 | 0.496 |
| ENSG00000095951 | *HIVEP1* | 2.00E-18 | 1.139 | 0.089 | 0.336 |
| ENSG00000127124 | *HIVEP3* | 2.10E-21 | 1.695 | 0.575 | 0.701 |
| ENSG00000159399 | *HK2* | 1.80E-06 | 1.444 | 0.030 | 0.614 |
| ENSG00000241935 | *HOGA1* | 5.17E-07 | -1.599 | 0.002 | -0.340 |
| ENSG00000196196 | *HRCT1* | 7.77E-03 | -1.700 | -0.014 | -0.124 |
| ENSG00000117594 | *HSD11B1* | 4.10E-40 | 4.420 | 1.662 | 2.628 |
| ENSG00000004776 | *HSPB6* | 1.62E-21 | -1.112 | -0.435 | -0.533 |
| ENSG00000173641 | *HSPB7* | 1.39E-25 | -1.750 | -0.719 | -1.026 |
| ENSG00000102468 | *HTR2A* | 6.79E-15 | 3.417 | 0.034 | 1.655 |
| ENSG00000169495 | *HTRA4* | 4.30E-04 | 1.547 | 0.015 | 0.212 |

| | | | | | |
|---|---|---|---|---|---|
| ENSG00000090339 | *ICAM1* | 7.71E-13 | 2.080 | 0.027 | 0.781 |
| ENSG00000115738 | *ID2* | 1.16E-26 | 2.417 | 0.885 | 1.336 |
| ENSG00000137331 | *IER3* | 3.41E-02 | 1.358 | 0.011 | 0.066 |
| ENSG00000185885 | *IFITM1* | 7.17E-03 | 1.635 | 0.014 | 0.147 |
| ENSG00000142089 | *IFITM3* | 7.08E-12 | 1.317 | 0.359 | 0.832 |
| ENSG00000017427 | *IGF1* | 7.44E-11 | 3.407 | 0.025 | 1.588 |
| ENSG00000141753 | *IGFBP4* | 3.85E-30 | 1.134 | 0.232 | 0.684 |
| ENSG00000164136 | *IL15* | 3.96E-04 | 1.093 | 0.013 | 0.163 |
| ENSG00000134470 | *IL15RA* | 9.58E-05 | 1.052 | 0.043 | 0.467 |
| ENSG00000115604 | *IL18R1* | 4.15E-10 | 1.238 | 0.452 | 0.585 |
| ENSG00000136689 | *IL1RN* | 2.15E-02 | 1.394 | 0.009 | 0.173 |
| ENSG00000162892 | *IL24* | 5.89E-05 | 3.224 | 0.007 | 0.037 |
| ENSG00000104998 | *IL27RA* | 6.73E-04 | -1.233 | -0.015 | -0.188 |
| ENSG00000164509 | *IL31RA* | 3.25E-12 | -1.511 | -0.079 | -0.912 |
| ENSG00000136696 | *IL36B* | 4.15E-03 | 1.027 | 0.032 | 0.830 |
| ENSG00000136695 | *IL36RN* | 3.73E-02 | 1.626 | 0.005 | 0.073 |
| ENSG00000104951 | *IL4I1* | 7.24E-07 | 2.266 | 0.018 | 0.378 |
| ENSG00000136244 | *IL6* | 6.73E-32 | 7.861 | 0.023 | 3.915 |
| ENSG00000168685 | *IL7R* | 8.14E-06 | 1.089 | 0.033 | 0.196 |
| ENSG00000122641 | *INHBA* | 2.87E-05 | 1.408 | 0.019 | 0.282 |
| ENSG00000188487 | *INSC* | 3.39E-05 | -1.577 | -0.048 | -1.012 |
| ENSG00000186480 | *INSIG1* | 2.70E-16 | 1.471 | 0.062 | 0.597 |
| ENSG00000125629 | *INSIG2* | 4.86E-36 | 1.206 | 0.204 | 0.431 |
| ENSG00000074706 | *IPCEF1* | 1.22E-03 | -1.053 | -0.006 | -1.443 |
| ENSG00000151151 | *IPMK* | 7.23E-09 | 1.161 | 0.062 | 0.505 |
| ENSG00000120645 | *IQSEC3* | 2.90E-03 | -1.067 | -0.013 | -0.188 |
| ENSG00000134070 | *IRAK2* | 2.59E-27 | 1.912 | 0.681 | 1.117 |
| ENSG00000090376 | *IRAK3* | 8.60E-17 | 1.124 | 0.040 | 0.266 |
| ENSG00000137265 | *IRF4* | 1.01E-02 | 1.761 | 0.010 | 0.125 |
| ENSG00000159387 | *IRX6* | 2.49E-05 | -1.550 | -0.046 | -0.602 |
| ENSG00000101230 | *ISM1* | 8.74E-05 | 1.006 | 0.029 | 0.330 |
| ENSG00000005961 | *ITGA2B* | 7.71E-03 | -2.161 | -0.017 | -0.105 |
| ENSG00000115232 | *ITGA4* | 8.51E-10 | -1.026 | -0.287 | -0.634 |
| ENSG00000105855 | *ITGB8* | 2.11E-05 | 1.331 | 0.023 | 0.261 |
| ENSG00000150995 | *ITPR1* | 1.03E-09 | 1.302 | 0.047 | 0.458 |
| ENSG00000148841 | *ITPRIP* | 3.90E-13 | 1.375 | 0.076 | 0.778 |
| ENSG00000096968 | *JAK2* | 1.26E-27 | 1.105 | 0.115 | 0.354 |
| ENSG00000188385 | *JAKMIP3* | 1.32E-04 | -1.235 | -0.031 | -0.255 |
| ENSG00000171223 | *JUNB* | 3.47E-17 | 1.836 | 0.560 | 1.072 |
| ENSG00000158445 | *KCNB1* | 6.22E-07 | -1.892 | -0.031 | -1.148 |

| | | | | | |
|---|---|---|---|---|---|
| ENSG00000102057 | *KCND1* | 2.43E-02 | -1.019 | -0.012 | -0.167 |
| ENSG00000152049 | *KCNE4* | 4.27E-25 | 1.293 | 0.509 | 0.916 |
| ENSG00000143473 | *KCNH1* | 1.01E-14 | 1.594 | 0.047 | 0.791 |
| ENSG0000055118 | *KCNH2* | 1.56E-04 | -1.158 | -0.060 | -0.779 |
| ENSG00000157551 | *KCNJ15* | 2.40E-10 | 1.409 | 0.058 | 0.992 |
| ENSG00000171303 | *KCNK3* | 6.73E-08 | 4.271 | 0.013 | 3.340 |
| ENSG00000104783 | *KCNN4* | 1.69E-07 | -1.192 | -0.037 | -0.500 |
| ENSG00000184156 | *KCNQ3* | 2.03E-21 | 1.923 | 0.667 | 1.085 |
| ENSG00000156486 | *KCNS2* | 1.06E-02 | -1.625 | 0.002 | -0.084 |
| ENSG00000170745 | *KCNS3* | 2.64E-03 | -1.334 | -0.011 | -0.139 |
| ENSG00000180332 | *KCTD4* | 1.56E-10 | -1.845 | -0.057 | -0.703 |
| ENSG00000138030 | *KHK* | 6.01E-03 | -1.226 | -0.019 | -0.067 |
| ENSG00000109265 | *KIAA1211* | 4.58E-05 | -1.109 | -0.054 | -0.664 |
| ENSG00000162849 | *KIF26B* | 2.06E-15 | -2.017 | -0.651 | -0.986 |
| ENSG00000155980 | *KIF5A* | 2.24E-02 | -1.404 | -0.020 | -0.091 |
| ENSG00000157404 | *KIT* | 3.05E-06 | -1.251 | -0.710 | -0.979 |
| ENSG00000155090 | *KLF10* | 8.15E-24 | 1.199 | 0.390 | 0.655 |
| ENSG00000171872 | *KLF17* | 2.34E-08 | 2.206 | 0.023 | 1.025 |
| ENSG00000119138 | *KLF9* | 2.85E-11 | 1.064 | 0.326 | 0.533 |
| ENSG00000162755 | *KLHDC9* | 1.02E-04 | -1.346 | -0.032 | -0.303 |
| ENSG00000171345 | *KRT19* | 1.95E-03 | -1.457 | -0.023 | -0.806 |
| ENSG00000221852 | *KRTAP1-5* | 2.88E-08 | -1.114 | -0.311 | -0.316 |
| ENSG00000174611 | *KY* | 2.53E-11 | -1.899 | -0.774 | -0.936 |
| ENSG00000115919 | *KYNU* | 1.25E-18 | 3.450 | 1.416 | 2.012 |
| ENSG00000101680 | *LAMA1* | 3.38E-07 | 1.001 | 0.156 | 0.542 |
| ENSG00000129988 | *LBP* | 1.14E-17 | 3.270 | 0.022 | 1.339 |
| ENSG00000148346 | *LCN2* | 2.87E-05 | 2.773 | 0.020 | 1.590 |
| ENSG00000136167 | *LCP1* | 2.22E-02 | -1.477 | -0.016 | -0.053 |
| ENSG00000169744 | *LDB2* | 1.17E-15 | -1.817 | -0.449 | -0.950 |
| ENSG00000134333 | *LDHA* | 5.52E-17 | 1.237 | 0.421 | 0.718 |
| ENSG00000166816 | *LDHD* | 1.08E-07 | -1.319 | -0.489 | -0.798 |
| ENSG00000168675 | *LDLRAD4* | 1.26E-15 | -1.662 | -0.704 | -1.117 |
| ENSG00000106003 | *LFNG* | 3.00E-05 | -1.194 | -0.052 | -0.639 |
| ENSG00000153012 | *LGI2* | 3.69E-08 | -1.814 | -0.051 | -0.935 |
| ENSG00000139292 | *LGR5* | 3.59E-07 | -1.049 | -0.295 | -0.585 |
| ENSG00000128342 | *LIF* | 1.03E-07 | 2.250 | 0.011 | 0.180 |
| ENSG00000072163 | *LIMS2* | 1.03E-11 | -1.317 | -0.261 | -0.608 |
| ENSG00000236882 | *LINC01554* | 5.61E-06 | 2.151 | 0.017 | 0.194 |
| ENSG00000169783 | *LINGO1* | 6.64E-05 | -1.357 | -0.018 | -0.699 |
| ENSG00000136153 | *LMO7* | 3.43E-15 | -1.261 | -0.189 | -0.647 |

| | | | | | |
|---|---|---|---|---|---|
| ENSG00000163431 | *LMOD1* | 2.98E-06 | -1.512 | -0.040 | -0.702 |
| ENSG00000103485 | *LOC105369247* | 1.49E-03 | -2.095 | -0.002 | -0.063 |
| ENSG00000173535 | *LOC254896* | 8.92E-04 | -1.037 | -0.041 | -0.189 |
| ENSG00000165379 | *LRFN5* | 1.30E-02 | -1.206 | -0.021 | -0.354 |
| ENSG00000128606 | *LRRC17* | 2.49E-09 | -2.721 | -0.037 | -1.610 |
| ENSG00000172731 | *LRRC20* | 3.28E-08 | -1.577 | -0.047 | -0.690 |
| ENSG00000128594 | *LRRC4* | 2.20E-06 | -2.034 | -0.036 | -1.102 |
| ENSG00000146006 | *LRRTM2* | 1.12E-07 | -1.450 | -0.473 | -0.591 |
| ENSG00000130592 | *LSP1* | 2.67E-14 | -1.158 | -0.263 | -0.473 |
| ENSG00000172264 | *MACROD2* | 1.49E-05 | -1.375 | -0.038 | -0.533 |
| ENSG00000185022 | *MAFF* | 2.99E-31 | 1.474 | 0.635 | 0.943 |
| ENSG00000144063 | *MALL* | 9.59E-08 | 1.106 | 0.098 | 0.610 |
| ENSG00000165072 | *MAMDC2* | 7.68E-35 | -2.104 | -0.570 | -1.126 |
| ENSG00000196782 | *MAML3* | 5.86E-04 | -1.231 | -1.017 | -1.138 |
| ENSG00000013619 | *MAMLD1* | 2.35E-20 | -1.770 | -0.748 | -1.382 |
| ENSG00000176909 | *MAMSTR* | 3.04E-04 | -1.423 | -0.035 | -0.290 |
| ENSG00000069535 | *MAOB* | 1.16E-15 | 1.803 | 0.052 | 0.622 |
| ENSG00000197769 | *MAP1LC3C* | 5.86E-13 | -2.132 | -1.052 | -1.715 |
| ENSG00000108984 | *MAP2K6* | 5.48E-08 | -2.502 | -0.827 | -1.438 |
| ENSG00000107968 | *MAP3K8* | 1.85E-188 | 2.425 | 1.551 | 1.929 |
| ENSG00000186868 | *MAPT* | 1.17E-02 | -1.235 | -0.019 | -0.713 |
| ENSG00000186205 | *MARC1* | 8.69E-05 | -1.215 | -0.629 | -0.541 |
| ENSG00000173926 | *MARCH3* | 1.68E-15 | 2.353 | 0.014 | 0.655 |
| ENSG00000144583 | *MARCH4* | 1.52E-03 | -1.606 | -0.018 | -0.704 |
| ENSG00000172197 | *MBOAT1* | 1.29E-10 | -1.210 | -0.092 | -0.868 |
| ENSG00000140563 | *MCTP2* | 1.30E-06 | 1.065 | 0.021 | 0.217 |
| ENSG00000102802 | *MEDAG* | 4.18E-37 | 2.175 | 0.820 | 1.351 |
| ENSG00000106511 | *MEOX2* | 2.16E-21 | -1.828 | -0.423 | -1.094 |
| ENSG00000188095 | *MESP2* | 3.57E-02 | -1.320 | -0.012 | -0.135 |
| ENSG00000170439 | *METTL7B* | 1.78E-07 | -2.638 | -0.029 | -1.219 |
| ENSG00000168389 | *MFSD2A* | 7.40E-09 | 2.118 | 0.036 | 1.157 |
| ENSG00000174514 | *MFSD4A* | 2.24E-02 | -1.057 | -0.017 | -0.783 |
| ENSG00000108960 | *MMD* | 1.86E-10 | 1.187 | 0.689 | 0.761 |
| ENSG00000196549 | *MME* | 1.59E-10 | 1.361 | 0.047 | 0.474 |
| ENSG00000196611 | *MMP1* | 1.83E-39 | 6.069 | 1.925 | 3.486 |
| ENSG00000166670 | *MMP10* | 1.15E-11 | 2.147 | 0.014 | 0.419 |
| ENSG00000137745 | *MMP13* | 2.54E-35 | 3.161 | 0.037 | 1.131 |
| ENSG00000149968 | *MMP3* | 7.44E-59 | 4.451 | 0.798 | 1.889 |
| ENSG00000120162 | *MOB3B* | 2.94E-04 | -0.715 | -1.202 | -0.217 |
| ENSG00000186732 | *MPPED1* | 5.45E-03 | -1.342 | -0.009 | -0.945 |

| | | | | | |
|---|---|---|---|---|---|
| ENSG00000066382 | *MPPED2* | 4.07E-05 | -1.193 | -0.040 | -0.469 |
| ENSG00000156968 | *MPV17L* | 1.06E-07 | -1.718 | -0.037 | -0.998 |
| ENSG00000135324 | *MRAP2* | 2.12E-15 | -1.915 | -0.644 | -1.070 |
| ENSG00000184350 | *MRGPRE* | 2.08E-02 | -1.190 | -0.007 | -0.170 |
| ENSG00000072952 | *MRVI1* | 3.51E-15 | -1.650 | -0.470 | -1.144 |
| ENSG00000178860 | *MSC* | 8.42E-07 | 1.032 | 0.033 | 0.277 |
| ENSG00000205362 | *MT1A* | 1.09E-02 | 1.320 | 0.017 | 0.057 |
| ENSG00000169715 | *MT1E* | 3.11E-14 | 1.362 | 0.488 | 0.934 |
| ENSG00000198417 | *MT1F* | 2.43E-17 | 2.308 | 0.049 | 1.192 |
| ENSG00000125144 | *MT1G* | 1.83E-09 | 3.682 | 0.024 | 2.092 |
| ENSG00000205364 | *MT1M* | 8.46E-08 | 1.277 | 0.041 | 0.518 |
| ENSG00000187193 | *MT1X* | 1.12E-14 | 2.105 | 0.681 | 1.340 |
| ENSG00000125148 | *MT2A* | 4.18E-47 | 2.423 | 0.980 | 1.507 |
| ENSG00000242114 | *MTFP1* | 5.72E-04 | 1.613 | 0.027 | 0.818 |
| ENSG00000185499 | *MUC1* | 1.09E-06 | 1.112 | 0.348 | 0.610 |
| ENSG00000133055 | *MYBPH* | 2.52E-10 | 1.754 | 0.544 | 1.003 |
| ENSG00000136449 | *MYCBPAP* | 2.90E-04 | -1.234 | -0.052 | -0.666 |
| ENSG00000065534 | *MYLK* | 5.17E-08 | -1.189 | -0.073 | -0.790 |
| ENSG00000128833 | *MYO5C* | 5.30E-03 | -1.058 | -0.023 | -0.329 |
| ENSG00000034971 | *MYOC* | 2.57E-14 | -1.238 | -0.441 | -0.734 |
| ENSG00000105835 | *NAMPT* | 9.42E-20 | 3.291 | 0.037 | 1.523 |
| ENSG00000186462 | *NAP1L2* | 2.21E-05 | -1.184 | -0.475 | -0.503 |
| ENSG00000186310 | *NAP1L3* | 4.37E-07 | -1.375 | -0.038 | -0.687 |
| ENSG00000156006 | *NAT2* | 3.67E-03 | -1.105 | -0.044 | -0.178 |
| ENSG00000067798 | *NAV3* | 8.35E-05 | -1.072 | -0.035 | -0.567 |
| ENSG00000160796 | *NBEAL2* | 3.67E-05 | -1.076 | -0.074 | -0.788 |
| ENSG00000104490 | *NCALD* | 1.84E-05 | -1.977 | -0.028 | -0.519 |
| ENSG00000154654 | *NCAM2* | 3.37E-11 | 2.096 | 0.034 | 0.896 |
| ENSG00000124479 | *NDP* | 7.29E-06 | 1.865 | 0.014 | 0.173 |
| ENSG00000104419 | *NDRG1* | 6.40E-16 | 1.720 | 0.073 | 0.803 |
| ENSG00000165795 | *NDRG2* | 7.29E-06 | -1.302 | -0.021 | -0.584 |
| ENSG00000165030 | *NFIL3* | 3.99E-21 | 1.858 | 0.438 | 1.058 |
| ENSG00000077150 | *NFKB2* | 2.86E-16 | 1.108 | 0.037 | 0.339 |
| ENSG00000100906 | *NFKBIA* | 1.63E-67 | 2.447 | 0.889 | 1.343 |
| ENSG00000144802 | *NFKBIZ* | 2.09E-246 | 3.596 | 2.648 | 2.975 |
| ENSG00000257108 | *NHLRC4* | 2.57E-02 | -1.066 | -0.019 | -0.085 |
| ENSG00000204131 | *NHSL2* | 1.20E-02 | -1.523 | -0.019 | -0.179 |
| ENSG00000165028 | *NIPSNAP3B* | 2.75E-06 | -1.095 | -0.336 | -0.463 |
| ENSG00000140807 | *NKD1* | 4.11E-06 | -1.014 | -0.043 | -0.825 |
| ENSG00000167034 | *NKX3-1* | 8.92E-18 | 1.420 | 0.426 | 0.511 |

| | | | | | |
|---|---|---|---|---|---|
| ENSG00000182261 | *NLRP10* | 4.17E-05 | -2.052 | -0.020 | -0.847 |
| ENSG00000166741 | *NNMT* | 1.43E-42 | 2.245 | 0.990 | 1.587 |
| ENSG00000151014 | *NOCT* | 4.89E-08 | 1.256 | 0.031 | 0.548 |
| ENSG00000167207 | *NOD2* | 2.53E-24 | 4.202 | 0.024 | 2.032 |
| ENSG00000007171 | *NOS2* | 7.49E-36 | 7.417 | 4.309 | 5.152 |
| ENSG00000163072 | *NOSTRIN* | 4.57E-02 | -1.134 | -0.005 | -0.002 |
| ENSG00000136999 | *NOV* | 4.23E-12 | -1.249 | -0.062 | -0.419 |
| ENSG00000139910 | *NOVA1* | 5.31E-09 | 1.037 | 0.118 | 0.593 |
| ENSG00000171246 | *NPTX1* | 1.63E-10 | -1.653 | -0.607 | -0.832 |
| ENSG00000144852 | *NR1I2* | 3.85E-02 | -1.893 | -0.012 | -0.032 |
| ENSG00000151623 | *NR3C2* | 7.96E-16 | -1.322 | -0.309 | -0.643 |
| ENSG00000153234 | *NR4A2* | 2.87E-12 | 1.755 | 0.068 | 0.798 |
| ENSG00000119508 | *NR4A3* | 2.63E-29 | 2.536 | 0.865 | 1.557 |
| ENSG00000134986 | *NREP* | 1.58E-15 | -1.584 | -0.520 | -0.995 |
| ENSG00000118257 | *NRP2* | 9.77E-17 | 1.388 | 0.160 | 0.660 |
| ENSG00000065320 | *NTN1* | 1.89E-13 | 1.528 | 0.386 | 0.982 |
| ENSG00000148053 | *NTRK2* | 4.29E-12 | -1.044 | -0.335 | -0.722 |
| ENSG00000186364 | *NUDT17* | 2.27E-05 | -1.084 | -0.039 | -0.779 |
| ENSG00000182575 | *NXPH3* | 6.62E-09 | -1.428 | -0.416 | -0.977 |
| ENSG00000205978 | *NYNRIN* | 3.89E-08 | -1.090 | -0.192 | -0.597 |
| ENSG00000154358 | *OBSCN* | 1.43E-18 | -1.093 | -0.384 | -0.605 |
| ENSG00000177989 | *ODF3B* | 2.35E-06 | 1.401 | 0.043 | 0.698 |
| ENSG00000105088 | *OLFM2* | 8.91E-14 | -1.224 | -0.255 | -0.441 |
| ENSG00000183801 | *OLFML1* | 2.80E-04 | -1.793 | -0.035 | -0.503 |
| ENSG00000162745 | *OLFML2B* | 1.45E-13 | -1.123 | -0.243 | -0.460 |
| ENSG00000127083 | *OMD* | 3.59E-10 | -1.017 | -0.102 | -0.606 |
| ENSG00000183715 | *OPCML* | 1.39E-02 | -1.587 | -0.018 | -0.102 |
| ENSG00000125510 | *OPRL1* | 4.77E-05 | -1.003 | -0.040 | -0.518 |
| ENSG00000070882 | *OSBPL3* | 2.33E-18 | -1.226 | -0.353 | -0.711 |
| ENSG00000006025 | *OSBPL7* | 2.70E-08 | -1.162 | -0.125 | -0.601 |
| ENSG00000145623 | *OSMR* | 2.37E-40 | 2.286 | 0.869 | 1.354 |
| ENSG00000180914 | *OXTR* | 1.12E-03 | -1.421 | -0.018 | -0.105 |
| ENSG00000099957 | *P2RX6* | 1.85E-06 | -1.437 | -0.037 | -0.787 |
| ENSG00000076641 | *PAG1* | 2.47E-06 | 1.096 | 0.033 | 0.222 |
| ENSG00000243444 | *PALM2* | 6.84E-08 | 1.626 | 0.064 | 1.128 |
| ENSG00000157654 | *PALM2-AKAP2* | 3.63E-03 | 2.753 | 0.006 | 0.068 |
| ENSG00000099260 | *PALMD* | 8.47E-22 | 1.838 | 0.610 | 1.063 |
| ENSG00000162073 | *PAQR4* | 1.16E-04 | -1.095 | -0.013 | -0.213 |
| ENSG00000102981 | *PARD6A* | 3.84E-04 | -1.009 | -0.055 | -0.703 |
| ENSG00000173599 | *PC* | 1.16E-08 | 1.102 | 0.094 | 0.595 |

| | | | | | |
|---|---|---|---|---|---|
| ENSG00000183570 | *PCBP3* | 2.24E-09 | 1.408 | 0.017 | 0.330 |
| ENSG00000150275 | *PCDH15* | 9.74E-03 | 1.621 | 0.010 | 0.034 |
| ENSG00000189184 | *PCDH18* | 1.18E-07 | -1.175 | -0.370 | -0.875 |
| ENSG00000124253 | *PCK1* | 3.93E-04 | 4.552 | 0.006 | 0.072 |
| ENSG00000175426 | *PCSK1* | 3.51E-17 | 1.822 | 0.362 | 0.911 |
| ENSG00000172572 | *PDE3A* | 1.27E-09 | 1.132 | 0.010 | 0.083 |
| ENSG00000184588 | *PDE4B* | 1.22E-61 | 2.876 | 1.289 | 1.816 |
| ENSG00000113448 | *PDE4D* | 1.21E-26 | 2.308 | 0.456 | 1.006 |
| ENSG00000138735 | *PDE5A* | 7.21E-12 | -2.086 | -0.605 | -1.322 |
| ENSG00000170962 | *PDGFD* | 1.10E-10 | -1.129 | -0.119 | -0.568 |
| ENSG00000152256 | *PDK1* | 1.17E-15 | 1.455 | 0.228 | 0.679 |
| ENSG00000162493 | *PDPN* | 2.79E-23 | 1.576 | 0.557 | 1.008 |
| ENSG00000186862 | *PDZD7* | 6.67E-03 | -1.008 | -0.018 | -0.609 |
| ENSG00000162366 | *PDZK1IP1* | 4.96E-09 | 2.736 | 0.026 | 1.388 |
| ENSG00000109272 | *PF4V1* | 1.88E-03 | 2.203 | 0.018 | 1.334 |
| ENSG00000170525 | *PFKFB3* | 1.36E-09 | 1.320 | 0.080 | 0.711 |
| ENSG00000070087 | *PFN2* | 4.22E-20 | -1.150 | -0.289 | -0.579 |
| ENSG00000176732 | *PFN4* | 5.81E-03 | -1.269 | -0.017 | -0.170 |
| ENSG00000119630 | *PGF* | 5.62E-08 | 2.104 | 0.006 | 0.080 |
| ENSG00000102144 | *PGK1* | 7.99E-12 | 1.083 | 0.245 | 0.540 |
| ENSG00000153823 | *PID1* | 4.07E-27 | 1.845 | 0.017 | 0.266 |
| ENSG00000137193 | *PIM1* | 8.32E-19 | 1.371 | 0.588 | 1.037 |
| ENSG00000198355 | *PIM3* | 5.24E-13 | 1.384 | 0.515 | 0.885 |
| ENSG00000154217 | *PITPNC1* | 2.86E-05 | 1.126 | 0.027 | 0.413 |
| ENSG00000166473 | *PKD1L2* | 6.35E-06 | -1.692 | -0.017 | -1.149 |
| ENSG00000165495 | *PKNOX2* | 4.56E-37 | 2.080 | 0.482 | 1.166 |
| ENSG00000138193 | *PLCE1* | 1.01E-16 | -1.461 | -0.450 | -0.944 |
| ENSG00000075651 | *PLD1* | 7.71E-67 | 2.041 | 0.598 | 1.030 |
| ENSG00000021300 | *PLEKHB1* | 1.03E-03 | -1.057 | -0.018 | -0.142 |
| ENSG00000171680 | *PLEKHG5* | 5.12E-08 | -1.025 | -0.234 | -0.379 |
| ENSG00000147872 | *PLIN2* | 2.19E-03 | 1.060 | 0.025 | 0.435 |
| ENSG00000145632 | *PLK2* | 1.70E-08 | 1.122 | 0.024 | 0.201 |
| ENSG00000152952 | *PLOD2* | 2.87E-12 | 1.370 | 0.249 | 0.620 |
| ENSG00000120756 | *PLS1* | 5.92E-04 | -1.200 | -0.036 | -0.442 |
| ENSG00000188313 | *PLSCR1* | 3.32E-03 | 1.213 | 0.005 | 0.073 |
| ENSG00000221866 | *PLXNA4* | 4.69E-09 | 3.372 | 1.266 | 1.980 |
| ENSG00000141682 | *PMAIP1* | 1.81E-10 | 1.965 | 0.979 | 1.539 |
| ENSG00000182013 | *PNMA8A* | 6.63E-06 | -1.479 | -0.030 | -0.915 |
| ENSG00000204851 | *PNMA8B* | 1.14E-04 | -1.291 | -0.042 | -0.446 |
| ENSG00000130653 | *PNPLA7* | 1.23E-08 | -1.385 | -0.092 | -0.712 |

| | | | | | |
|---|---|---|---|---|---|
| ENSG00000146278 | *PNRC1* | 4.82E-06 | 1.054 | 0.025 | 0.273 |
| ENSG00000028277 | *POU2F2* | 1.15E-29 | 2.726 | 1.458 | 1.909 |
| ENSG00000109819 | *PPARGC1A* | 2.83E-03 | 1.321 | 0.027 | 0.644 |
| ENSG00000110841 | *PPFIBP1* | 2.21E-19 | 1.046 | 0.176 | 0.470 |
| ENSG00000108179 | *PPIF* | 5.14E-16 | 1.278 | 0.395 | 0.791 |
| ENSG00000156475 | *PPP2R2B* | 6.70E-11 | -2.243 | -0.045 | -1.067 |
| ENSG00000214140 | *PRCD* | 1.96E-02 | -1.622 | -0.013 | -0.106 |
| ENSG00000057657 | *PRDM1* | 2.00E-04 | 1.887 | 0.010 | 0.099 |
| ENSG00000186652 | *PRG2* | 1.81E-02 | 1.104 | 0.032 | 0.516 |
| ENSG00000139174 | *PRICKLE1* | 2.64E-10 | -1.465 | -0.455 | -0.998 |
| ENSG00000146143 | *PRIM2* | 1.14E-08 | -1.246 | -0.077 | -0.616 |
| ENSG00000175785 | *PRIMA1* | 1.11E-03 | -1.537 | -0.006 | -0.280 |
| ENSG00000005249 | *PRKAR2B* | 7.90E-23 | 1.903 | 0.063 | 0.994 |
| ENSG00000138669 | *PRKG2* | 5.05E-04 | -1.822 | -0.028 | -0.314 |
| ENSG00000135406 | *PRPH* | 1.71E-06 | -1.918 | -0.037 | -0.518 |
| ENSG00000167371 | *PRRT2* | 1.99E-12 | -1.247 | -0.409 | -0.676 |
| ENSG00000146250 | *PRSS35* | 1.28E-11 | -2.521 | -0.046 | -1.041 |
| ENSG00000125637 | *PSD4* | 6.35E-08 | 1.226 | 0.065 | 0.595 |
| ENSG00000117425 | *PTCH2* | 2.59E-09 | -1.434 | -0.073 | -0.783 |
| ENSG00000148344 | *PTGES* | 6.11E-14 | 2.616 | 0.022 | 0.466 |
| ENSG00000122420 | *PTGFR* | 1.91E-45 | 2.022 | 0.871 | 1.315 |
| ENSG00000073756 | *PTGS2* | 4.17E-12 | 3.153 | 0.016 | 0.546 |
| ENSG00000087494 | *PTHLH* | 9.10E-07 | 1.282 | 0.561 | 1.080 |
| ENSG00000163629 | *PTPN13* | 2.79E-14 | -1.131 | -0.366 | -0.767 |
| ENSG00000163661 | *PTX3* | 4.05E-04 | 1.148 | 0.038 | 0.566 |
| ENSG00000068976 | *PYGM* | 4.76E-03 | -1.517 | -0.025 | -0.396 |
| ENSG00000103485 | *QPRT* | 1.49E-03 | -2.095 | -0.002 | -0.063 |
| ENSG00000139832 | *RAB20* | 2.62E-04 | 1.252 | 0.032 | 0.677 |
| ENSG00000167964 | *RAB26* | 9.84E-06 | -1.505 | -0.034 | -0.465 |
| ENSG00000041353 | *RAB27B* | 1.02E-25 | 2.460 | 1.060 | 1.576 |
| ENSG00000172794 | *RAB37* | 1.75E-04 | -1.168 | -0.035 | -0.530 |
| ENSG00000169750 | *RAC3* | 5.60E-10 | -1.220 | -0.057 | -0.306 |
| ENSG00000166349 | *RAG1* | 1.73E-04 | -1.221 | -0.018 | -0.840 |
| ENSG00000164188 | *RANBP3L* | 2.29E-02 | -1.006 | 0.001 | -0.093 |
| ENSG00000108551 | *RASD1* | 2.39E-10 | 2.710 | 0.024 | 1.313 |
| ENSG00000103710 | *RASL12* | 1.29E-05 | -1.542 | -0.038 | -0.967 |
| ENSG00000163694 | *RBM47* | 3.45E-04 | 1.122 | 0.037 | 0.494 |
| ENSG00000159200 | *RCAN1* | 2.03E-07 | 1.210 | 0.024 | 0.337 |
| ENSG00000172348 | *RCAN2* | 7.86E-18 | -3.059 | -0.077 | -1.624 |
| ENSG00000167771 | *RCOR2* | 5.78E-05 | -1.537 | -0.033 | -0.566 |

| | | | | | |
|---|---|---|---|---|---|
| ENSG00000104856 | *RELB* | 2.20E-22 | 1.716 | 0.334 | 0.760 |
| ENSG00000054967 | *RELT* | 3.67E-06 | 1.316 | 0.022 | 0.982 |
| ENSG00000088320 | *REM1* | 1.07E-04 | -1.040 | -0.037 | -0.306 |
| ENSG00000174236 | *REP15* | 1.99E-04 | 1.104 | 0.039 | 0.467 |
| ENSG00000154153 | *RETREG1* | 1.18E-17 | 1.586 | 0.356 | 0.817 |
| ENSG00000009413 | *REV3L* | 2.18E-17 | 1.223 | 0.154 | 0.477 |
| ENSG00000178882 | *RFLNA* | 2.75E-04 | -1.472 | -0.038 | -0.755 |
| ENSG00000131378 | *RFTN1* | 1.55E-18 | -1.043 | -0.387 | -0.651 |
| ENSG00000143333 | *RGS16* | 2.21E-13 | 1.196 | 0.341 | 0.625 |
| ENSG00000132554 | *RGS22* | 1.49E-02 | -1.142 | -0.024 | -0.206 |
| ENSG00000108370 | *RGS9* | 1.58E-02 | -1.429 | -0.015 | -0.133 |
| ENSG00000072422 | *RHOBTB1* | 3.65E-17 | -1.372 | -0.532 | -0.866 |
| ENSG00000111913 | *RIPOR2* | 4.17E-06 | 1.034 | 0.020 | 0.222 |
| ENSG00000042062 | *RIPOR3* | 6.20E-07 | 1.573 | 0.014 | 0.670 |
| ENSG00000108830 | *RND2* | 9.52E-04 | -1.250 | -0.020 | -0.157 |
| ENSG00000101695 | *RNF125* | 1.10E-03 | -1.462 | -0.019 | -0.917 |
| ENSG00000137393 | *RNF144B* | 2.03E-10 | 1.068 | 0.008 | 0.124 |
| ENSG00000108375 | *RNF43* | 3.04E-05 | -1.388 | -0.057 | -0.583 |
| ENSG00000185008 | *ROBO2* | 1.81E-07 | -0.969 | -0.279 | -1.007 |
| ENSG00000169071 | *ROR2* | 4.34E-06 | 1.114 | 0.086 | 0.511 |
| ENSG00000143365 | *RORC* | 2.25E-02 | -1.433 | -0.015 | -0.133 |
| ENSG00000156313 | *RPGR* | 2.81E-09 | 1.026 | 0.271 | 0.686 |
| ENSG00000215853 | *RPTN* | 1.48E-02 | -1.545 | -0.016 | -0.157 |
| ENSG00000166592 | *RRAD* | 1.48E-05 | 1.292 | 0.033 | 0.589 |
| ENSG00000146374 | *RSPO3* | 4.14E-20 | 1.701 | 0.633 | 1.056 |
| ENSG00000179300 | *RTL3* | 1.32E-04 | -1.032 | -0.040 | -0.438 |
| ENSG00000242732 | *RTL5* | 3.00E-17 | -1.066 | -0.425 | -0.529 |
| ENSG00000040608 | *RTN4R* | 4.82E-02 | -1.059 | 0.000 | -0.070 |
| ENSG00000186907 | *RTN4RL2* | 3.21E-09 | 2.475 | 0.029 | 1.593 |
| ENSG00000171509 | *RXFP1* | 2.70E-03 | 1.111 | 0.030 | 0.109 |
| ENSG00000170989 | *S1PR1* | 5.56E-11 | 1.886 | 0.537 | 1.197 |
| ENSG00000213694 | *S1PR3* | 1.23E-05 | 1.144 | 0.030 | 0.454 |
| ENSG00000173432 | *SAA1* | 2.81E-28 | 6.628 | 0.033 | 4.583 |
| ENSG00000134339 | *SAA2* | 1.63E-17 | 6.156 | 0.030 | 4.201 |
| ENSG00000177570 | *SAMD12* | 8.30E-06 | -1.894 | -0.012 | -0.956 |
| ENSG00000203943 | *SAMD13* | 5.60E-04 | -1.538 | -0.024 | -0.684 |
| ENSG00000164483 | *SAMD3* | 1.34E-02 | -1.185 | -0.008 | -0.171 |
| ENSG00000203727 | *SAMD5* | 8.65E-15 | -1.299 | -0.267 | -0.728 |
| ENSG00000123453 | *SARDH* | 6.98E-05 | -1.678 | -0.026 | -0.550 |
| ENSG00000130066 | *SAT1* | 1.03E-30 | 1.594 | 0.276 | 0.648 |

| | | | | | |
|---|---|---|---|---|---|
| ENSG00000064932 | *SBNO2* | 7.65E-27 | 1.791 | 0.783 | 1.247 |
| ENSG00000073060 | *SCARB1* | 3.38E-12 | 1.087 | 0.370 | 0.685 |
| ENSG00000184178 | *SCFD2* | 2.13E-10 | -1.104 | -0.409 | -0.653 |
| ENSG00000171951 | *SCG2* | 3.17E-06 | 1.217 | 0.428 | 0.888 |
| ENSG00000161929 | *SCIMP* | 6.46E-04 | -1.006 | -0.013 | -0.521 |
| ENSG00000006747 | *SCIN* | 1.97E-06 | -1.206 | -0.041 | -0.542 |
| ENSG00000136531 | *SCN2A* | 6.29E-03 | -1.399 | -0.017 | -0.340 |
| ENSG00000149575 | *SCN2B* | 1.97E-08 | -1.536 | -0.050 | -0.568 |
| ENSG00000153253 | *SCN3A* | 5.28E-03 | -1.499 | -0.010 | -0.838 |
| ENSG00000103184 | *SEC14L5* | 4.45E-04 | -1.255 | -0.591 | -0.829 |
| ENSG00000120341 | *SEC16B* | 3.21E-05 | -1.076 | -0.035 | -0.696 |
| ENSG00000174175 | *SELP* | 3.55E-04 | 1.944 | 0.009 | 0.148 |
| ENSG00000170381 | *SEMA3E* | 1.31E-14 | -1.120 | -0.188 | -0.596 |
| ENSG00000137872 | *SEMA6D* | 1.75E-05 | 1.014 | 0.018 | 0.174 |
| ENSG00000108387 | *SEPT4* | 1.86E-05 | -1.468 | -0.036 | -0.658 |
| ENSG00000135919 | *SERPINE2* | 4.30E-11 | 1.381 | 0.056 | 0.539 |
| ENSG00000198879 | *SFMBT2* | 1.84E-08 | 1.439 | 0.004 | 0.195 |
| ENSG00000104332 | *SFRP1* | 8.13E-11 | 2.622 | 0.031 | 1.204 |
| ENSG00000167037 | *SGSM1* | 5.88E-04 | -1.524 | -0.013 | -0.112 |
| ENSG00000183918 | *SH2D1A* | 1.57E-02 | -1.499 | -0.011 | -0.103 |
| ENSG00000169247 | *SH3TC2* | 4.95E-03 | -1.051 | -0.020 | -0.676 |
| ENSG00000162105 | *SHANK2* | 1.54E-05 | -1.122 | -0.057 | -0.680 |
| ENSG00000107338 | *SHB* | 1.68E-17 | 1.951 | 0.422 | 1.116 |
| ENSG00000185634 | *SHC4* | 7.72E-07 | -1.502 | -0.477 | -0.702 |
| ENSG00000138771 | *SHROOM3* | 2.40E-13 | -1.768 | -0.622 | -1.255 |
| ENSG00000197046 | *SIGLEC15* | 4.55E-03 | -1.345 | -0.019 | -0.930 |
| ENSG00000142178 | *SIK1* | 4.87E-16 | 2.314 | 0.038 | 1.065 |
| ENSG00000198053 | *SIRPA* | 3.08E-34 | 1.125 | 0.116 | 0.409 |
| ENSG00000101307 | *SIRPB1* | 1.20E-04 | 1.119 | 0.031 | 0.163 |
| ENSG00000110911 | *SLC11A2* | 1.21E-34 | 1.278 | 0.495 | 0.731 |
| ENSG00000141469 | *SLC14A1* | 2.60E-16 | -3.065 | -1.073 | -1.855 |
| ENSG00000112394 | *SLC16A10* | 9.05E-07 | 3.011 | 0.003 | 0.089 |
| ENSG00000174326 | *SLC16A11* | 2.40E-02 | -1.070 | -0.016 | 0.011 |
| ENSG00000163053 | *SLC16A14* | 2.86E-06 | -1.387 | -0.033 | -0.592 |
| ENSG00000141526 | *SLC16A3* | 2.02E-11 | 1.187 | 0.253 | 0.723 |
| ENSG00000108932 | *SLC16A6* | 4.90E-02 | 1.236 | 0.004 | 0.011 |
| ENSG00000118596 | *SLC16A7* | 2.84E-12 | 1.629 | 0.059 | 0.591 |
| ENSG00000165449 | *SLC16A9* | 1.05E-05 | -1.593 | -0.041 | -0.436 |
| ENSG00000146409 | *SLC18B1* | 1.41E-13 | 1.132 | 0.097 | 0.451 |
| ENSG00000117479 | *SLC19A2* | 1.26E-12 | 1.334 | 0.069 | 0.654 |

| | | | | | |
|---|---|---|---|---|---|
| ENSG00000135917 | *SLC19A3* | 1.32E-43 | 3.422 | 1.244 | 2.043 |
| ENSG00000112499 | *SLC22A2* | 7.10E-03 | 1.944 | 0.003 | 0.024 |
| ENSG00000137266 | *SLC22A23* | 9.23E-20 | 1.576 | 0.620 | 0.964 |
| ENSG00000258708 | *SLC25A21-AS1* | 6.55E-03 | -1.002 | -0.031 | -0.357 |
| ENSG00000155287 | *SLC25A28* | 1.62E-21 | 1.803 | 0.185 | 0.818 |
| ENSG00000147454 | *SLC25A37* | 1.43E-11 | 1.069 | 0.040 | 0.306 |
| ENSG00000091137 | *SLC26A4* | 3.33E-07 | -1.623 | -0.046 | -0.932 |
| ENSG00000174502 | *SLC26A9* | 2.87E-05 | 2.849 | 0.001 | 0.067 |
| ENSG00000143554 | *SLC27A3* | 4.65E-09 | -1.113 | -0.056 | -0.408 |
| ENSG00000117394 | *SLC2A1* | 2.22E-06 | 1.119 | 0.026 | 0.473 |
| ENSG00000146411 | *SLC2A12* | 1.70E-09 | -1.417 | -0.060 | -0.882 |
| ENSG00000059804 | *SLC2A3* | 1.72E-06 | 1.590 | 0.033 | 0.740 |
| ENSG00000142583 | *SLC2A5* | 1.50E-23 | 2.401 | 0.893 | 1.618 |
| ENSG00000170385 | *SLC30A1* | 4.90E-27 | 1.591 | 0.522 | 0.818 |
| ENSG00000139209 | *SLC38A4* | 4.18E-05 | -1.631 | -0.022 | -0.983 |
| ENSG00000104635 | *SLC39A14* | 3.49E-63 | 3.314 | 1.548 | 2.172 |
| ENSG00000138821 | *SLC39A8* | 6.20E-40 | 3.420 | 0.048 | 1.223 |
| ENSG00000138449 | *SLC40A1* | 2.74E-15 | -1.888 | -0.079 | -0.930 |
| ENSG00000167703 | *SLC43A2* | 6.82E-43 | 2.283 | 0.761 | 1.247 |
| ENSG00000134802 | *SLC43A3* | 1.38E-27 | 1.441 | 0.406 | 0.840 |
| ENSG00000180638 | *SLC47A2* | 3.29E-09 | -1.426 | -0.261 | -0.466 |
| ENSG00000131389 | *SLC6A6* | 5.46E-06 | 1.261 | 0.036 | 0.374 |
| ENSG00000196517 | *SLC6A9* | 1.99E-06 | 1.089 | 0.043 | 0.550 |
| ENSG00000003989 | *SLC7A2* | 5.29E-29 | 3.335 | 0.804 | 1.424 |
| ENSG00000103257 | *SLC7A5* | 4.02E-10 | 1.743 | 0.039 | 0.736 |
| ENSG00000183023 | *SLC8A1* | 1.00E-17 | -1.473 | -0.428 | -0.946 |
| ENSG00000124107 | *SLPI* | 4.44E-10 | 1.593 | 0.042 | 0.885 |
| ENSG00000198732 | *SMOC1* | 2.13E-04 | 1.385 | 0.011 | 0.172 |
| ENSG00000088826 | *SMOX* | 2.44E-10 | 1.584 | 0.449 | 0.837 |
| ENSG00000124216 | *SNAI1* | 5.57E-05 | 1.017 | 0.057 | 0.435 |
| ENSG00000065609 | *SNAP91* | 1.17E-02 | -1.169 | -0.014 | -1.511 |
| ENSG00000023608 | *SNAPC1* | 1.09E-11 | 1.102 | 0.346 | 0.619 |
| ENSG00000145335 | *SNCA* | 7.74E-03 | -1.663 | -0.008 | -0.063 |
| ENSG00000064692 | *SNCAIP* | 1.36E-02 | -1.093 | -0.030 | -0.169 |
| ENSG00000173267 | *SNCG* | 5.00E-11 | -1.374 | -0.092 | -0.485 |
| ENSG00000184557 | *SOCS3* | 2.99E-31 | 1.734 | 0.544 | 1.037 |
| ENSG00000112096 | *SOD2* | 1.02E-44 | 4.185 | 1.970 | 2.777 |
| ENSG00000112096 | *SOD2-OT1* | 1.02E-44 | 4.185 | 1.970 | 2.777 |
| ENSG00000137642 | *SORL1* | 2.25E-06 | -1.024 | -0.244 | -0.817 |
| ENSG00000137098 | *SPAG8* | 1.90E-03 | -1.073 | -0.029 | -0.424 |

| | | | | | |
|---|---|---|---|---|---|
| ENSG00000164266 | *SPINK1* | 2.27E-02 | 1.480 | 0.019 | 1.844 |
| ENSG00000187678 | *SPRY4* | 1.89E-10 | 1.449 | 0.021 | 0.432 |
| ENSG00000137877 | *SPTBN5* | 1.88E-03 | -1.169 | -0.035 | -1.017 |
| ENSG00000196542 | *SPTSSB* | 2.34E-09 | -2.019 | -0.054 | -1.195 |
| ENSG00000122862 | *SRGN* | 7.70E-23 | 1.195 | 0.041 | 0.298 |
| ENSG00000184343 | *SRPK3* | 6.40E-05 | -1.339 | -0.020 | -0.545 |
| ENSG00000117155 | *SSX2IP* | 1.90E-10 | -1.174 | -0.051 | -0.495 |
| ENSG00000144057 | *ST6GAL2* | 6.48E-03 | -1.614 | -0.021 | -0.095 |
| ENSG00000136840 | *ST6GALNAC4* | 1.35E-13 | 1.020 | 0.371 | 0.660 |
| ENSG00000141750 | *STAC2* | 3.87E-08 | 2.019 | 0.017 | 0.660 |
| ENSG00000168610 | *STAT3* | 5.08E-31 | 1.279 | 0.465 | 0.811 |
| ENSG00000159167 | *STC1* | 2.29E-40 | 4.459 | 1.831 | 2.826 |
| ENSG00000164647 | *STEAP1* | 1.18E-45 | 2.596 | 0.880 | 1.521 |
| ENSG00000157214 | *STEAP2* | 4.31E-24 | 1.844 | 0.085 | 0.719 |
| ENSG00000127954 | *STEAP4* | 1.34E-12 | 2.272 | 0.040 | 0.895 |
| ENSG00000165209 | *STRBP* | 7.30E-16 | -1.072 | -0.271 | -0.513 |
| ENSG00000128578 | *STRIP2* | 2.18E-11 | 1.818 | 0.834 | 1.360 |
| ENSG00000135604 | *STX11* | 1.05E-03 | 1.177 | 0.028 | 0.541 |
| ENSG00000130540 | *SULT4A1* | 4.48E-05 | 1.355 | -0.002 | 0.112 |
| ENSG00000078269 | *SYNJ2* | 1.68E-23 | 1.294 | 0.469 | 0.747 |
| ENSG00000182253 | *SYNM* | 1.57E-19 | -1.167 | -0.318 | -0.625 |
| ENSG00000143028 | *SYPL2* | 2.83E-02 | -1.091 | -0.006 | -0.050 |
| ENSG00000019505 | *SYT13* | 1.33E-02 | -1.629 | -0.008 | -0.136 |
| ENSG00000137501 | *SYTL2* | 9.06E-08 | -1.076 | -0.069 | -0.535 |
| ENSG00000165929 | *TC2N* | 2.56E-09 | 2.618 | 0.019 | 1.394 |
| ENSG00000081059 | *TCF7* | 2.28E-11 | -1.165 | -0.069 | -0.407 |
| ENSG00000182134 | *TDRKH* | 1.44E-04 | -1.218 | -0.027 | -0.640 |
| ENSG00000120156 | *TEK* | 8.87E-08 | -1.877 | -0.045 | -1.082 |
| ENSG00000105825 | *TFPI2* | 1.65E-07 | 2.166 | 0.023 | 0.842 |
| ENSG00000072274 | *TFRC* | 3.83E-14 | 1.188 | 0.458 | 0.663 |
| ENSG00000092969 | *TGFB2* | 9.61E-19 | -1.089 | -0.409 | -0.715 |
| ENSG00000106799 | *TGFBR1* | 1.02E-22 | 1.125 | 0.300 | 0.561 |
| ENSG00000177426 | *TGIF1* | 1.38E-11 | 1.042 | 0.259 | 0.596 |
| ENSG00000157150 | *TIMP4* | 5.58E-07 | -1.138 | -0.039 | -0.482 |
| ENSG00000137462 | *TLR2* | 4.35E-06 | 2.375 | 0.009 | 0.104 |
| ENSG00000187554 | *TLR5* | 3.71E-05 | -1.362 | -0.040 | -0.703 |
| ENSG00000166292 | *TMEM100* | 7.77E-11 | 2.097 | 0.771 | 0.944 |
| ENSG00000183160 | *TMEM119* | 1.53E-23 | -2.257 | -0.682 | -1.119 |
| ENSG00000166448 | *TMEM130* | 1.57E-02 | -1.677 | -0.014 | -0.147 |
| ENSG00000157111 | *TMEM171* | 4.35E-13 | -1.651 | -0.516 | -1.198 |

| | | | | | |
|---|---|---|---|---|---|
| ENSG00000172738 | *TMEM217* | 8.14E-06 | 1.647 | 0.025 | 0.838 |
| ENSG00000196932 | *TMEM26* | 9.94E-24 | -2.393 | -0.813 | -1.337 |
| ENSG00000126950 | *TMEM35A* | 1.73E-05 | -2.574 | -0.020 | -0.257 |
| ENSG00000171227 | *TMEM37* | 2.49E-03 | -1.280 | -0.033 | -0.864 |
| ENSG00000147027 | *TMEM47* | 1.94E-34 | -1.079 | -0.395 | -0.728 |
| ENSG00000136842 | *TMOD1* | 7.91E-20 | 2.336 | 0.786 | 1.473 |
| ENSG00000128872 | *TMOD2* | 2.31E-05 | -1.158 | -0.042 | -0.790 |
| ENSG00000041982 | *TNC* | 3.92E-14 | 1.354 | 0.075 | 0.616 |
| ENSG00000185215 | *TNFAIP2* | 1.79E-24 | 1.668 | 0.071 | 0.535 |
| ENSG00000118503 | *TNFAIP3* | 2.62E-50 | 1.978 | 0.262 | 0.675 |
| ENSG00000123610 | *TNFAIP6* | 9.68E-38 | 5.293 | 1.605 | 2.813 |
| ENSG00000145779 | *TNFAIP8* | 6.62E-12 | 1.202 | 0.010 | 0.184 |
| ENSG00000173535 | *TNFRSF10C* | 8.92E-04 | -1.037 | -0.041 | -0.189 |
| ENSG00000164761 | *TNFRSF11B* | 2.72E-15 | 1.711 | 0.580 | 0.971 |
| ENSG00000127863 | *TNFRSF19* | 1.00E-17 | -2.323 | -0.753 | -1.361 |
| ENSG00000028137 | *TNFRSF1B* | 6.99E-21 | 2.501 | 0.464 | 1.185 |
| ENSG00000049249 | *TNFRSF9* | 1.70E-04 | 1.759 | 0.007 | 0.062 |
| ENSG00000121858 | *TNFSF10* | 3.79E-02 | 1.039 | 0.010 | 0.051 |
| ENSG00000120659 | *TNFSF11* | 5.25E-04 | 2.614 | 0.010 | 0.123 |
| ENSG00000154310 | *TNIK* | 2.64E-14 | 1.084 | 0.219 | 0.328 |
| ENSG00000131746 | *TNS4* | 1.18E-05 | 1.562 | 0.028 | 0.695 |
| ENSG00000198846 | *TOX* | 1.00E-12 | -2.133 | -0.060 | -1.047 |
| ENSG00000078804 | *TP53INP2* | 3.82E-25 | 1.166 | 0.216 | 0.569 |
| ENSG00000111669 | *TPI1* | 1.53E-12 | 1.024 | 0.260 | 0.588 |
| ENSG00000056972 | *TRAF3IP2* | 1.81E-17 | 1.010 | 0.325 | 0.637 |
| ENSG00000124731 | *TREM1* | 3.71E-09 | 3.986 | 0.010 | 1.826 |
| ENSG00000173334 | *TRIB1* | 9.21E-19 | 1.772 | 0.837 | 1.302 |
| ENSG00000121060 | *TRIM25* | 2.96E-06 | 1.020 | 0.008 | 0.157 |
| ENSG00000134253 | *TRIM45* | 1.69E-07 | -1.413 | -0.046 | -0.880 |
| ENSG00000158022 | *TRIM63* | 3.45E-02 | 1.631 | 0.016 | 0.133 |
| ENSG00000137672 | *TRPC6* | 2.99E-06 | -1.897 | -0.025 | -0.760 |
| ENSG00000157514 | *TSC22D3* | 2.73E-24 | -1.014 | -0.255 | -0.374 |
| ENSG00000099282 | *TSPAN15* | 1.58E-04 | -1.463 | -0.032 | -0.238 |
| ENSG00000157570 | *TSPAN18* | 6.47E-07 | -1.570 | -0.036 | -0.681 |
| ENSG00000064201 | *TSPAN32* | 2.54E-07 | -1.578 | -0.026 | -0.736 |
| ENSG00000122691 | *TWIST1* | 2.52E-17 | 1.387 | 0.456 | 0.825 |
| ENSG00000025708 | *TYMP* | 3.81E-11 | 2.102 | 0.046 | 1.054 |
| ENSG00000145040 | *UCN2* | 5.69E-05 | 1.270 | -0.003 | 0.000 |
| ENSG00000112715 | *VEGFA* | 2.62E-08 | 1.960 | 0.030 | 0.796 |
| ENSG00000112299 | *VNN1* | 1.20E-31 | 3.323 | 0.048 | 1.656 |

| | | | | | |
|---|---|---|---|---|---|
| ENSG00000112303 | *VNN2* | 1.89E-02 | 1.555 | 0.003 | 0.056 |
| ENSG00000093134 | *VNN3* | 7.11E-26 | 5.096 | 0.014 | 2.715 |
| ENSG00000214376 | *VSTM5* | 5.62E-04 | -2.034 | -0.008 | -0.722 |
| ENSG00000188730 | *VWC2* | 4.69E-04 | 1.596 | 0.024 | 1.410 |
| ENSG00000166272 | *WBP1L* | 5.64E-24 | 1.051 | 0.272 | 0.627 |
| ENSG00000162643 | *WDR63* | 8.08E-04 | -1.140 | -0.036 | -0.504 |
| ENSG00000187260 | *WDR86* | 1.46E-06 | 1.905 | 0.005 | 0.236 |
| ENSG00000103175 | *WFDC1* | 5.21E-07 | -1.376 | -0.439 | -0.887 |
| ENSG00000070540 | *WIPI1* | 4.81E-26 | 1.542 | 0.497 | 0.888 |
| ENSG00000104415 | *WISP1* | 4.73E-20 | 1.562 | 0.052 | 0.488 |
| ENSG00000114251 | *WNT5A* | 1.37E-10 | 1.138 | 0.057 | 0.352 |
| ENSG00000075035 | *WSCD2* | 8.32E-19 | -1.953 | -0.649 | -1.198 |
| ENSG00000146457 | *WTAP* | 1.64E-42 | 2.368 | 0.830 | 1.355 |
| ENSG00000047597 | *XK* | 8.54E-03 | -0.855 | -0.025 | -1.347 |
| ENSG00000103489 | *XYLT1* | 5.83E-11 | 1.129 | 0.055 | 0.439 |
| ENSG00000166793 | *YPEL4* | 4.03E-06 | -1.011 | -0.102 | -0.356 |
| ENSG00000163874 | *ZC3H12A* | 5.33E-56 | 3.384 | 1.932 | 2.424 |
| ENSG00000066185 | *ZMYND12* | 1.60E-05 | -1.369 | -0.011 | -0.202 |
| ENSG00000168661 | *ZNF30* | 1.15E-04 | -1.093 | -0.038 | -0.232 |
| ENSG00000161642 | *ZNF385A* | 7.29E-07 | 1.101 | 0.107 | 0.644 |
| ENSG00000144331 | *ZNF385B* | 1.09E-04 | -1.022 | -0.019 | -0.233 |
| ENSG00000175322 | *ZNF519* | 7.13E-05 | -1.299 | -0.038 | -0.375 |
| ENSG00000176472 | *ZNF575* | 1.39E-05 | -1.149 | -0.033 | -0.439 |
| ENSG00000178882 | *ZNF664-RFLNA* | 2.75E-04 | -1.472 | -0.038 | -0.755 |
| ENSG00000143067 | *ZNF697* | 5.77E-09 | 1.141 | 0.063 | 0.670 |
| ENSG00000164684 | *ZNF704* | 7.74E-11 | -1.126 | -0.187 | -0.626 |
| ENSG00000197385 | *ZNF860* | 2.75E-07 | -1.183 | -0.024 | -0.839 |
| ENSG00000132003 | *ZSWIM4* | 9.33E-10 | 1.152 | 0.078 | 0.566 |

**Supplementary Table 5.** Overview of differentially expressed genes in end-stage osteoarthritis-derived synovial fibroblasts between vehicle control and IL-17 treatment (10 ng/ml) with an adjusted p-value of <0.05. Green boxes note logarithmic fold changes (LFC) of at least +1 and red boxes note LFC of at least -1, while blank boxes show an LFC -1 and 1. Genes are sorted alphabetically on SYMBOL gene name.

| **Ensemble gene name** | **SYMBOL gene name** | **Adjusted p-value** | **Log2 Fold Change (LFC)** | | |
|---|---|---|---|---|---|
| | | | IL-17A | IL-17F | IL-17AF |
| ENSG00000198691 | *ABCA4* | 1.41E-02 | 1.608 | 0.017 | 0.043 |
| ENSG00000129048 | *ACKR4* | 5.44E-07 | -1.132 | -0.024 | -0.075 |
| ENSG00000068366 | *ACSL4* | 9.93E-17 | 1.072 | 0.213 | 0.352 |
| ENSG00000159251 | *ACTC1* | 1.15E-08 | -1.709 | -0.823 | -0.968 |
| ENSG00000138316 | *ADAMTS14* | 1.06E-11 | -1.173 | -0.374 | -0.541 |
| ENSG00000158859 | *ADAMTS4* | 9.61E-03 | 1.363 | 0.009 | 0.040 |
| ENSG00000111452 | *ADGRD1* | 2.21E-11 | 1.993 | 0.041 | 0.900 |
| ENSG00000150594 | *ADRA2A* | 6.37E-04 | -1.397 | -0.017 | -0.053 |
| ENSG00000111863 | *ADTRP* | 5.40E-04 | 1.027 | 0.047 | 0.505 |
| ENSG00000042286 | *AIFM2* | 2.05E-16 | 1.192 | 0.445 | 0.637 |
| ENSG00000154027 | *AK5* | 3.95E-03 | -1.120 | -0.032 | -0.074 |
| ENSG00000171819 | *ANGPTL7* | 2.52E-04 | -2.096 | -0.020 | -0.068 |
| ENSG00000151150 | *ANK3* | 1.35E-06 | -1.430 | -0.020 | -0.055 |
| ENSG00000154856 | *APCDD1* | 1.24E-03 | 1.100 | 0.023 | 0.496 |
| ENSG00000171388 | *APLN* | 6.14E-08 | 1.679 | 0.684 | 1.079 |
| ENSG00000221963 | *APOL6* | 9.66E-10 | 1.078 | 0.555 | 0.682 |
| ENSG00000165272 | *AQP3* | 2.16E-10 | -1.920 | -0.951 | -1.104 |
| ENSG00000165269 | *AQP7* | 8.36E-07 | -1.958 | -0.030 | -0.786 |
| ENSG00000103569 | *AQP9* | 2.07E-10 | 3.261 | 0.018 | 1.766 |
| ENSG00000106819 | *ASPN* | 6.59E-05 | -1.163 | -0.390 | -0.592 |
| ENSG00000018625 | *ATP1A2* | 1.75E-02 | -1.055 | -0.024 | -0.065 |
| ENSG00000104043 | *ATP8B4* | 3.24E-04 | 1.082 | 0.416 | 0.666 |
| ENSG00000140379 | *BCL2A1* | 2.38E-09 | 2.812 | 0.033 | 1.549 |
| ENSG00000069399 | *BCL3* | 1.61E-38 | 1.728 | 0.849 | 1.037 |
| ENSG00000100739 | *BDKRB1* | 5.09E-05 | 1.169 | 0.021 | 0.066 |
| ENSG00000023445 | *BIRC3* | 1.25E-08 | 1.167 | 0.009 | 0.048 |
| ENSG00000153162 | *BMP6* | 1.28E-06 | 1.711 | 0.024 | 0.101 |
| ENSG00000166920 | *C15orf48* | 2.19E-25 | 4.851 | 0.024 | 2.328 |
| ENSG00000145861 | *C1QTNF2* | 2.92E-08 | -1.767 | -0.036 | -0.322 |
| ENSG00000163145 | *C1QTNF7* | 2.41E-03 | -1.068 | 0.006 | -0.004 |
| ENSG00000139178 | *C1RL* | 7.97E-26 | 1.331 | 0.943 | 1.039 |
| ENSG00000198535 | *C2CD4A* | 8.65E-12 | 2.015 | 0.037 | 0.742 |

| | | | | | |
|---|---|---|---|---|---|
| ENSG00000205502 | *C2CD4B* | 2.47E-13 | 2.029 | 0.020 | 1.095 |
| ENSG00000125730 | *C3* | 7.08E-08 | 1.400 | 0.034 | 0.514 |
| ENSG00000134830 | *C5AR2* | 2.21E-02 | 1.194 | 0.007 | 0.030 |
| ENSG00000074410 | *CA12* | 9.22E-12 | 1.017 | 0.324 | 0.450 |
| ENSG00000204682 | *CASC10* | 1.13E-03 | -1.142 | -0.002 | -0.076 |
| ENSG00000143318 | *CASQ1* | 4.27E-03 | -1.458 | -0.014 | 0.003 |
| ENSG00000198624 | *CCDC69* | 4.54E-05 | 1.282 | 0.031 | 0.676 |
| ENSG00000181374 | *CCL13* | 3.19E-03 | 2.551 | 0.004 | 0.022 |
| ENSG00000108691 | *CCL2* | 7.03E-09 | 2.150 | 0.023 | 0.769 |
| ENSG00000115009 | *CCL20* | 3.46E-15 | 6.252 | 0.021 | 0.068 |
| ENSG00000163823 | *CCR1* | 1.93E-03 | 1.096 | 0.016 | 0.097 |
| ENSG00000126353 | *CCR7* | 1.97E-05 | 2.089 | 0.018 | 1.022 |
| ENSG00000004468 | *CD38* | 2.02E-03 | 1.861 | 0.001 | 0.007 |
| ENSG00000121594 | *CD80* | 2.00E-03 | 2.310 | 0.009 | 0.017 |
| ENSG00000163814 | *CDCP1* | 9.80E-14 | 1.655 | 0.592 | 0.903 |
| ENSG00000138395 | *CDK15* | 8.80E-04 | -1.863 | 0.001 | -0.034 |
| ENSG00000221869 | *CEBPD* | 8.01E-19 | 1.888 | 0.792 | 1.063 |
| ENSG00000105792 | *CFAP69* | 1.38E-18 | 1.498 | 0.494 | 0.605 |
| ENSG00000243649 | *CFB* | 8.25E-13 | 1.676 | 0.035 | 0.625 |
| ENSG00000138135 | *CH25H* | 3.82E-03 | 2.043 | 0.004 | 0.030 |
| ENSG00000016391 | *CHDH* | 1.54E-05 | -1.015 | -0.376 | -0.096 |
| ENSG00000064886 | *CHI3L2* | 8.97E-24 | 3.314 | 0.770 | 1.760 |
| ENSG00000159212 | *CLIC6* | 2.02E-06 | 1.648 | 0.029 | 0.866 |
| ENSG00000134326 | *CMPK2* | 1.93E-02 | 1.671 | 0.871 | 0.039 |
| ENSG00000154529 | *CNTNAP3B* | 4.06E-03 | -1.012 | -0.029 | -0.604 |
| ENSG00000100473 | *COCH* | 3.55E-04 | 1.317 | 0.026 | 0.028 |
| ENSG00000204291 | *COL15A1* | 7.51E-06 | 1.019 | 0.382 | 0.583 |
| ENSG00000081052 | *COL4A4* | 2.92E-03 | 1.067 | 0.019 | 0.037 |
| ENSG00000047457 | *CP* | 3.00E-24 | 1.472 | 0.475 | 0.707 |
| ENSG00000108342 | *CSF3* | 8.69E-18 | 8.823 | 0.023 | 5.969 |
| ENSG00000135047 | *CTSL* | 3.03E-30 | 1.443 | 0.659 | 0.865 |
| ENSG00000163131 | *CTSS* | 1.35E-23 | 1.492 | 0.261 | 0.648 |
| ENSG00000163739 | *CXCL1* | 3.52E-43 | 7.132 | 4.338 | 4.983 |
| ENSG00000081041 | *CXCL2* | 1.76E-25 | 4.725 | 2.923 | 3.067 |
| ENSG00000163734 | *CXCL3* | 2.85E-28 | 4.971 | 2.435 | 2.917 |
| ENSG00000163735 | *CXCL5* | 2.70E-14 | 3.685 | 0.018 | 1.398 |
| ENSG00000124875 | *CXCL6* | 2.68E-27 | 6.126 | 3.437 | 4.140 |
| ENSG00000169429 | *CXCL8* | 2.64E-19 | 3.686 | 0.918 | 1.386 |
| ENSG00000137869 | *CYP19A1* | 2.38E-06 | 1.844 | 1.032 | 1.569 |
| ENSG00000172817 | *CYP7B1* | 2.19E-03 | 1.703 | 0.012 | 0.045 |

| | | | | | |
|---|---|---|---|---|---|
| ENSG00000164488 | *DACT2* | 1.80E-04 | 2.179 | 0.019 | 1.661 |
| ENSG00000123977 | *DAW1* | 3.19E-05 | 2.235 | 0.023 | 0.982 |
| ENSG00000164825 | *DEFB1* | 3.48E-02 | 1.159 | 0.009 | 0.029 |
| ENSG00000165507 | *DEPP1* | 2.11E-09 | 1.402 | 0.468 | 0.598 |
| ENSG00000062282 | *DGAT2* | 4.02E-25 | 2.007 | 1.117 | 1.327 |
| ENSG00000211448 | *DIO2* | 7.23E-07 | 1.661 | 0.028 | 0.603 |
| ENSG00000149927 | *DOC2A* | 6.79E-03 | -1.323 | -0.012 | -0.051 |
| ENSG00000102385 | *DRP2* | 8.72E-06 | -1.610 | -0.046 | -0.556 |
| ENSG00000134769 | *DTNA* | 1.24E-12 | 1.162 | 0.430 | 0.546 |
| ENSG00000139318 | *DUSP6* | 5.38E-11 | 1.498 | 0.509 | 0.782 |
| ENSG00000136160 | *EDNRB* | 8.44E-10 | 1.099 | 0.420 | 0.637 |
| ENSG00000164181 | *ELOVL7* | 9.27E-10 | 1.698 | 0.034 | 0.763 |
| ENSG00000198018 | *ENTPD7* | 9.37E-09 | 1.030 | 0.092 | 0.282 |
| ENSG00000133106 | *EPSTI1* | 6.45E-03 | 1.281 | 0.926 | 0.818 |
| ENSG00000124882 | *EREG* | 3.28E-04 | 1.095 | 0.027 | 0.222 |
| ENSG00000178752 | *ERFE* | 2.16E-09 | 1.346 | 0.331 | 0.802 |
| ENSG00000139083 | *ETV6* | 3.02E-11 | 1.014 | 0.474 | 0.633 |
| ENSG00000185862 | *EVI2B* | 3.69E-10 | 1.934 | 0.585 | 1.265 |
| ENSG00000110723 | *EXPH5* | 7.62E-13 | -1.347 | -0.569 | -0.886 |
| ENSG00000181104 | *F2R* | 1.33E-13 | -1.213 | -0.052 | -0.400 |
| ENSG00000121769 | *FABP3* | 7.39E-12 | -1.154 | -0.536 | -0.697 |
| ENSG00000170323 | *FABP4* | 1.11E-03 | -1.406 | -0.028 | -0.058 |
| ENSG00000150510 | *FAM124A* | 2.79E-08 | 1.299 | 0.039 | 0.634 |
| ENSG00000159784 | *FAM131B* | 1.16E-02 | -1.061 | -0.015 | -0.065 |
| ENSG00000147724 | *FAM135B* | 1.68E-03 | -1.137 | -0.031 | -0.079 |
| ENSG00000148541 | *FAM13C* | 1.77E-03 | -1.198 | -0.021 | -0.091 |
| ENSG00000197520 | *FAM177B* | 1.94E-04 | 1.496 | 0.016 | 0.621 |
| ENSG00000204767 | *FAM196B* | 4.71E-03 | 1.046 | 0.436 | 0.398 |
| ENSG00000164125 | *FAM198B* | 4.24E-18 | 1.249 | 0.595 | 0.661 |
| ENSG00000108950 | *FAM20A* | 8.35E-15 | 1.680 | 0.614 | 0.949 |
| ENSG00000185112 | *FAM43A* | 1.50E-05 | -1.107 | -0.041 | -0.370 |
| ENSG00000165323 | *FAT3* | 1.34E-07 | -1.146 | -0.364 | -0.833 |
| ENSG00000146267 | *FAXC* | 1.07E-02 | -1.332 | -0.002 | -0.015 |
| ENSG00000162894 | *FCMR* | 2.36E-03 | 1.275 | -0.005 | 0.020 |
| ENSG00000118407 | *FILIP1* | 4.84E-02 | 1.127 | 0.005 | 0.020 |
| ENSG00000204131 | *FLJ44635* | 1.19E-03 | -1.201 | -0.020 | -0.110 |
| ENSG00000125848 | *FLRT3* | 1.24E-09 | 2.141 | 0.026 | 0.746 |
| ENSG00000138759 | *FRAS1* | 7.46E-05 | 1.587 | 0.019 | 0.986 |
| ENSG00000131386 | *GALNT15* | 1.58E-19 | 1.225 | 0.569 | 0.720 |
| ENSG00000162645 | *GBP2* | 1.19E-05 | 1.262 | 0.025 | 0.618 |

| | | | | | |
|---|---|---|---|---|---|
| ENSG00000131979 | *GCH1* | 4.03E-04 | 1.401 | 0.011 | 0.044 |
| ENSG00000131459 | *GFPT2* | 3.48E-12 | 1.017 | 0.163 | 0.436 |
| ENSG00000168546 | *GFRA2* | 3.87E-02 | 1.130 | 0.017 | 0.062 |
| ENSG00000165474 | *GJB2* | 4.02E-06 | 1.112 | 0.017 | 0.169 |
| ENSG00000107249 | *GLIS3* | 4.46E-16 | 1.251 | 0.353 | 0.499 |
| ENSG00000173221 | *GLRX* | 2.16E-13 | 1.168 | 0.399 | 0.589 |
| ENSG00000127920 | *GNG11* | 5.58E-23 | 1.276 | 0.582 | 0.763 |
| ENSG00000167588 | *GPD1* | 3.10E-07 | -1.705 | -0.031 | -0.121 |
| ENSG00000164850 | *GPER1* | 6.99E-09 | -1.022 | -0.123 | -0.132 |
| ENSG00000158292 | *GPR153* | 4.90E-17 | -1.122 | -0.413 | -0.514 |
| ENSG00000105509 | *HAS1* | 2.72E-06 | 1.649 | 0.016 | 0.081 |
| ENSG00000188290 | *HES4* | 2.65E-04 | 1.072 | 0.033 | 1.028 |
| ENSG00000135245 | *HILPDA* | 4.57E-08 | 1.764 | 0.014 | 0.059 |
| ENSG00000127124 | *HIVEP3* | 1.30E-07 | 1.548 | 0.726 | 0.538 |
| ENSG00000108924 | *HLF* | 2.21E-07 | -1.564 | -0.833 | -1.054 |
| ENSG00000164120 | *HPGD* | 1.14E-05 | -3.144 | -0.019 | -0.049 |
| ENSG00000117594 | *HSD11B1* | 9.16E-15 | 1.219 | 0.466 | 0.628 |
| ENSG00000169271 | *HSPB3* | 1.77E-03 | -1.657 | -0.018 | -0.057 |
| ENSG00000173641 | *HSPB7* | 2.12E-21 | -1.279 | -0.470 | -0.586 |
| ENSG00000102468 | *HTR2A* | 2.72E-04 | 1.727 | 0.017 | 0.061 |
| ENSG00000169495 | *HTRA4* | 1.14E-02 | 1.113 | 0.013 | 0.061 |
| ENSG00000090339 | *ICAM1* | 3.33E-28 | 1.765 | 0.528 | 0.782 |
| ENSG00000160223 | *ICOSLG* | 5.86E-07 | 1.466 | 0.669 | 0.667 |
| ENSG00000131203 | *IDO1* | 3.66E-03 | 2.037 | 0.014 | 0.039 |
| ENSG00000068079 | *IFI35* | 3.00E-04 | 1.172 | 0.427 | 0.544 |
| ENSG00000103742 | *IGDCC4* | 1.76E-07 | -1.160 | -0.255 | -0.449 |
| ENSG00000146678 | *IGFBP1* | 7.61E-10 | 1.294 | 0.405 | 0.741 |
| ENSG00000134470 | *IL15RA* | 2.63E-10 | 1.622 | 0.674 | 0.573 |
| ENSG00000115604 | *IL18R1* | 9.66E-10 | 1.346 | 0.616 | 0.799 |
| ENSG00000115602 | *IL1RL1* | 3.15E-08 | 1.087 | 0.039 | 0.597 |
| ENSG00000136689 | *IL1RN* | 1.69E-03 | 1.066 | 0.011 | 0.016 |
| ENSG00000142677 | *IL22RA1* | 2.27E-02 | -1.101 | -0.017 | -0.023 |
| ENSG00000008517 | *IL32* | 3.73E-13 | 1.719 | 0.582 | 1.167 |
| ENSG00000137033 | *IL33* | 6.84E-20 | 1.898 | 0.719 | 1.047 |
| ENSG00000136696 | *IL36B* | 1.92E-08 | 1.541 | 0.053 | 1.105 |
| ENSG00000136695 | *IL36RN* | 1.66E-04 | 1.555 | 0.015 | 0.067 |
| ENSG00000104951 | *IL4I1* | 2.17E-04 | 1.627 | 0.014 | 1.354 |
| ENSG00000136244 | *IL6* | 6.10E-23 | 5.532 | 2.586 | 3.071 |
| ENSG00000168685 | *IL7R* | 4.60E-04 | 1.031 | 0.049 | 0.440 |
| ENSG00000125347 | *IRF1* | 2.57E-22 | 1.183 | 0.537 | 0.655 |

| | | | | | |
|---|---|---|---|---|---|
| ENSG00000137265 | *IRF4* | 2.31E-03 | 1.058 | 0.018 | 0.048 |
| ENSG00000128604 | *IRF5* | 5.34E-03 | 1.522 | 0.019 | 0.042 |
| ENSG00000091409 | *ITGA6* | 1.52E-12 | -1.016 | -0.498 | -0.758 |
| ENSG00000105855 | *ITGB8* | 1.79E-10 | 1.019 | 0.399 | 0.476 |
| ENSG00000105639 | *JAK3* | 3.98E-07 | 1.728 | 0.758 | 0.817 |
| ENSG00000158445 | *KCNB1* | 1.02E-04 | -1.076 | -0.020 | -0.023 |
| ENSG00000143473 | *KCNH1* | 8.53E-04 | 1.229 | 0.025 | 0.635 |
| ENSG00000184156 | *KCNQ3* | 2.56E-06 | 1.361 | 0.589 | 0.839 |
| ENSG00000157404 | *KIT* | 1.25E-05 | -1.175 | -0.031 | -0.502 |
| ENSG00000239474 | *KLHL41* | 8.44E-18 | -1.099 | -0.474 | -0.639 |
| ENSG00000170454 | *KRT75* | 9.33E-06 | 1.836 | 0.025 | 0.846 |
| ENSG00000115919 | *KYNU* | 3.00E-17 | 1.320 | 0.632 | 0.813 |
| ENSG00000179630 | *LACC1* | 6.12E-09 | 1.035 | 0.342 | 0.523 |
| ENSG00000129988 | *LBP* | 5.74E-10 | 4.020 | 0.014 | 1.532 |
| ENSG00000169744 | *LDB2* | 2.01E-22 | -1.345 | -0.330 | -0.280 |
| ENSG00000174697 | *LEP* | 5.90E-08 | 3.769 | 0.013 | 1.784 |
| ENSG00000128342 | *LIF* | 3.24E-19 | 1.799 | 0.512 | 0.661 |
| ENSG00000236882 | *LINC01554* | 7.50E-03 | 2.247 | 0.011 | 0.036 |
| ENSG00000178734 | *LMO7DN* | 1.99E-04 | -1.012 | -0.789 | -0.667 |
| ENSG00000163431 | *LMOD1* | 4.54E-26 | -2.020 | -0.658 | -0.847 |
| ENSG00000111684 | *LPCAT3* | 4.25E-29 | 1.076 | 0.548 | 0.745 |
| ENSG00000165379 | *LRFN5* | 5.92E-13 | -1.251 | -0.066 | -0.588 |
| ENSG00000173114 | *LRRN3* | 1.60E-12 | -1.017 | -0.627 | -0.700 |
| ENSG00000162951 | *LRRTM1* | 3.74E-02 | 1.511 | 0.012 | 0.035 |
| ENSG00000172901 | *LVRN* | 2.28E-05 | 1.187 | 0.023 | 0.132 |
| ENSG00000165072 | *MAMDC2* | 1.92E-05 | -1.019 | -0.032 | -0.330 |
| ENSG00000107968 | *MAP3K8* | 2.92E-45 | 1.692 | 1.346 | 1.484 |
| ENSG00000081189 | *MEF2C* | 1.32E-09 | -1.091 | -0.308 | -0.611 |
| ENSG00000005102 | *MEOX1* | 3.34E-03 | 1.052 | -0.004 | 0.005 |
| ENSG00000106511 | *MEOX2* | 2.90E-07 | -1.495 | -0.031 | -0.180 |
| ENSG00000106484 | *MEST* | 5.39E-15 | -1.346 | -0.503 | -0.673 |
| ENSG00000185432 | *METTL7A* | 2.94E-13 | -1.075 | -0.401 | -0.611 |
| ENSG00000168389 | *MFSD2A* | 2.74E-04 | 1.450 | 0.015 | 0.069 |
| ENSG00000111341 | *MGP* | 3.90E-13 | -1.158 | -0.466 | -0.652 |
| ENSG00000108960 | *MMD* | 2.26E-21 | 1.029 | 0.409 | 0.588 |
| ENSG00000196611 | *MMP1* | 6.39E-18 | 3.020 | 0.876 | 1.393 |
| ENSG00000166670 | *MMP10* | 4.41E-06 | 1.390 | 0.020 | 0.753 |
| ENSG00000149968 | *MMP3* | 3.27E-14 | 2.020 | 0.051 | 0.916 |
| ENSG00000082126 | *MPP4* | 2.46E-02 | -1.057 | -0.022 | -0.068 |
| ENSG00000072952 | *MRVI1* | 6.13E-14 | -1.613 | -0.045 | -0.159 |

| | | | | | |
|---|---|---|---|---|---|
| ENSG00000178860 | *MSC* | 1.72E-14 | 1.255 | 0.255 | 0.436 |
| ENSG00000138379 | *MSTN* | 1.45E-09 | -2.534 | -0.045 | -0.687 |
| ENSG00000120149 | *MSX2* | 2.74E-09 | 1.079 | 0.090 | 0.580 |
| ENSG00000169715 | *MT1E* | 1.37E-18 | 1.151 | 0.438 | 0.631 |
| ENSG00000198417 | *MT1F* | 1.73E-02 | 1.132 | 0.007 | 0.031 |
| ENSG00000125144 | *MT1G* | 3.17E-06 | 2.363 | 0.018 | 0.047 |
| ENSG00000187193 | *MT1X* | 9.02E-21 | 1.646 | 0.534 | 0.946 |
| ENSG00000125148 | *MT2A* | 9.47E-13 | 1.399 | 0.390 | 0.705 |
| ENSG00000183486 | *MX2* | 6.01E-03 | 2.070 | 1.135 | 0.034 |
| ENSG00000136449 | *MYCBPAP* | 4.20E-02 | -1.452 | -0.004 | -0.009 |
| ENSG00000034971 | *MYOC* | 1.07E-20 | -1.110 | -0.592 | -0.780 |
| ENSG00000101605 | *MYOM1* | 3.38E-05 | -1.389 | -0.036 | -1.203 |
| ENSG00000105835 | *NAMPT* | 2.76E-13 | 2.208 | 0.039 | 0.758 |
| ENSG00000176771 | *NCKAP5* | 1.34E-02 | -1.281 | -0.012 | -0.030 |
| ENSG00000124479 | *NDP* | 1.92E-15 | 2.278 | 0.850 | 1.032 |
| ENSG00000167604 | *NFKBID* | 3.08E-08 | 1.137 | 0.793 | 1.091 |
| ENSG00000144802 | *NFKBIZ* | 2.09E-163 | 3.681 | 2.950 | 3.025 |
| ENSG00000204131 | *NHSL2* | 1.19E-03 | -1.201 | -0.020 | -0.110 |
| ENSG00000166741 | *NNMT* | 1.46E-29 | 2.315 | 1.196 | 1.540 |
| ENSG00000183691 | *NOG* | 2.85E-05 | 1.411 | 0.037 | 0.924 |
| ENSG00000171246 | *NPTX1* | 1.16E-11 | -1.203 | -0.437 | -0.680 |
| ENSG00000182575 | *NXPH3* | 2.54E-04 | -1.157 | -0.006 | -0.018 |
| ENSG00000177989 | *ODF3B* | 3.64E-08 | 1.190 | 0.042 | 0.687 |
| ENSG00000125510 | *OPRL1* | 3.71E-02 | -1.259 | -0.007 | -0.041 |
| ENSG00000145623 | *OSMR* | 2.03E-19 | 1.335 | 0.513 | 0.710 |
| ENSG00000187950 | *OVCH1* | 9.50E-03 | 1.398 | 0.018 | 0.035 |
| ENSG00000243444 | *PALM2* | 7.00E-07 | 1.195 | 0.059 | 0.614 |
| ENSG00000138496 | *PARP9* | 5.02E-05 | 1.118 | 0.752 | 0.673 |
| ENSG00000175426 | *PCSK1* | 5.06E-03 | 1.197 | 0.014 | 0.047 |
| ENSG00000115252 | *PDE1A* | 1.13E-16 | -1.013 | -0.313 | -0.530 |
| ENSG00000113448 | *PDE4D* | 1.97E-07 | 2.047 | 0.024 | 0.907 |
| ENSG00000138735 | *PDE5A* | 1.28E-26 | -1.500 | -0.452 | -0.633 |
| ENSG00000162493 | *PDPN* | 1.10E-32 | 1.809 | 0.745 | 1.046 |
| ENSG00000109272 | *PF4V1* | 2.61E-02 | 1.309 | 0.001 | 0.031 |
| ENSG00000114268 | *PFKFB4* | 1.61E-04 | 1.331 | 0.014 | 0.057 |
| ENSG00000119630 | *PGF* | 3.04E-15 | 1.985 | 0.034 | 1.013 |
| ENSG00000153823 | *PID1* | 3.84E-08 | 1.598 | 0.034 | 0.528 |
| ENSG00000122861 | *PLAU* | 2.46E-04 | 1.060 | 0.045 | 0.748 |
| ENSG00000115896 | *PLCL1* | 1.06E-13 | -1.576 | -0.846 | -0.974 |
| ENSG00000075651 | *PLD1* | 3.55E-07 | 1.400 | 0.025 | 0.406 |

| | | | | | |
|---|---|---|---|---|---|
| ENSG00000147872 | *PLIN2* | 3.26E-10 | 1.020 | 0.064 | 0.356 |
| ENSG00000188313 | *PLSCR1* | 4.31E-08 | 1.150 | 0.754 | 0.843 |
| ENSG00000221866 | *PLXNA4* | 1.04E-03 | 1.046 | 0.013 | 0.099 |
| ENSG00000141682 | *PMAIP1* | 8.72E-09 | 1.447 | 0.065 | 0.728 |
| ENSG00000028277 | *POU2F2* | 6.86E-36 | 2.188 | 1.344 | 1.434 |
| ENSG00000158528 | *PPP1R9A* | 1.64E-03 | -1.531 | -0.017 | -0.045 |
| ENSG00000156475 | *PPP2R2B* | 4.73E-05 | -1.987 | -0.028 | -0.973 |
| ENSG00000061455 | *PRDM6* | 1.51E-03 | -1.144 | -0.022 | -0.049 |
| ENSG00000186652 | *PRG2* | 2.76E-06 | 1.052 | 0.052 | 0.519 |
| ENSG00000146250 | *PRSS35* | 8.58E-12 | -1.544 | -0.732 | -0.766 |
| ENSG00000140368 | *PSTPIP1* | 2.99E-02 | 0.150 | 0.014 | 1.586 |
| ENSG00000117425 | *PTCH2* | 2.14E-03 | -1.028 | -0.022 | -0.119 |
| ENSG00000148344 | *PTGES* | 2.40E-13 | 1.121 | 0.075 | 0.366 |
| ENSG00000122420 | *PTGFR* | 6.82E-26 | 1.698 | 0.727 | 0.921 |
| ENSG00000073756 | *PTGS2* | 4.12E-09 | 1.958 | 0.036 | 0.659 |
| ENSG00000163661 | *PTX3* | 7.13E-07 | 1.178 | 0.563 | 0.854 |
| ENSG00000041353 | *RAB27B* | 1.26E-03 | 1.077 | 0.020 | 0.088 |
| ENSG00000108551 | *RASD1* | 3.87E-06 | 1.895 | 0.018 | 0.645 |
| ENSG00000198774 | *RASSF9* | 8.29E-09 | -1.655 | -0.727 | -1.054 |
| ENSG00000088320 | *REM1* | 2.15E-06 | -1.622 | 0.000 | -0.078 |
| ENSG00000117152 | *RGS4* | 1.50E-06 | -2.045 | -0.678 | -1.080 |
| ENSG00000108370 | *RGS9* | 2.26E-06 | -1.121 | -0.389 | -0.535 |
| ENSG00000143878 | *RHOB* | 1.75E-07 | 1.491 | 0.025 | 0.503 |
| ENSG00000072422 | *RHOBTB1* | 5.90E-10 | -1.219 | -0.297 | -0.273 |
| ENSG00000126785 | *RHOJ* | 2.09E-06 | -1.163 | -0.039 | -0.076 |
| ENSG00000079841 | *RIMS1* | 4.57E-04 | -1.113 | -0.032 | -0.486 |
| ENSG00000042062 | *RIPOR3* | 2.99E-14 | 1.877 | 0.466 | 0.738 |
| ENSG00000169071 | *ROR2* | 2.80E-05 | 1.070 | 0.020 | 0.077 |
| ENSG00000134321 | *RSAD2* | 1.48E-04 | 2.810 | 1.313 | 0.043 |
| ENSG00000186907 | *RTN4RL2* | 1.03E-22 | 1.886 | 0.670 | 0.897 |
| ENSG00000102445 | *RUBCNL* | 7.49E-23 | 1.805 | 0.861 | 1.156 |
| ENSG00000231274 | *SBK3* | 3.88E-05 | 1.810 | 0.020 | 1.017 |
| ENSG00000064932 | *SBNO2* | 2.54E-24 | 1.187 | 0.443 | 0.642 |
| ENSG00000170381 | *SEMA3E* | 9.66E-15 | -1.005 | -0.325 | -0.501 |
| ENSG00000197632 | *SERPINB2* | 1.26E-26 | 2.110 | 0.628 | 1.076 |
| ENSG00000102683 | *SGCG* | 3.28E-04 | -1.620 | -0.018 | -0.545 |
| ENSG00000164690 | *SHH* | 1.80E-06 | 1.846 | 0.028 | 1.085 |
| ENSG00000142178 | *SIK1* | 1.11E-05 | 1.118 | 0.020 | 0.246 |
| ENSG00000141469 | *SLC14A1* | 9.79E-05 | -1.217 | -0.021 | -0.072 |
| ENSG00000110446 | *SLC15A3* | 1.02E-03 | 1.072 | 0.026 | 0.134 |

| | | | | | |
|---|---|---|---|---|---|
| ENSG00000118596 | *SLC16A7* | 1.80E-15 | 1.335 | 0.367 | 0.267 |
| ENSG00000135917 | *SLC19A3* | 4.07E-17 | 2.331 | 0.577 | 0.831 |
| ENSG00000137266 | *SLC22A23* | 1.69E-14 | 1.005 | 0.350 | 0.472 |
| ENSG00000185052 | *SLC24A3* | 3.17E-08 | 1.601 | 0.043 | 0.709 |
| ENSG00000140284 | *SLC27A2* | 3.29E-02 | 1.586 | 0.007 | 0.039 |
| ENSG00000142583 | *SLC2A5* | 7.20E-24 | 1.277 | 0.506 | 0.798 |
| ENSG00000169507 | *SLC38A11* | 2.79E-02 | -1.491 | -0.006 | -0.028 |
| ENSG00000104635 | *SLC39A14* | 9.59E-05 | 1.293 | 0.021 | 0.076 |
| ENSG00000138821 | *SLC39A8* | 2.09E-06 | 2.061 | 0.014 | 0.058 |
| ENSG00000138449 | *SLC40A1* | 4.85E-15 | -1.246 | -0.520 | -0.797 |
| ENSG00000167703 | *SLC43A2* | 4.18E-14 | 1.934 | 0.659 | 0.931 |
| ENSG00000134802 | *SLC43A3* | 1.73E-17 | 1.011 | 0.307 | 0.486 |
| ENSG00000003989 | *SLC7A2* | 3.90E-18 | 2.875 | 1.107 | 1.566 |
| ENSG00000155465 | *SLC7A7* | 6.15E-06 | 1.080 | 0.602 | 0.574 |
| ENSG00000184557 | *SOCS3* | 1.48E-19 | 1.343 | 0.322 | 0.631 |
| ENSG00000112096 | *SOD2* | 6.72E-07 | 1.756 | 0.030 | 0.595 |
| ENSG00000112096 | *SOD2-OT1* | 6.72E-07 | 1.756 | 0.030 | 0.595 |
| ENSG00000137767 | *SQOR* | 3.26E-19 | 1.068 | 0.536 | 0.701 |
| ENSG00000122862 | *SRGN* | 3.45E-17 | 1.112 | 0.055 | 0.352 |
| ENSG00000115415 | *STAT1* | 3.46E-12 | 1.004 | 0.628 | 0.739 |
| ENSG00000138378 | *STAT4* | 3.57E-13 | 1.978 | 0.967 | 1.474 |
| ENSG00000159167 | *STC1* | 1.14E-22 | 2.934 | 1.167 | 1.536 |
| ENSG00000164647 | *STEAP1* | 4.95E-24 | 1.721 | 0.754 | 0.971 |
| ENSG00000157214 | *STEAP2* | 1.74E-09 | 1.548 | 0.591 | 0.731 |
| ENSG00000127954 | *STEAP4* | 1.62E-04 | 1.337 | 0.583 | 0.965 |
| ENSG00000140022 | *STON2* | 1.76E-04 | 1.055 | 0.396 | 0.331 |
| ENSG00000128578 | *STRIP2* | 5.19E-16 | 1.702 | 0.558 | 0.844 |
| ENSG00000170921 | *TANC2* | 2.21E-28 | -1.005 | -0.482 | -0.673 |
| ENSG00000170379 | *TCAF2* | 2.47E-03 | 1.055 | 0.005 | 0.021 |
| ENSG00000176907 | *TCIM* | 7.51E-03 | 1.850 | 0.008 | 0.022 |
| ENSG00000197905 | *TEAD4* | 1.08E-08 | 1.127 | 0.069 | 0.586 |
| ENSG00000105825 | *TFPI2* | 2.05E-19 | 2.438 | 0.907 | 1.345 |
| ENSG00000038295 | *TLL1* | 6.70E-07 | -1.060 | -0.405 | -0.485 |
| ENSG00000137462 | *TLR2* | 8.72E-09 | 2.258 | 0.010 | 0.050 |
| ENSG00000183160 | *TMEM119* | 2.21E-28 | -1.188 | -0.414 | -0.454 |
| ENSG00000139364 | *TMEM132B* | 3.37E-05 | 1.255 | 0.023 | 0.076 |
| ENSG00000164124 | *TMEM144* | 3.53E-03 | 1.019 | 0.025 | 0.051 |
| ENSG00000249992 | *TMEM158* | 2.35E-08 | 1.493 | 0.446 | 0.865 |
| ENSG00000196932 | *TMEM26* | 5.87E-04 | -1.396 | -0.022 | -0.079 |
| ENSG00000182107 | *TMEM30B* | 5.16E-19 | -1.018 | -0.332 | -0.558 |

| | | | | | |
|---|---|---|---|---|---|
| ENSG00000133687 | *TMTC1* | 2.39E-15 | 1.015 | 0.325 | 0.454 |
| ENSG00000185215 | *TNFAIP2* | 8.69E-18 | 1.371 | 0.457 | 0.602 |
| ENSG00000123610 | *TNFAIP6* | 4.04E-12 | 3.429 | 0.822 | 1.478 |
| ENSG00000127863 | *TNFRSF19* | 1.64E-09 | -1.236 | -0.049 | -0.198 |
| ENSG00000028137 | *TNFRSF1B* | 1.84E-14 | 1.273 | 0.460 | 0.680 |
| ENSG00000125735 | *TNFSF14* | 3.42E-04 | 2.187 | 0.008 | 0.030 |
| ENSG00000111907 | *TPD52L1* | 4.27E-17 | -1.066 | -0.367 | -0.469 |
| ENSG00000124731 | *TREM1* | 3.10E-06 | 2.256 | 0.013 | 0.053 |
| ENSG00000173334 | *TRIB1* | 2.06E-08 | 1.277 | 0.036 | 0.554 |
| ENSG00000158022 | *TRIM63* | 1.98E-02 | -1.277 | -0.020 | -0.047 |
| ENSG00000104321 | *TRPA1* | 1.66E-03 | 1.046 | 0.020 | 0.083 |
| ENSG00000182463 | *TSHZ2* | 5.00E-10 | 1.054 | 0.329 | 0.499 |
| ENSG00000025708 | *TYMP* | 7.04E-39 | 2.023 | 1.057 | 1.337 |
| ENSG00000162692 | *VCAM1* | 8.74E-07 | 1.198 | 0.035 | 0.191 |
| ENSG00000170162 | *VGLL2* | 6.70E-03 | 1.047 | 0.013 | 0.084 |
| ENSG00000119703 | *ZC2HC1C* | 6.46E-03 | 1.671 | 0.014 | 0.032 |
| ENSG00000163874 | *ZC3H12A* | 2.17E-47 | 3.152 | 2.226 | 2.377 |
| ENSG00000144331 | *ZNF385B* | 1.52E-10 | -1.205 | -0.315 | -0.284 |

**Supplementary Table 6.** Results from GSEA Hallmark pathway analysis. (A) Overview of hallmark pathways significantly (padj<0.05) changed by IL-17 treatment, with the normalised enrichment score (NES) in **black** for an increase in NES and **red** for decreases in NES.

| **Pathway** | **Chondrocytes** | | | **Synovial fibroblasts** | | |
|---|---|---|---|---|---|---|
| | IL-17A | IL-17AF | IL-17F | IL-17A | IL-17AF | IL-17F |
| Allograft Rejection | 2.43 | 2.10 | 1.83 | 2.35 | 1.86 | 1.98 |
| Angiogenesis | 1.98 | 2.09 | 2.00 | 1.75 | | 1.80 |
| Apical Junction | | | -1.55 | | | |
| Apoptosis | 2.04 | 1.80 | | | | |
| Bile Acid Metabolism | | | | 1.78 | | |
| Coagulation | 2.03 | 2.05 | 1.74 | 1.87 | 1.70 | |
| Complement | 2.46 | 2.30 | 1.98 | 2.24 | 2.14 | 2.06 |
| Epithelial Mesenchymal Transition | 2.31 | 2.19 | 1.79 | 1.77 | 1.78 | 1.82 |
| Estrogen Response Early | 1.71 | 1.75 | | | | |
| Glycolysis | 2.28 | 2.06 | 1.75 | 1.66 | | |
| Heme Metabolism | | 1.55 | | | | |
| Hypoxia | 2.71 | 2.59 | 1.92 | 2.05 | 1.63 | |
| Il2 Stat5 Signaling | 2.36 | 1.90 | 1.76 | 1.69 | | 1.64 |
| Il6 Jak Stat3 Signaling | 2.59 | 2.54 | 2.10 | 2.42 | 2.20 | 2.09 |
| Inflammatory Response | 3.20 | 2.88 | 2.28 | 2.88 | 2.37 | 2.26 |
| Interferon Alpha Response | 2.28 | 1.87 | | 2.04 | 1.99 | 1.97 |
| Interferon Gamma Response | 2.85 | 2.61 | 2.24 | 2.73 | 2.37 | 2.25 |
| Kras Signaling Dn | | | -1.65 | | | |
| Kras Signaling Up | 2.34 | 1.99 | 1.74 | 2.28 | 1.80 | |
| Mitotic Spindle | | | | | -1.64 | |
| Mtorc1 Signaling | 2.09 | 2.03 | 1.53 | 1.78 | | |
| Myc Targets V1 | 1.56 | 2.02 | 1.81 | 1.70 | | |
| Myc Targets V2 | | | | 1.75 | | |
| Myogenesis | | | -1.61 | -1.97 | -1.82 | -1.87 |
| Oxidative Phosphorylation | | 1.59 | | | | |
| Reactive Oxygen Species Pathway | | 1.70 | 1.77 | | | |
| Tnfa Signaling Via Nfkb | 3.43 | 3.27 | 2.60 | 2.95 | 2.46 | 2.35 |
| Unfolded Protein Response | 1.98 | 1.93 | | | | |
| Uv Response Up | 2.41 | 2.38 | 1.92 | 1.80 | -1.59 | |
| Xenobiotic Metabolism | 2.07 | 2.09 | 1.70 | | | |
